# Gliomas phenocopy an inborn error of metabolism to drive neuronal activity and tumor growth

**DOI:** 10.1101/2025.09.15.676412

**Authors:** Kalil G. Abdullah, Kenji Miki, Charles K. Edgar, Shuangcheng Alivia Wu, Yi Xiao, Milan R. Savani, Maged T. Ghoche, Shawn E. Kotermanski, Yuan-Tai Huang, Jeffrey I. Traylor, Lei Guo, Kathryn Gunn, William H. Hicks, Diana D. Shi, Min Tang, Michael M. Levitt, Mohamad El-Shami, Skyler Oken, Namya Manoj, Bailey C. Smith, Vinesh T. Puliyappadamba, Pranita Kaphle, Tracey Shipman, Raymond E. West, Chaoying Liang, Toral R. Patel, Kimmo J. Hatanpaa, Prithvi Raj, Shang Ma, Alexander Ksendzovsky, Thomas D. Nolin, Bradley C. Lega, Pascal O. Zinn, Bianca J. Kuhn, Natalie M. Clark, C. Williams, D.R. Mani, Michael A. Gillette, Marco A. Calzado, Lin Xu, Lauren G. Zacharias, Feng Cai, Thomas P. Mathews, Julie-Aurore Losman, Timothy E. Richardson, Benjamin Levi, Michelle Monje, Jay R. Gibson, Osaama H. Khan, Sameer Agnihotri, Kimberly M. Huber, Simon Chamberland, Ralph J. DeBerardinis, Samuel K. McBrayer

## Abstract

The metabolic hallmarks of high-grade glioma (HGG) are not fully understood. Human brain tissue metabolomics revealed that the creatine synthesis pathway intermediate guanidinoacetate (GAA) accumulated ∼100-fold in HGGs relative to controls, which was caused by imbalanced activities of enzymes in this pathway. Glioma cells secreted GAA rather than using it to produce creatine, implicating an unexpected function. GAA accumulates in GAMT deficiency, an inborn error of metabolism, and elevates neuronal excitability. Neuronal excitability is also increased in glioma and drives tumor growth through neuron-glioma interactions. We hypothesized that glioma-generated GAA excites surrounding neurons. Indeed, GAA induced neuronal hyperactivity by activating GABA_A_ receptors and causing depolarizing GABA currents in glioma-associated neurons with dysregulated chloride homeostasis. Depleting tumoral GAA decreased electrochemical activity, neuron-glioma interactions, and tumor aggressiveness. Our findings unveil a new mechanism linking cancer metabolism with cancer neuroscience and leverage human genetics to nominate GAA synthesis as a target in gliomas.

## INTRODUCTION

Gliomas are the most common type of central nervous system malignancy diagnosed in adults. Its most aggressive forms, high grade gliomas (HGG, Grade 4), carry a median life expectancy of only 1-2 years. Therefore, there is an urgent need for insights into molecular mechanisms underlying aggressive glioma behavior and new therapeutic strategies that disrupt these processes. Adult HGGs encompass two diagnostic categories that are defined by isocitrate dehydrogenase (IDH) mutational status^1^: 1) Glioblastoma (GBM), IDH-wildtype, Grade 4, and 2) Astrocytoma, IDH-mutant, Grade 4. IDH mutant enzymes display neomorphic synthesis of the oncometabolite (*R*)-2-hydroxyglutarate [(*R*)-2HG], which reprograms the epigenome^2–9^ to promote glioma development and maintenance^10–12^. This metabolic alteration underpins the antitumor activity of the mutant IDH1/2 inhibitor vorasidenib, which was recently approved by the United States Food and Drug Administration for the treatment of IDH-mutant low-grade glioma^13^. These advances provide proof-of-principle that altered metabolism can serve as both an oncogenic driver and a therapeutic target in glioma. However, in contrast to our understanding of metabolic reprogramming by IDH oncogenes, which represent an uncommon subset of HGG, knowledge of metabolic mechanisms that sustain HGGs across genetic and transcriptomic subtypes is limited.

Prior research has provided vital insights into the biochemical landscape of adult-type gliomas^14–17^. HGGs display reduced creatine content^15, 18, 19^ and harbor elevated levels of nucleotides and their precursors^14, 20, 21^, tryptophan^15^, lysine catabolites^22^, lipid species^14, 17^, and redox metabolites^14^ including glutathione^15^. Nevertheless, comprehensive metabolomic analyses of HGG, LGG, and non-malignant brain specimens have not kept pace with comparable genomics and transcriptomics analyses of these tissues^23–25^. This limited momentum has impeded insights into metabolic alterations that are causally linked to aggressive glioma behavior.

Growth-promoting interactions of gliomas with the neural circuits they invade have emerged as a major axis of pathophysiology. The activity of neurons, especially glutamatergic neurons, robustly drives the growth^26–31^ and invasion^32, 33^ of multiple forms of gliomas through paracrine signaling factors^26^ and bona fide neuron-to-cancer synapses^28, 31^. In turn, gliomas increase the excitability of neurons^34–37^, contributing to glioma-associated seizures and further augmenting glioma growth-promoting, activity-regulated neuronal interactions. The role that metabolic mechanisms may play in this pathophysiology, however, is largely unexplored.

To complement previous studies and enhance the depth of metabolomic characterization of HGGs, we collected a set of 91 primary human brain tissue samples and used them to conduct multi omics analyses including quantification of 563 polar metabolites and lipids. To our knowledge, this dataset represents the most detailed metabolic portrait of HGG constructed to date. These specimens spanned non malignant cortex, brain metastases, lower grade gliomas (LGG, Grades 2–3), and HGGs. We integrated polar metabolomics, lipidomics, and transcriptomics data with clinical and molecular tissue features. Through this effort, we sought to discover biochemical patterns unique to the HGG metabolome and define their contributions to HGG maintenance and progression. We report the discovery of a metabolic phenotype present in HGG that spans genetic and transcriptomic disease subtypes and recapitulates the biology of a rare inborn error of creatine metabolism, GAMT deficiency. We further demonstrate that this metabolic phenotype drives hyperexcitability of the HGG microenvironment and tumor growth, thereby revealing a tractable target for therapeutic intervention in patients with HGG.

## RESULTS

### Metabolomic and lipidomic profiling of human brain and brain tumor tissues

To generate new insights into brain tumor metabolism, we prospectively collected 91 adult human brain tissue samples from tumor or seizure foci resection surgeries. These samples included LGGs, HGGs, brain metastases, and non-malignant brain specimens (Figure S1A). Each tissue was divided and processed for metabolomic, lipidomic, and transcriptomic analyses. We used defined gene expression programs^25^ to classify glioma samples as Proneural, Classical, or Mesenchymal (Figure S1B). Next, we integrated data for 563 polar metabolites and lipid species with clinical and molecular information associated with each specimen and performed unsupervised clustering to detect shared biochemical profiles in an unbiased manner (Figure 1A and Table S1). This approach revealed seven dominant clusters, including subgroups enriched in Mesenchymal GBMs, recurrent LGGs, index LGGs, brain metastases, and non-malignant brain samples, as well as two mixed populations that did not display clearly defined correlates with molecular or clinical features. Importantly, supervised dimensionality reduction analysis demonstrated that HGGs, LGGs, brain metastases, and non-malignant brain samples were readily distinguishable by biochemical profiles (Figure 1B). Moreover, glioma samples of different histological subtypes, grades, and transcriptional subtypes could be discriminated based on metabolite content (Figures S1C-E).

**Figure 1.**
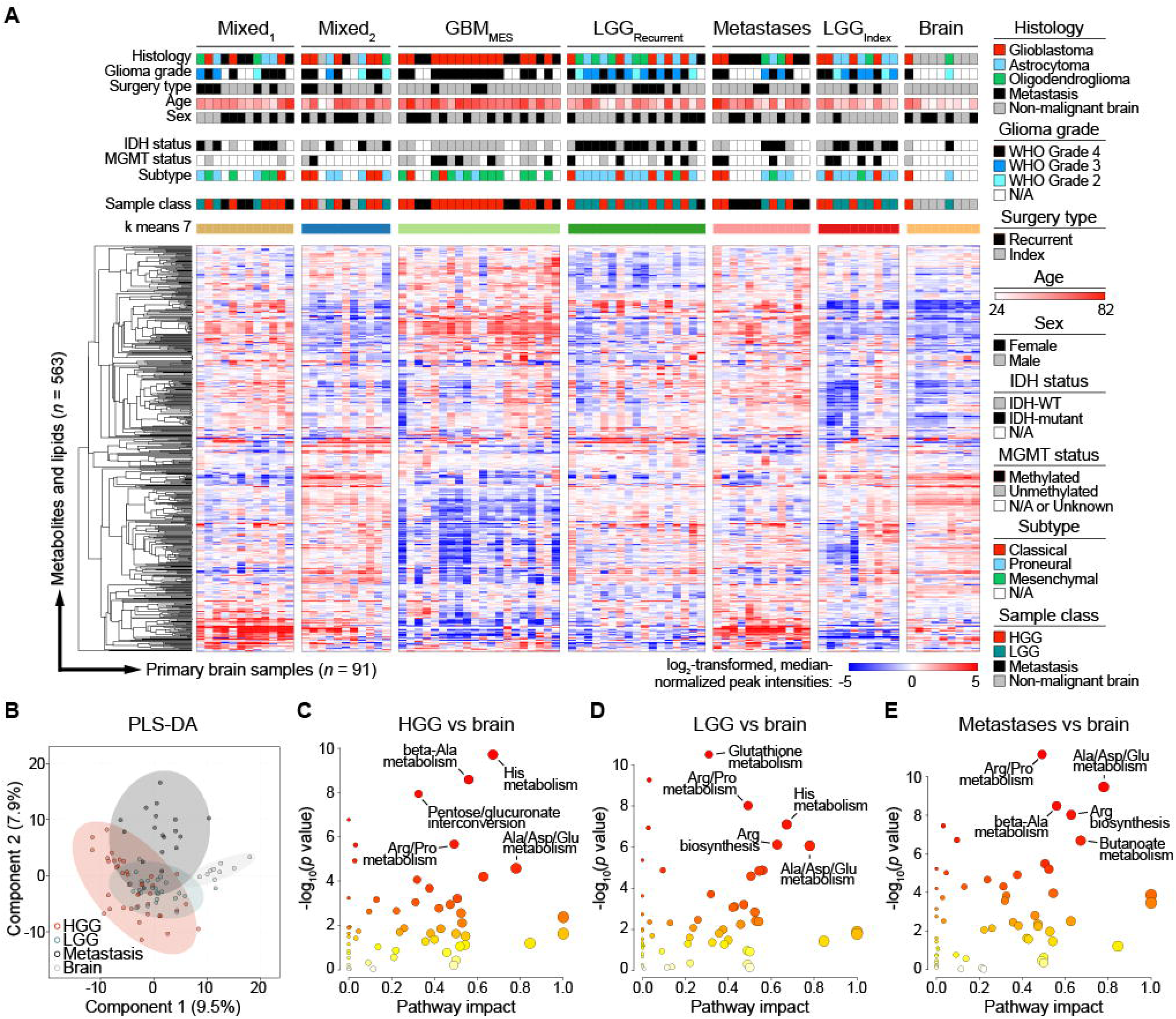
Metabolomic stratification of human brain tissues. **(A)** Profiling of polar metabolites and lipids (*n=*563) in tissue specimens from adult HGG (*n*=39), LGG (*n*=25), brain metastases (*n*=18), and non-malignant brain (“Brain”; *n*=9). Metabolite peak intensities were log_2_-transformed and median normalized. Data are stratified by *k*-means clustering with *k*=7. **(B)** Partial least-squares discriminant analysis (PLS-DA) of metabolite content of patient tissue samples grouped by sample class. **(C-E)** Metabolite set enrichment analyses in (C) HGG, (D) LGG, or (E) metastases relative to non-malignant brain tissue. See also Figure S1.

Before evaluating novel biochemical hallmarks of brain tumor classes, we first assessed expected metabolic differences among glioma and non-malignant brain samples. 2HG levels were markedly elevated in IDH-mutant glioma specimens relative to IDH-wildtype gliomas and non-malignant brain controls (Figure S1F). L-DOPA, a precursor to dopamine, was enriched in non-malignant brain samples compared with HGGs (Figure S1G), consistent with prior work^38^. Additionally, intermediates in lysine degradation^22^ (aminoadipate) and nucleotide metabolism^20, 21^ (orotate and deoxyguanosine) pathways were elevated while the ratio of N-acetylaspartate to creatine (an imaging biomarker of HGG^18, 19^) decreased in HGG versus non-malignant brain tissues (Figures S1H-K). Observing these anticipated biochemical patterns affirmed the validity of our dataset.

Next, we conducted pathway analyses to identify biochemical processes that are altered in specific brain tumor subtypes. Changes in metabolism of amino acids, including arginine, proline, glutamate, histidine, alanine, and aspartate, were observed independent of tumor subtype in all cancer versus non-malignant brain tissue comparisons (Figures 1C-1E). HGGs showed specific changes in in the pentose-glucuronate interconversion pathway (Figure 1C), which were driven by enrichment of levels of hexose monophosphates and glucuronic acid and depletion of pentitols relative to non-malignant brain. Comparing non-malignant brain tissues with LGGs, which are defined by the presence of *IDH1* or *IDH2* mutations^1^, glutathione metabolism ranked among the top differentially regulated pathways (Figure 1D). This finding aligns with our prior study showing that 2HG inhibits branched chain amino acid transaminase-dependent synthesis of glutamate^39^, a precursor to glutathione, as well as others’ work describing an inverse correlation between 2HG and glutathione abundance in human gliomas^40, 41^. Finally, butanoate metabolism differed markedly between brain metastases and non-malignant brain samples (Figure 1E), principally reflecting lower levels of the neurotransmitter GABA in the former. This alteration may be attributable to lower neuronal content of metastases relative to primary brain tumors given that metastases tend not to display the same degree of diffuse infiltration and integration into the brain that is characteristic of gliomas.

### Guanidinoacetate accumulates in HGG and is associated with a bottleneck in the creatine synthesis pathway

Accumulation of the oncometabolite 2HG is a unique and nearly universal metabolic hallmark of IDH-mutant LGGs, as well as the HGGs that develop from them^42^. This discovery has opened new avenues for diagnosing^43, 44^ and monitoring^45^ IDH-mutant gliomas and prompted approval of the mutant IDH1/2 inhibitor vorasidenib to treat these tumors by the U.S. Food and Drug Administration. Considering these transformative effects on the neuro-oncology field, we questioned whether similar patterns of recurrent metabolite accumulation may also be present in predominantly IDH-wildtype HGGs. We rank-ordered metabolites according to the degree of enrichment or depletion they display in HGG specimens compared to non-malignant brain tissues (Figure 2A). We found that an intermediate in the creatine synthesis pathway, guanidinoacetate (GAA), markedly accumulated in HGGs (Figures 2B and 2C). Upon extending our analysis to all metabolites in this pathway, incorporating data from LGG samples, and focusing on tumor samples from index surgical cases, we found that preferential upregulation of GAA in HGG was not associated with a global increase in creatine or its products. GAA levels were elevated ∼100-fold in HGG versus non-malignant brain (Figure 2C). In contrast, creatine and creatinine were decreased while phosphocreatine showed a similar, albeit statistically insignificant, trend (Figures 2D-F). Interestingly, the effect size of GAA accumulation in HGG was similar to 2HG enrichment in IDH-mutant gliomas when compared with relevant controls (Figures 2C and S1F).

**Figure 2.**
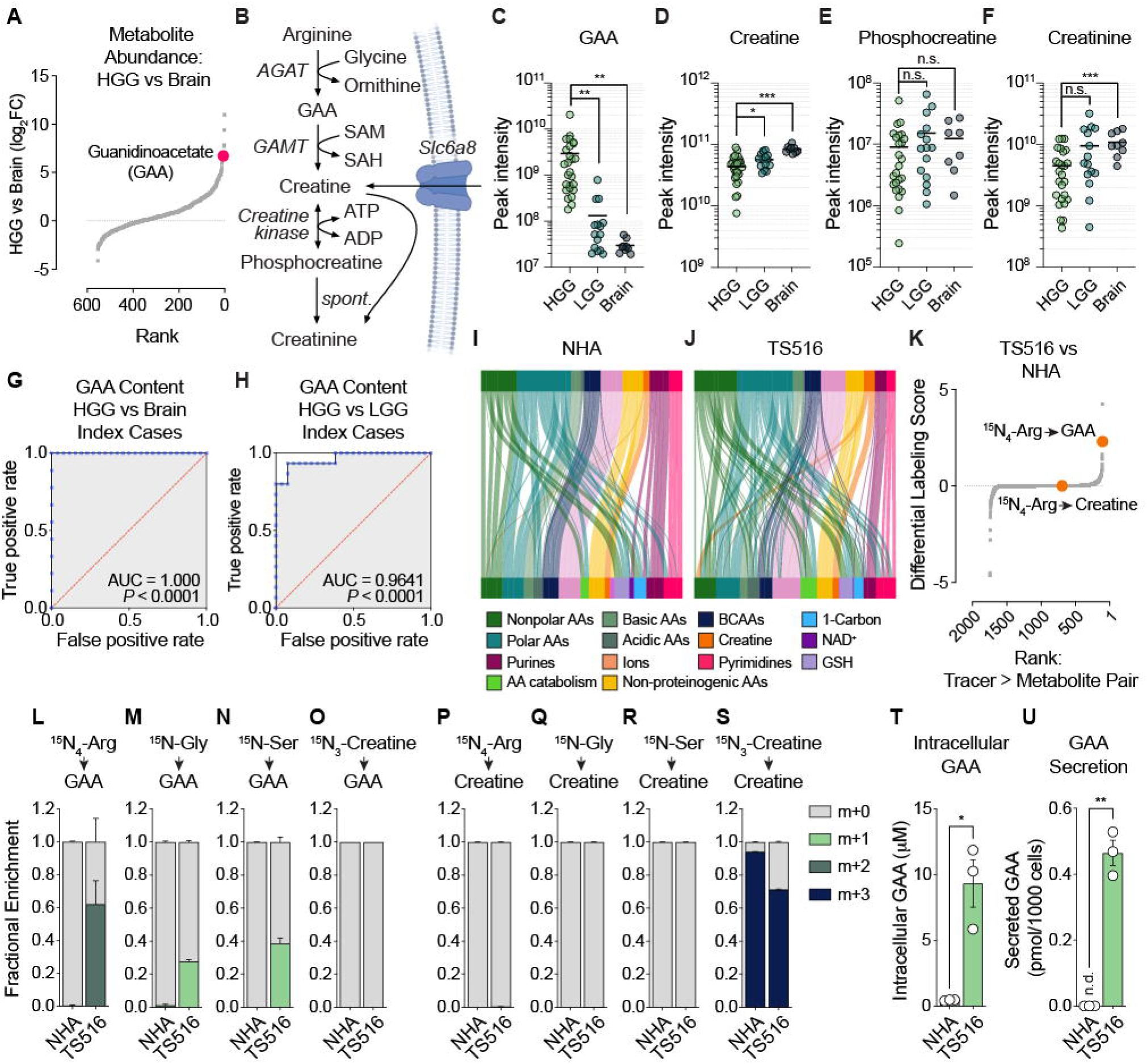
Identification of guanidinoacetate accumulation as a metabolic hallmark of high-grade glioma. **(A)** Waterfall plot of relative metabolite abundance in HGG vs non-malignant brain samples. **(B)** Schema depicting the creatine synthesis pathway. **(C-F)** Peak intensities of (C) guanidinoacetate (GAA), (D) creatine, (E) phosphocreatine, and (F) creatinine in index samples of HGG (*n=*26), LGG (*n=*15), and non-malignant (“Brain”, *n=*9). **(G-H)** ROC analysis of GAA content as a discriminant marker of index HGG tissue (*n*=26) compared to (G) non-malignant brain tissue (*n*=9) or (H) index LGG tissue (*n*=15). **(I-J)** Sankey diagrams depicting labeling of ^15^N-labeled tracers (top bars) in nitrogen metabolism pathway intermediates in immortalized astrocytes (NHA Donor #1) (I) or TS516 GSCs (J). **(K)** Waterfall plot of all tracer to metabolite labeling interations ranked by differential labeling score. **(L-S)** Fractional enrichment of label from indicated tracer in (L-O) GAA or (P-S) creatine following incubation of NHA Donor #1 or TS516 cells with ^15^N-labeled tracer for 18 hours (*n*=3 per cell line). **(T-U)** Intracellular (T) and secreted (U) GAA content of NHA Donor #1 and TS516 cultures at 48 hours in HPLM (*n=*3 per cell line). Data are means ± SEM. n.s.= not significant, n.d. = not detected, *p<0.05, **p<0.01, ***p<0.001 (Welch’s *t*-test). See also Figure S2.

We next evaluated patterns of GAA accumulation across glioma subgroups. GAA, but not other creatine synthesis pathway intermediates, was numerically but not significantly increased in recurrent relative to index LGG samples (Figures S2A-D). These data imply that GAA enrichment is associated with aggressive glioma disease and may manifest during progression of some LGGs in addition to occurring broadly in HGGs. Stratifying HGGs by IDH mutational status and gene expression programs, we found that GAA robustly and selectively accumulates across all subgroups of Grade 4 IDH-mutant astrocytoma and GBM (Figures S2E-H). These effects were unique to glioma because we observed <3-fold enrichment of GAA in lung squamous tumors relative to benign adjacent lung tissues^46^, an effect that was associated with a modest increase in tumoral creatine (Figures S2I-L).

To more directly address the prevalence of GAA upregulation in HGGs versus LGGs and non-malignant brain, we constructed receiver operating characteristic (ROC) curves describing the relationship between GAA content and glioma presence and type in surgical specimens from index and recurrent resections or those from index cases alone. GAA content served as a robust tissue biomarker, effectively discriminating HGG tissue from LGG or non-malignant brain comparators (Figures 2G, 2H, S2M, and S2N). In contrast, GAA levels weakly distinguished LGG and non-malignant brain samples (Figures S2O and S2P). Our data align conceptually with a prior study showing that GAA was elevated in the interstitial fluid of 8 adult-type HGGs^47^. The statistical power provided by our collection of primary tissue samples establishes that GAA accumulation is a highly recurrent metabolic hallmark of GBM tumors and likely of Grade 4 IDH-mutant astrocytomas as well.

Observing simultaneous enrichment of GAA and depletion of creatine in HGG suggested that the creatine synthesis pathway may be reprogrammed in these tumors. To obtain a comprehensive view of substrate-product interactions involving this pathway, we leveraged a platform for nitrogen metabolism profiling recently developed by our group^48^. This platform supports parallel stable isotope tracing of 31 distinct ^15^N-labeled nutrients in Human Plasma-like Medium^49^ (HPLM) in two cell cultures. We used this platform to probe global nitrogen metabolism programs in NHA non-transformed, immortalized astrocytes and TS516 GBM glioma stem-like cells (GSCs) (Figures 2I and 2J, Table S2). In the creatine synthesis pathway, arginine and glycine serve as substrates for the enzyme arginine:glycine amidinotransferase (AGAT), which produces both GAA and ornithine (Figure 2B). GAA is then methylated by the enzyme guanidinoacetate N-methyltransferase (GAMT) to produce creatine in an S-adenosylmethionine (SAM)-dependent manner. Ranking tracer-metabolite interactions based on differential labeling in NHA and TS516 lines showed that ^15^N_4_-arginine-dependent labeling of GAA was strongly enriched in GBM cells but ^15^N_4_-arginine-dependent labeling of creatine was not (Figure 2K). These data indicate that HGGs harbor a bottleneck in the creatine synthesis pathway, displaying robust AGAT-dependent GAA formation that is uncoupled from GAMT-dependent creatine generation.

To further investigate this model and validate nitrogen metabolism profiling results, we performed low-throughput tracing experiments with ^15^N_4_-arginine, ^15^N-glycine, ^15^N-serine (a precursor to glycine), and ^15^N_3_-creatine in NHA and TS516 cultures. Only TS516 cells displayed strong GAA labeling by ^15^N_4_-arginine, ^15^N-glycine, and ^15^N-serine (but not ^15^N_3_-creatine) tracers (Figure 2L-2O). In contrast, neither NHA nor TS516 cells used ^15^N_4_-arginine, ^15^N-glycine, or ^15^N-serine tracers to produce creatine (Figures 2P-2R). Rather, both cell types predominantly consumed extracellular ^15^N_3_-creatine to sustain their intracellular creatine pools (Figure 2S). These data revealed an unexpected pattern of metabolism: TS516 GBM cells robustly synthesize GAA but do not use it to produce creatine. Because GAMT is the only enzyme in human cells known to use GAA as a substrate, we hypothesized that GBM cells may constitutively export newly synthesized GAA. Indeed, we found that TS516 cells readily accumulate and secrete GAA while NHA cells do not (Figures 2T and 2U).

### Creatine synthesis pathway reprogramming and GAA secretion are recurring features of HGG

We next sought to evaluate the generalizability of GAA accumulation and secretion in HGG. We assessed metabolite profiling data from two large independent collections of human brain tissue specimens. The first included HGG, LGG, and non-malignant brain specimens (Figures 3A-3D) while the second comprised only HGG and LGG samples (Figures 3E-3G). Using these validation cohorts, we confirmed preferential upregulation of GAA, but not other creatine synthesis pathway intermediates, in HGG versus LGG or non-malignant brain tissues. Therefore, pervasive GAA accumulation in HGGs cannot be explained by our patient sample collection or metabolite quantification approach.

**Figure 3.**
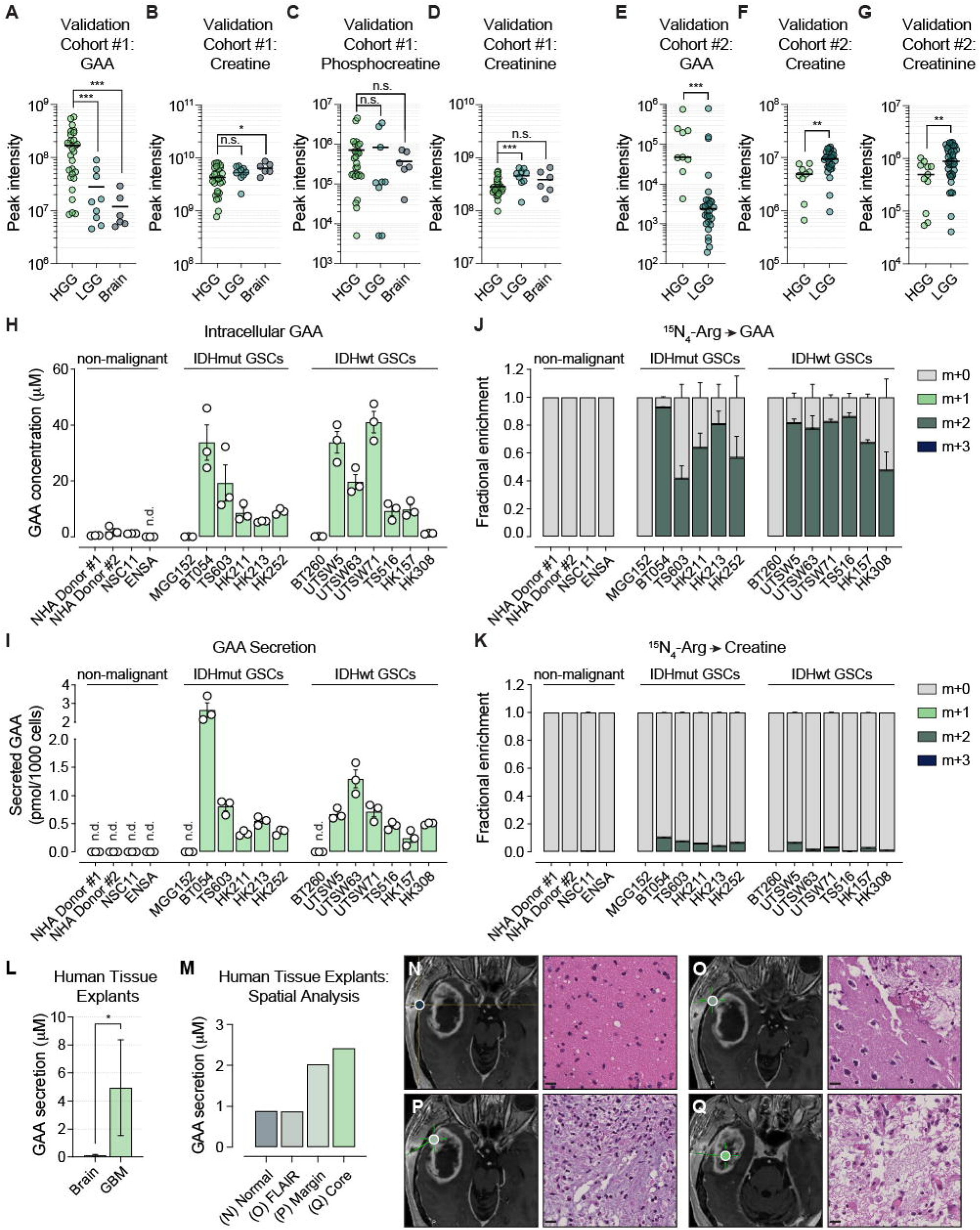
HGG cells display a bottleneck in the creatine synthesis pathway and GAA secretion. (A-D) Peak intensities of (A) GAA, (B) creatine, (C) phosphocreatine, and (D) creatinine in Validation Cohort #1 of HGG (*n=*28), LGG (*n=*9), and non-malignant brain (*n=*6) tissue samples. **(E-G)** Peak intensities of (E) GAA, (F) creatine, and (G) creatinine in Validation Cohort #2 of HGG (*n=*9) and LGG (*n=*28) human surgical specimens. **(H-I)** (H) Intracellular GAA content or (I) GAA secretion in non-malignant cells (*n*=4), IDH-mutant GSCs (*n*=6), and IDH WT GSCs (*n=*7) following 48-hour culture in HPLM (*n=*3 per cell line). **(J-K)** Fractional enrichment of label from ^15^N_4_-arginine in (J) GAA and (K) creatine in the same cell line panel following 18-hour incubation in tracer-containing HPLM (*n*=3 per cell line). (**L**) GAA secretion following 18-hour culture of explanted human largely non-malignant neocortex (brain, *n*=7) or GBM (*n*=5) tissues in HPLM. (**M-Q**) Spatially resolved GAA secretion in human GBM. (M) Explant GAA secretion in samples generated from four distinct points within a single GBM corresponding to (N) largely non-malignant brain (“Normal”), (O) FLAIR-enhancing tumor-adjacent neocortex (“FLAIR”), (P) tumor margin (“Margin”), and (Q) tumor core (“Core”) as assessed on intraoperative magnetic resonance imaging (MRI). Scale bar = 20 µm. Data are means (A-D and H-L) or medians (E-G) ± SEM. n.d. = not detected. n.s.= not significant, *p<0.05, **p<0.01, ***p<0.001 (Welch’s *t*-test in A-D, Mann-Whitney test in E-G, Kolmogorov-Smirnoff test in L). See also Figure S3.

Differential amino acid metabolism activities in TS516 and NHA cells (Figures 2I-2U) suggested that GAA enrichment and creatine depletion in HGGs may be driven by cell-autonomous metabolic alterations in tumor cells. If true, evidence of creatine synthesis pathway reprogramming should be readily observable in diverse patient-derived GSC lines. To address this possibility, we quantified intracellular GAA content, rates of GAA secretion, and ^15^N_4_- arginine-dependent labeling of GAA and creatine in a panel of IDH-mutant and IDH-wildtype GSC lines, NHA non-transformed, immortalized astrocyte lines derived from two donors, and non-transformed neural stem cell (NSC) lines (ENSA and NSC11). IDH-mutant GSC lines were derived from Grade 4 IDH-mutant astrocytomas (MGG152, HK211, HK213 and HK252 cells) or Grade 3 IDH-mutant and 1p/19q-codeleted oligodendrogliomas (BT054 and TS603 cells). All IDH-wildtype GSC lines were derived from GBMs except for BT260 cells, which were generated from a Grade 3 IDH-wildtype oligodendroglioma prior to the World Health Organization’s update to CNS tumor classification criteria in 2021^1^. All non-malignant NHA and NSC lines displayed low GAA cellular content, negligible GAA secretion, and minimal de novo creatine synthesis pathway activity (Figures 3H-3K). In contrast, the vast majority of GSC lines demonstrated clear evidence of creatine synthesis pathway reprogramming, including GAA accumulation and secretion as well as robust labeling of GAA, but not creatine, by ^15^N_4_-arginine. The two exceptions were MGG152 and BT260 cells. Notably, HK308 GBM cells displayed robust GAA synthesis and secretion but not intracellular accumulation. These data suggest that GAA is rapidly exported by these cells upon synthesis. Importantly, differences in ^15^N_4_-arginine metabolism between GSC lines and non-transformed neural cells were not confounded by reduced tracer uptake in the latter (Figure S3).

These findings imply that HGGs display high rates of GAA secretion, but non-malignant brain tissues do not. To test this prediction, we leveraged recent advances in organoid explant modeling of non-malignant brain and brain tumor tissues^50–53^ to evaluate GAA secretion from primary human brain tissues specimens. We created a series of human brain tissue explants from non-malignant brain and GBM samples (Table S3). Next, we cultured these explants in a customized formulation of HPLM^50^ immediately upon surgical resection. Then we collected conditioned media and measured GAA released by each explant. As anticipated, GBM but not non-malignant brain explants displayed robust GAA secretion (Figure 3L). To determine whether GAA release is broadly observed in brain tissue from patients with GBM or if this is a specific feature of the GBM microenvironment, we used stereotactic neurosurgical navigation to produce spatially defined brain tissue explants from a patient who underwent surgical tumor resection of a GBM. We established explants from normal-appearing non-malignant tissue, tumor-adjacent edematous tissue marked by Fluid-Attenuated Inversion Recovery (FLAIR) hyperintensity, and tissues from the tumor margin and tumor core (Figures 3M-Q). Explants with high tumor cell content derived from the GBM margin and core displayed elevated GAA secretion relative to FLAIR-positive and normal-appearing tissues with low tumor cell content. Considering data from human tissue explants together with those from GSC lines, our findings indicate that HGG cells synthesize GAA and release it into tumor interstitial fluid, leading to 10- to 100-fold increases in GAA content in the HGG microenvironment.

### Guanidinoacetate secretion is associated with imbalanced AGAT and GAMT expression

To begin to define the molecular mechanism underlying aberrant GAA secretion by HGG cells, we evaluated expression of creatine synthesis pathway enzymes in human HGGs and non-malignant brain specimens by querying TCGA and GTEx RNA sequencing datasets (Figure 4A). We found that AGAT expression was broadly upregulated while expression of GAMT and distal enzymes in the pathway were downregulated in HGG tissues relative to non-malignant brain. These differences were observed across tumor transcriptional subtypes and IDH genotypes. We hypothesized that an increase in AGAT expression without a commensurate change in GAMT may saturate GAMT catalytic activity and result in GAA accumulation and release from HGG cells. This conceptual model would imply that the ratio of AGAT:GAMT expression may serve as a biomarker of creatine synthesis pathway reprogramming and GAA enrichment in HGG. Indeed, the AGAT:GAMT mRNA expression ratio was markedly elevated in HGGs versus non-malignant human brain tissues (Figure 4B).

**Figure 4.**
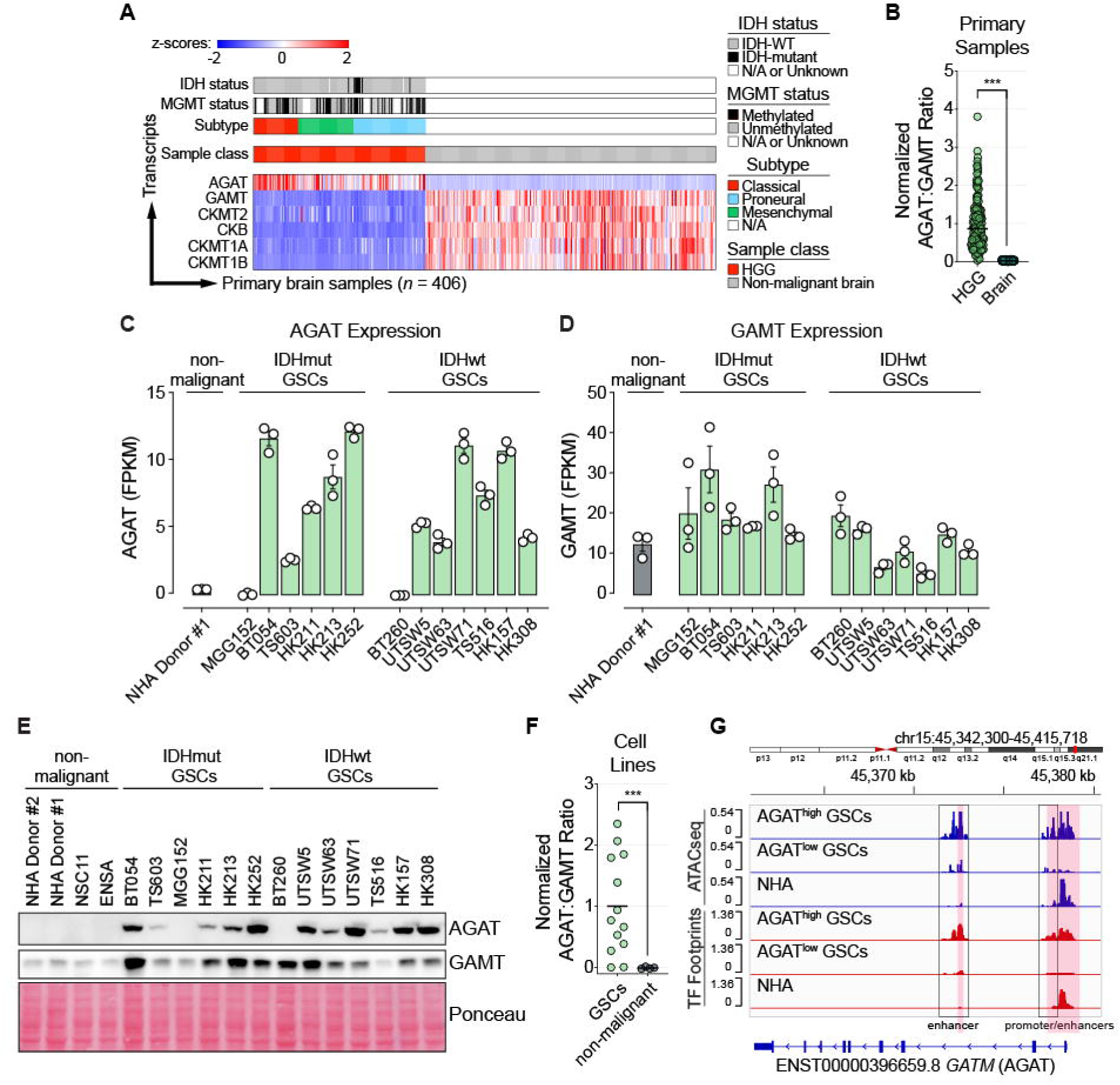
Imbalanced expression of AGAT and GAMT is associated with GAA accumulation in glioma. **(A)** RNAseq analysis of transcript levels of AGAT, GAMT, and creatine kinase isoforms (CKMT2, CKB, CKMT1A/B) in HGG surgical specimens (*n=*151) and non-malignant brain specimens (*n=*255). Data are z-scores of log_2_(transcripts per million +1) values. Data for HGG samples were reanalyzed from the Cancer Genome Atlas (TCGA)^74^. Data for non-malignant brain specimens were reanalyzed from the Genome Tissue Expression Project (GTEX)^75, 76^. **(B)** Relative expression of AGAT and GAMT, quantified as ratio of transcript abundances for a given sample, in HGG and non-malignant brain RNAseq from TCGA and GTEX. **(C-D)** RNAseq analysis of AGAT and GAMT transcript abundance in a panel of IDH mutant GSCs (*n=*6), IDH WT GSCs (*n=*7), and non-malignant (*n=*1) cells (*n=*3 per cell line). Data are fragments per kilobase of transcript per million mapped reads (FPKM). **(E)** Representative immunoblots of AGAT and GAMT expression in a panel of IDH mutant GSCs (*n*=6), IDH WT GSCs (*n*=7), and non-malignant (*n=*4) cells. **(F)** Densitometry analysis of the ratio of AGAT and GAMT protein expression from (E). Data are presented as means. (**G**) Aggregated chromatin accessibility and transcription factor footprint tracks from AGAT^high^ GSCs (*n*=6), AGAT^low^ GSCs (*n*=2), and NHA Donor #1 cells. ***p<0.001 (Welch’s *t*-test). See also Figure S4.

These data provided a global portrait of creatine synthesis pathway gene expression in brain tissues but did not address how AGAT and GAMT expression are regulated within discrete cell populations. Therefore, we assessed AGAT and GAMT transcript abundance in a published single-cell RNA sequencing dataset of human GBM. Tumor cells and oligodendrocytes displayed the highest levels of AGAT expression and AGAT:GAMT ratios (Figures S4A-C). Myeloid cells in the GBM microenvironment synthesize creatine and export it to neighboring tumor cells, thereby supporting tumoral bioenergetics^54^. While AGAT and GAMT transcripts were expressed by macrophage and microglia populations, we did not observe GAA secretion from cultured microglia (Figure S4D). Taken together, these data and findings from others suggest that myeloid cells in the HGG TME display canonical creatine synthesis pathway activity whereas tumor cells repurpose this pathway to drive GAA secretion. Reinforcing this idea, evaluation of transcript- and protein-level AGAT and GAMT expression in GSC lines and NHA cells revealed tumor cell-autonomous increases in AGAT abundance and the AGAT:GAMT ratio in nearly all GSC cultures compared to astrocytes (Figures 4C-4F). Exceptions included MGG152 and BT260 cell lines that do not synthesize or secrete GAA (Figures 3H-3K).

We next asked if the pervasive increase in AGAT transcription in HGG cells was associated with changes in chromatin accessibility or transcription factor activity at the *GATM* genomic locus (note that the *GATM* gene encodes the AGAT enzyme). We analyzed the expression distribution of AGAT splice isoforms in AGAT^high^ (BT054, HK211, HK213, HK252, HK157, and HK308) and AGAT^low^ (MGG152 and BT260) GSC lines and in NHA cells (Figures S4E and S4F). AGAT^high^ cells nearly exclusively expressed one transcript (ENST00000396659), which is associated with promoter and enhancer elements surrounding exon 1 and an intragenic enhancer located between exons 2 and 3 of the *GATM* gene. We performed ATAC-sequencing of AGAT^high^ GSCs, AGAT^low^ GSCs, and NHA cells (Figure 4G). AGAT^high^ GSCs displayed more open chromatin at these promoter and enhancer sequences. Moreover, footprinting analysis demonstrated increased transcription factor occupancy at these loci in AGAT^high^ GSCs. Therefore, overexpression of the AGAT enzyme in HGGs is linked to chromatin remodeling within the *GATM* gene and enhanced transcription factor recruitment to constituent cis-regulatory elements.

### Imbalanced AGAT and GAMT activities are necessary and sufficient for GAA accumulation

Having shown that GAA accumulation and secretion correlate with an increase in the ratio of AGAT to GAMT expression, we next asked if this relationship extends to primary tumor samples. Using matched proteomics and metabolomics data from 50 human gliomas, we conducted a correlation analysis, comparing protein-level AGAT:GAMT expression with tumoral GAA content (Figure 5A). We observed a positive association between these properties that tracked with tumor grade.

**Figure 5.**
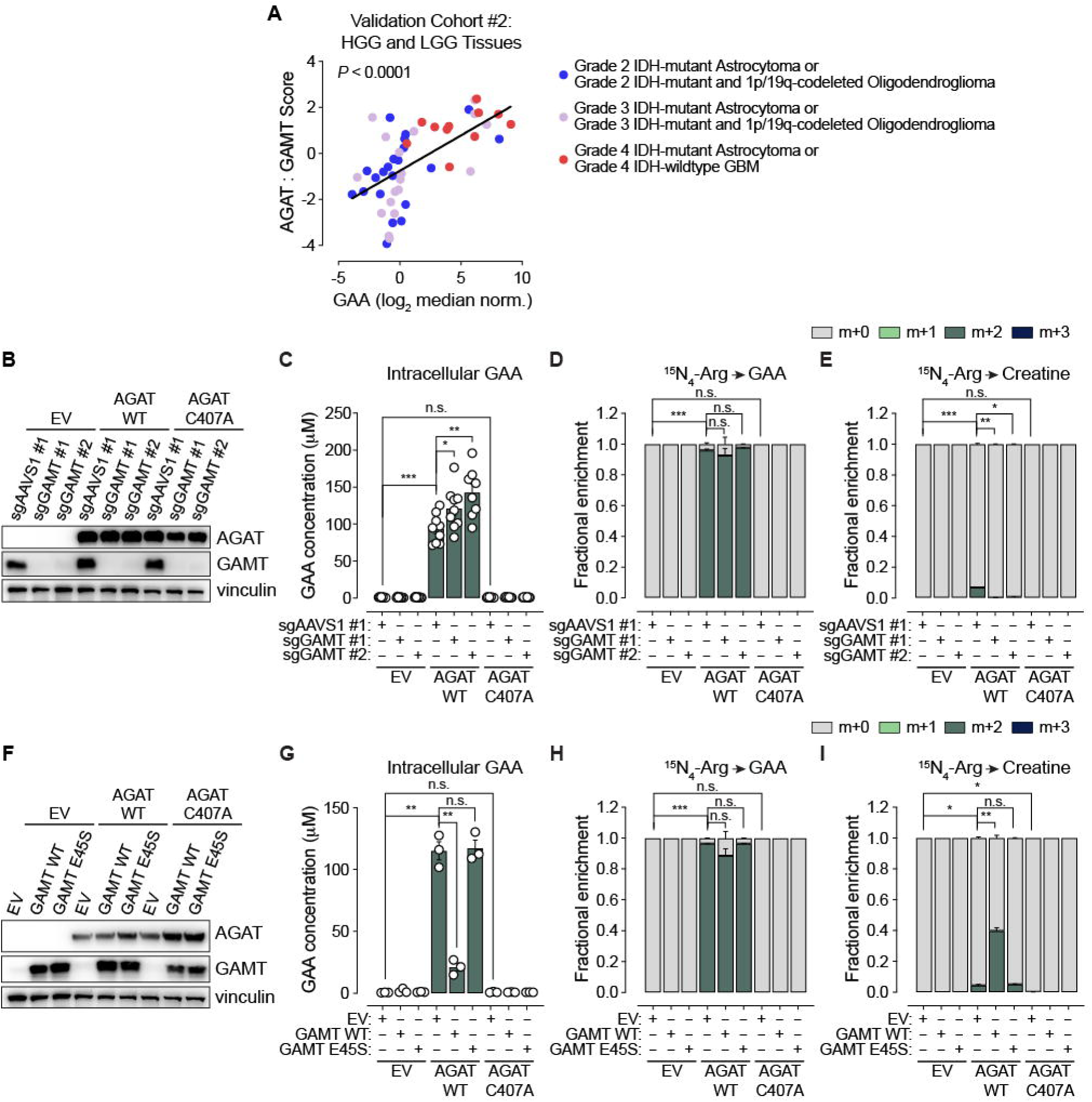
Imbalanced AGAT and GAMT expression is necessary and sufficient for GAA accumulation. **(A)** Correlation of tumor GAA content with AGAT and GAMT expression profiles from Validation Cohort #2. AGAT : GAMT Scores and GAA abundance values are derived from proteomics and metabolomics analyses of primary tissues, respectively (*n=*50). **(B)** Immunoblot analyses of NHA Donor #1 cells engineered to express either empty vector (EV), wild-type AGAT cDNA (AGAT WT), or catalytically-dead AGAT cDNA (AGAT C407A) and Cas9 with an sgRNA targeting either the *AAVS1* safe-harbor locus control or one of two sgRNAs targeting *GAMT* (*n*=3), **(C)** Intracellular GAA content of engineered NHA Donor #1 cells following 48-hour culture in HPLM (*n=*9 for all lines except AGAT WT, sgGAMT #1 cells, for which *n*=8). **(D-E)** Fractional enrichment of cellular (D) GAA and (E) creatine by ^15^N_4_-arginine tracer in engineered NHA Donor #1 cells following 18-hour incubation in HPLM (*n*=3 per line). **(F)** Immunoblot analyses of NHA cells engineered to express either empty vector (EV), wild-type AGAT (AGAT WT), or catalytically-dead AGAT cDNA (AGAT C407A) and either EV, wild-type GAMT cDNA (GAMT WT), or catalytically-dead GAMT cDNA (GAMT E45S) (*n=*3). **(G)** Intracellular GAA content of engineered NHA Donor #1 lines following 48-hour culture in HPLM (*n*=3 for all lines). **(H-I)** Fractional enrichment of cellular (H) GAA and (I) creatine by ^15^N_4_-arginine tracer in engineered NHA Donor #1 cell lines following 18-hour incubation in HPLM (*n*=3 per line). Data are means ± SEM. n.s.= not significant, *p<0.05, **p<0.01, ***p<0.001 (Welch’s *t*-test). See also Figure S5.

To test whether increased AGAT activity relative to GAMT causes GAA accumulation, we transduced natively AGAT^low^ NHA cells with wildtype AGAT, a catalytically dead AGAT point mutant (C407A^55^), or an empty vector control (Figure 5B). To further modulate the ratio of AGAT to GAMT activities, we expressed Cas9 and one of two *GAMT*-targeted sgRNAs or a control sgRNA targeting the *AAVS1* safe harbor locus. Overexpressing wildtype AGAT was sufficient to increase GAA levels (Figure 5C) and triggered robust labeling of GAA, but not creatine, by ^15^N_4_-arginine (Figures 5D, 5E, and S5A). These effects were dependent on the catalytic activity of AGAT because they were abolished by the C407A point mutation. Increasing the ratio of AGAT to GAMT activity further by simultaneously overexpressing wildtype AGAT and knocking out *GAMT* evoked higher levels of GAA accumulation relative to AGAT overexpression alone. Therefore, imbalanced expression and activity of AGAT and GAMT enzymes is sufficient to explain GAA enrichment in human HGGs.

To ask if imbalanced AGAT and GAMT activities are necessary to sustain high cellular GAA levels, we further engineered NHA lines stably expressing AGAT wildtype or C407A mutant enzymes or an empty vector control. We transduced these lines to overexpress wildtype GAMT, a catalytically dead GAMT point mutant (E45S^56^), or an empty vector control (Figure 5F). Increasing GAMT activity in AGAT-overexpressing cells depleted intracellular GAA (Figure 5G). This effect was associated with substantial ^15^N_4_-arginine labeling of both GAA and creatine pools (Figures 5H, 5I, and S5B). In all experiments, the catalytically dead mutants of AGAT and GAMT failed to recapitulate effects of the respective wildtype enzyme. These studies show that low GAMT activity is required for GAA to accumulate upon AGAT upregulation. Moreover, our findings indicate that GAMT occupies a critical node in the creatine synthesis pathway, ultimately dictating whether AGAT activity leads to creatine production or GAA accumulation and secretion.

### GAA-dependent GABA_A_ agonism and chloride dysregulation drive neuronal activity in the HGG microenvironment

Recurrent enrichment of GAA in HGGs but not index LGGs or non-malignant brain tissues (Figures 2C, 3A, and 3E) suggested that this metabolite may play a functional role in promoting aggressive brain tumor behavior. Because HGGs display a bottleneck in the creatine synthesis pathway, we posited that GAA’s role in tumor promotion may be distinct from its canonical role as a creatine precursor. GAA is known to accumulate broadly in tissues of patients with GAMT deficiency, an inborn error of creatine metabolism caused by loss-of-function mutations in the *GAMT* gene (Figure 6A). These patients suffer from intractable seizures, a symptom that is not commonly observed in patients with other inborn errors of creatine metabolism that do not result in GAA accumulation^57^. Seizure promotion suggests that GAA elicits excitatory actions in the brain. Indeed, preclinical studies have shown that GAA regulates neuronal electrochemical signaling by stimulating GABA_A_ receptors^58, 59^ as well as inhibiting synaptic glutamate uptake^60^ and Na^+^/K^+^ ATPase activity^61^. These properties were intriguing because patients with HGG, like those with GAMT deficiency, also experience seizures, cognitive dysfunction, and neurological deficits. Glioma-infiltrated tissues are more electrically active than non-malignant brain tissues^28^, and microenvironmental neuronal activity in turn drives glioma growth through neuron-to-glioma synapses and activity-regulated signaling^26–28, 33, 36, 62, 63^. Although glutamate release by tumor cells represents one process by which glioma cells augment the activity of local neurons^28, 35, 64^, we do not have a comprehensive understanding of the molecular mechanisms driving electrical activity in the HGG microenvironment. Thus, we hypothesized that GAA secretion by HGG cells may represent a critical yet unappreciated mechanism of glioma-neuron crosstalk.

**Figure 6.**
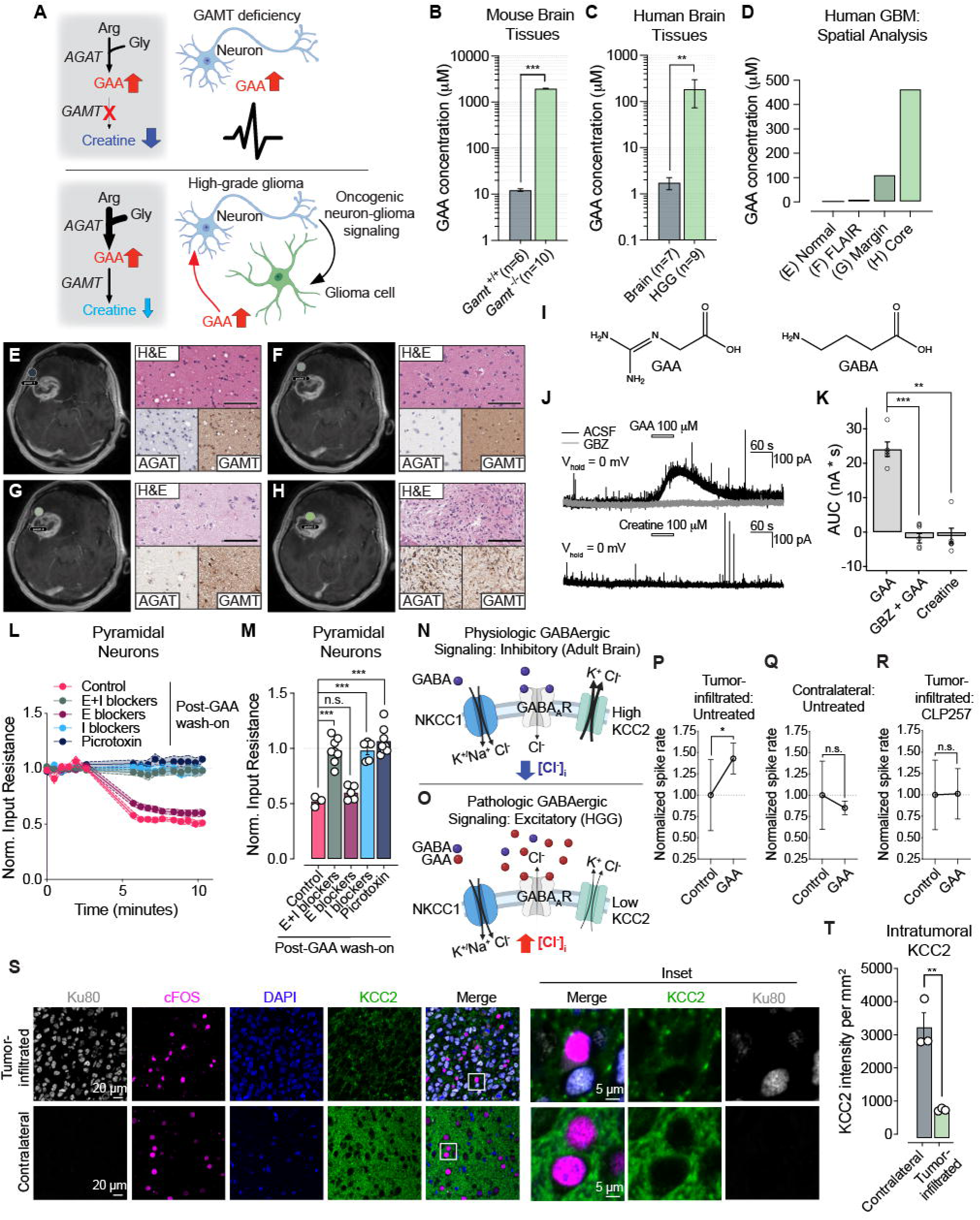
GAA accumulation in the glioma microenvironment activates GABAergic signaling. (**A**) Schema depicting metabolic alterations and phenotypes in GAMT deficiency and HGG. (**B-C**) Tissue GAA levels in (B) a murine model of GAMT deficiency (*n*=10) and littermate controls (*n=*6) or (C) human non-malignant brain (*n=*7) and HGG (*n*=9) tissues. In (B), data are from Schmidt et al^77^. (**D-H**) Spatially resolved GAA content in human GBM. (D) GAA content in samples taken from four distinct points within a single GBM corresponding to (E) largely non-malignant brain (“Normal”), (F) FLAIR-enhancing tumor-adjacent neocortex, (G) tumor margin, and (H) tumor core as assessed on intraoperative magnetic resonance imaging (MRI). (E-H) MRI, hematoxylin and eosin (H&E), and immunohistochemistry (IHC) for AGAT and GAMT of sampled tissues in (D). Scale bar = 50 µm. (**I**) Chemical structures of GAA and γ- aminobutyric acid (GABA). (**J**) Representative traces of patch-clamp recordings of neurons in murine brain slices treated with 100 µM GAA or creatine in 0 µM or 10 µM of the GABA_A_ receptor inhibitor gabazine. (**K**) Integrated area under the curve (AUC) of patch-clamp traces for slices treated with 100 µM GAA (*n*=5), 100 µM GAA + 10 µM gabazine (*n*=6), or 100 µM creatine (*n*=5). (**L-M**) Normalized input resistance of neurons treated with GAA alone (*n*=3), 10 µM of the AMPA receptor inhibitor 3-(2-carboxypiperazin-4-yl)propyl-1-phosphonic acid (CPP) and 20 µM of the NMDA receptor inhibitor 6,7-dinitroquinoxaline-2,3-dione (DNQX; E blockers, *n*=5), 100 µM of the GABA_A_ receptor inhibitor picrotoxin and 2 µM of the GABA_B_ receptor inhibitor CGP52432 (I blockers, *n=*5), all inhibitors (E+I blockers, *n*=8), or picrotoxin alone (*n*=8). In (L), data depict changes in normalized input resistance over time following GAA wash-on. In (M), data depict average post-wash-on normalized input resistance. Data are normalized to average input resistance of each neuron prior to GAA wash-on. (**N-O**) Schemas depicting maintenance of [Cl^-^]_i_ by NKCC1 and KCC2 in (N) the brain under normal physiologic conditions and (O) HGGs that display pathologic KCC2 downregulation. (**P-R**) Multielectrode array (MEA) assay of neuronal activity in response to GAA treatment in tumor-bearing brain slices. Spike rates normalized to mean pre-treatment spiking rates, prior to and following treatment with 20 µM GAA for (P) tumor-infiltrated right hemisphere (*n*=3 slices), (Q) tumor-free contralateral cortex (*n*=3 slices), and (R) tumor-infiltrated neocortex treated with 30 µM CLP-257 (*n*=3 slices). (**S-T**) Immunofluorescence analysis of cell markers and KCC2 expression in the tumor microenvironment. (S) Representative images of markers Ku80 (human GBM cells), cFOS (neuronal activity), and DAPI (nuclei) as well as KCC2 expression in tumor-infiltrated and contralateral neocortex. (T) Quantification of KCC2 expression in tumor-infiltrated and contralateral neocortex (n=3 mice). Data are means ± SEM. n.s. = not significant, *p<0.05, **p<0.01, ***p<0.001 (Welch’s *t*-test in B-C, unpaired *t*-test in M and T, paired *t*-test in P-R). See also Figure S6.

To better understand the pathophysiological relevance of GAA accumulation in the contexts of GAMT deficiency and HGG, we quantified concentrations of GAA in tissue specimens from HGG and non-malignant brain samples, comparing these values with reported GAA concentrations in brain tissues of *Gamt-*wildtype and *Gamt-*knockout mice (Figures 6B and 6C). Although control samples showed species-specific numerical differences in GAA content, we observed ∼100-fold increases in GAA levels in brain tissues from both *GAMT*-knockout mice and HGG patients relative to these controls. Therefore, it is plausible that GAA accumulation in the HGG microenvironment surpasses a threshold that elicits neuronal excitation in the setting of GAMT deficiency. To further elucidate local GAA concentrations in human GBM, we again leveraged stereotactic neurosurgical navigation to collect normal-appearing, FLAIR-positive, and tumor margin and core tissue samples from a patient undergoing GBM resection (Figure 6D-6H). We found that GAA levels sharply rose with increasing proximity to the tumor core, an effect that was associated with an increase in the ratio of AGAT to GAMT protein expression (as measured by validated immunohistochemistry assays, Figures S6A and S6B). Critically, GAA content of the tumor margin, where interactions between tumor cells and neurons are heightened, was orders of magnitude higher than normal-appearing brain tissue.

We investigated mechanisms through which GAA may contribute to increased electrochemical signaling in the glioma microenvironment. GAA bears striking structural similarity to the neurotransmitter GABA (Figure 6I), a neurotransmitter than can be hyperpolarizing and inhibitory or depolarizing and excitatory, depending on the intracellular concentration of chloride. Whole-cell voltage-clamp recordings (V_hold_ = 0 mV) from CA1 pyramidal neurons in acute mouse hippocampal slices revealed that exogenous GAA application (but not creatine) induced a large outward current that was completely blocked by the GABA_A_ receptor inhibitor gabazine (GBZ) (Figures 6J and 6K). Similarly, current-clamp recordings from cortical pyramidal neurons in acute slices from adolescent murine brain revealed that GAA reduced neuron input resistance, consistent with ion channel opening (Figures 6L and 6M). Co-treatment with blockers of GABAergic, but not glutamatergic, receptors fully prevented the decrease in input resistance. GABA_A_ receptor blockade explained this effect, because treating slice cultures with the GABA_A_-selective inhibitor picrotoxin alone was sufficient to completely reverse the activity of GAA on input resistance/channel opening (Figures 6L and 6M). Together, these results support the function of GAA as a GABA_A_ receptor agonist, opening the GABA_A_ receptor channel.

Our findings presented an apparent paradox: GAA engaged GABA receptors and elicited hyperpolarization of mature murine neurons but also accumulates dramatically in the hyperexcitable HGG microenvironment. Intracellular chloride ion concentrations ([Cl^-^]_i_) in neurons dictate whether GABA receptor agonism is depolarizing or hyperpolarizing. NKCC1 and KCC2 are chloride transporters that play important roles in importing and exporting Cl^-^ from neurons, respectively. Healthy adult neurons express high levels of KCC2 and low levels of NKCC1, thereby reducing [Cl^-^]_i_ and resulting in chloride influx and neuronal hyperpolarization upon GABA receptor opening (Figure 6N). Interestingly, prior research has shown that GBM-infiltrated brain displays hyperexcitability in part due to dysregulation of [Cl^-^]_i_ homeostasis^65, 66^. Neurons in the GBM-infiltrated brain display reduced expression of KCC2, increased chloride levels, and depolarizing (excitatory) GABA signaling (Figure 6O). Therefore, we asked if GAA accumulation in HGGs cooperates with tumor-specific [Cl^-^]_i_ dysregulation to drive excitatory GABAergic neurotransmission in glioma-associated neurons. We established orthotopic TS516 xenografts in immunodeficient mice displaying tumor infiltration throughout one hemisphere but not the other. In acute cortical slices from these mice, high-density multi-electrode array (MEA) analysis revealed that GAA wash-on led to an increase in neuronal firing in tumor-infiltrated brain tissue but not in the contralateral hemisphere (Figures 6P, 6Q, and S6C). Treating tumor-infiltrated brain slices with the KCC2 activator CLP257 to decrease [Cl^-^]_i_ abolished the excitatory effect of GAA in the tumor microenvironment (Figure 6R). In agreement with prior work^65, 66^, neuronal cells exhibited markedly lower expression of KCC2 in the brain hemisphere infiltrated by TS516 GBM cells (marked by human antigen Ku80) compared to the contralateral side (Figures 6S and 6T). Together, our findings establish that interstitial GAA accumulation promotes neuronal activity in gliomas in a GABA_A_ receptor- and [Cl^-^]_i_-dependent manner.

### AGAT inhibition represses electrochemical signaling and glioma growth

Neuronal activity plays an important role in driving glioma cell proliferation^26^. Therefore, we sought to determine whether inhibiting AGAT-dependent GAA synthesis and consequent neuronal activation attenuates GBM growth. To genetically repress AGAT, we sequentially transduced TS516 GBM GSCs with adenoviral expression vectors to transiently express Cas9 and two sgRNAs targeting the *GATM* gene (which encodes the AGAT enzyme) or two sgRNAs targeting the *AAVS1* safe harbor locus. We established isogenic AGAT-expressing (AGAT WT) and AGAT-knockout (AGAT KO) bulk TS516 cell cultures (Figure 7A). This approach allowed us to achieve nearly complete silencing of AGAT in the latter without resorting to single cell cloning, which can be challenging in GSC lines that display long doubling times. As expected, AGAT knockout reversed intracellular GAA accumulation and GAA secretion (Figures 7B and 7C). Although AGAT inhibition did not affect the growth of TS516 monocultures in vitro (Figure 7D), it did prolong survival of mice bearing TS516 orthotopic GBM xenografts (Figure 7E), underscoring the central role of the neural tumor microenvironment to this mechanism. To assess the effects of AGAT knockout on tumor metabolite content, we developed assays to assess tissue and GAA and creatine content (Figure S7A, S7B). The survival benefit seen in AGAT KO tumors was associated with depletion of tumoral GAA but not creatine (Figures 7F and 7G), consistent with minimal de novo creatine synthesis in HGG cells (Figure 3K). These data indicate that GAA accumulation drives HGG growth in a microenvironment-dependent manner. Importantly, this oncogenic function of AGAT activity in glioma cells cannot be explained by canonical effects on creatine synthesis.

**Figure 7.**
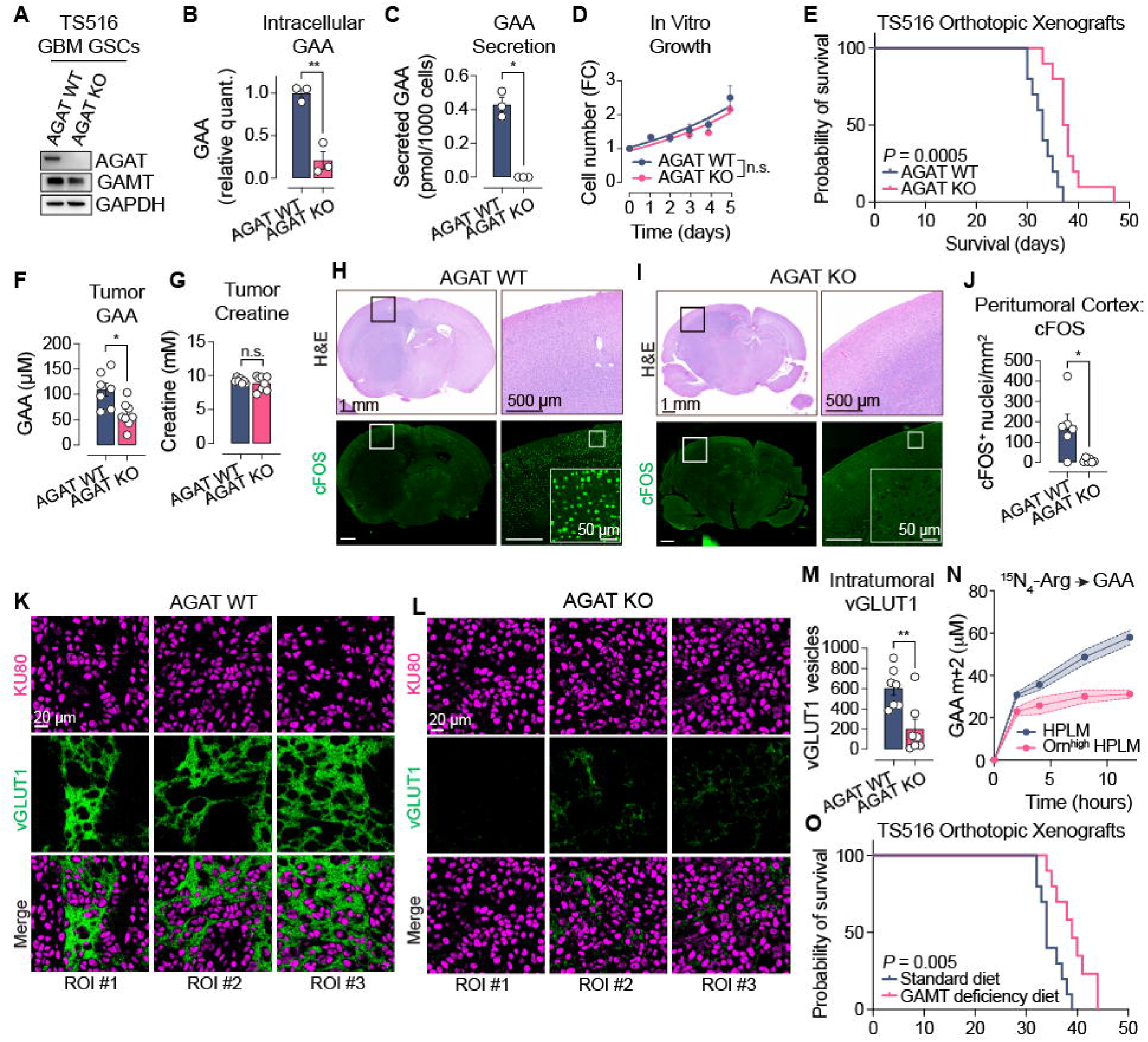
GAA accumulation drives neuronal hyperexcitability and glioma aggressiveness. (**A**) Representative immunoblots of AGAT, GAMT, and GAPDH in AGAT wild-type (WT) and AGAT knockout (AGAT KO) TS516 GSCs. (**B**) Intracellular GAA in AGAT WT and AGAT KO TS516 GSCs after 48-hour culture in HPLM (*n*=3 per cell line). (**C**) GAA secretion by AGAT WT and AGAT KO TS516 GSCs during 48-hour culture in HPLM (*n*=3 per cell line). (**D**) In vitro growth assay of AGAT WT and AGAT KO TS516 GSCs (*n*=3 per time point per cell line). Growth curves are generated by nonlinear exponential regression. (**E**) Overall survival of mice bearing AGAT WT or AGAT KO TS516 xenografts (*n*=10 per arm). (**F**-**G**) Absolute (F) GAA and (G) creatine content for tumors from (E) (*n=*7 for AGAT WT, *n*=8 for AGAT KO). (**H-I**) Representative H&E and cFOS immunofluorescence for (H) AGAT WT and (I) AGAT KO TS516 tumors. Scale bars = 1 mm, 500 µm, or 50 µm. (**J**) Quantification of cFOS^+^ nuclei in peritumoral cortex in AGAT WT and AGAT KO TS516 tumors (*n*=6 per tumor type). (K-L) Immunofluorescent imaging of AGAT WT and AGAT KO TS516 tumors. Scale bars = 20 µm. (**M**) Quantification of vGLUT1 positive foci in predetermined regions of interest within tumor beds of AGAT WT and AGAT KO TS516 tumors (*n*=7 per genotype). (**N**) Intracellular content of the M+2 isotopologue of GAA following ^15^N_4_-arginine tracing in the presence or absence of 150 µM ornithine supplementation (*n*=3 per condition). (**O**) Survival of TS516 xenograft bearing mice receiving a standard (*n*=10) or an ornithine-supplemented, arginine-restricted (GAMT deficiency, *n*=10) diet. Data are means ± SEM. n.s.= not significant, *p<0.05, **p<0.01 (Welch’s *t*-test in B, C, and J; log-rank test in E and O; unpaired *t*-test in F, G, and M). See also Figure S7.

We next evaluated whether AGAT knockout blocks neuronal hyperactivity in the HGG microenvironment. We quantified nuclei expressing cFOS^+^, an immediate early gene indicative of recent neuronal activity, in cortical sections of tumor-infiltrated brain tissues harvested from mice bearing AGAT WT or AGAT KO xenografts. Electrochemically active cells marked by cFOS expression were enriched in GAA-replete, AGAT WT glioma xenograft tissues and reduced in GAA-depleted, AGAT KO comparators (Figures 7H-7J and S7C). The majority of cFOS^+^ cells in the microenvironment of AGAT WT xenografts also expressed NeuN (Figures S7C and S7D), affirming their neuronal identity. Consistently, we also observed increased levels of vGLUT1, a marker of excitatory synapses, in tumor-infiltrated brain regions of mice with AGAT WT versus AGAT KO xenografts (Figures 7K-7M, S7E, and S7F), suggesting that local hyperexcitability induced by GAA promotes interactions between glutamatergic neurons and tumor cells in the HGG microenvironment.

Results from studies involving genetic manipulation of AGAT support therapeutically targeting AGAT for HGG treatment but do not outline a tractable means to do so. Interestingly, a dietary therapy that attenuates AGAT activity and GAA accumulation is effective in reducing neuronal hyperactivity and seizure incidence in patients with GAMT deficiency^67, 68^. This diet features reduced arginine content and high levels of ornithine, which reduce AGAT substrate availability and promote end-product inhibition of AGAT, respectively. Given the efficacy of this diet in GAMT deficiency, we tested its therapeutic activity against HGGs. We first modeled the impact of ornithine supplementation on AGAT function by measuring ^15^N_4_-arginine labeling of GAA and found that high levels of ornithine reduced AGAT activity in cultured TS516 cells (Figure 7N). Next, we tested tolerability and antitumor efficacy of an arginine-depleted and ornithine-supplemented diet (hereafter the “GAMT deficiency diet”). Mice administered the GAMT deficiency diet did not display weight loss nor reduced chow consumption relative to mice fed a standard diet (Figures S7G and S7H). Critically, placing mice bearing TS516 orthotopic xenografts on the GAMT deficiency diet prolonged their survival relative to mice given a standard diet (Figure 7O). The survival effect size of the GAMT deficiency diet was similar to the effect size associated with genetic inhibition of AGAT (Figure 7E). Therefore, this dietary therapy may constitute an effective, clinically validated approach to block aberrant AGAT activity and oncogenic GAA signaling in patients with HGG.

## DISCUSSION

We generated a deep metabolite profiling dataset that spanned primary brain tumors, brain metastases, and non-malignant brain specimens to discover GAA accumulation as a recurrent metabolic feature of HGG that manifests independently of canonical genetic and transcriptomic disease subtypes. We went on to validate this finding in two independent cohorts. Unlike GAA, creatine and creatine derivatives were downregulated or unchanged in HGGs compared to non-malignant brain tissues. Stable isotope tracing studies revealed that these metabolic changes were tied to a bottleneck in the de novo creatine synthesis pathway in GSC lines that functionally disconnects GAA production from creatine synthesis. Our findings reveal conceptual parallels between accumulation of GAA in HGGs and enrichment of the oncometabolite (*R*)-2HG in IDH-mutant gliomas. Both metabolites reach levels in glioma tissues that are at least 1-2 orders of magnitude higher than those in non-malignant brain and display enrichment profiles that map onto clinically relevant glioma features. Specifically, increased GAA content serves as a surrogate marker of tumor grade while (*R*)-2HG accumulation indicates the presence of an *IDH1* or *IDH2* mutation. These properties suggest that GAA, like (*R*)-2HG, may have utility as a glioma tissue biomarker in diagnostic and therapeutic applications, including spectroscopic imaging and intraoperative mass-spectrometry based margin detection^43, 44, 69–71^. From a biological standpoint, GAA and (*R*)-2HG share another feature: both metabolites function as signaling molecules rather than serving as substrates for anabolic or bioenergetic processes in tumor cells. These insights reinforce the importance of metabolite signaling in the molecular pathogenesis of glioma.

Findings from our work and prior research indicate that the creatine synthesis pathway is regulated in a highly cell type-specific manner in HGGs. Tumor-associated macrophages and microglia provide a local supply of creatine to GBM cells that promotes tumor tolerance to microenvironmental stressors, including hypoxia and acidification^54^. This interaction may allow HGG cells to repurpose the creatine synthesis pathway to produce and secrete GAA, a finding from our work that aligns with observation of GAA accumulation in HGG interstitial fluid in a previous study^47^. Our data demonstrate that GAA secretion by HGG cells is explained by AGAT upregulation in the absence of a commensurate increase in GAMT. RNA seq comparisons of human HGG and reference brain showed broad AGAT upregulation with reduced expression of GAMT and distal enzymes across transcriptional subtypes and IDH genotypes. In patient derived GSCs, AGAT levels and the ratio of AGAT:GAMT expression rose relative to non-malignant neural cells, which was associated with increased chromatin accessibility and transcription factor occupancy at key cis-regulatory elements in the *GATM* gene. In primary tumors, the ratio of AGAT:GAMT protein expression positively correlated with tissue GAA content and tumor grade. Gain and loss of function studies support this model: AGAT overexpression raised cellular GAA and ^15^N arginine flux into the GAA pool in a catalytic activity dependent fashion, while concurrent GAMT knockout or overexpression amplified or reversed these effects, respectively. Therefore, we propose that imbalanced AGAT and GAMT activities saturate the catalytic capacity of GAMT to convert GAA to creatine, leading to GAA buildup and secretion from HGG cells.

Human genetics and pathophysiology underscore the clinical relevance of GAA accumulation. In GAMT deficiency, an inborn error of creatine metabolism marked by systemic GAA accumulation, patients present with seizures, cognitive impairment, and psychomotor deficits^72^ that mirror the clinical phenotype commonly seen in glioma. GAA has been shown to regulate neuronal activity by serving as an agonist for GABA_A_ receptors^58, 59^ and repressing Na^+^/K^+^ ATPase activity^60^ and synaptic glutamate uptake^61^. The standard therapy for GAMT deficiency involves restricting arginine (an AGAT substrate) and supplementing creatine and ornithine (an end-product AGAT inhibitor). This treatment causes GAA depletion, electroencephalogram normalization, and decreased seizure frequency in GAMT deficiency patients^67, 68^. These results are not achieved by creatine supplementation alone and intractable seizures are uncommon in other inborn errors of creatine metabolism caused by loss of function mutations in genes encoding the SLC6A8 creatine transporter or AGAT^50^, thereby reinforcing the link between GAA accumulation and neuronal hyperactivity. We found that inhibiting GAA synthesis by genetic or dietary approaches constrained HGG growth in vivo: CRISPR knockout of AGAT in orthotopic GBMs decreased tumoral GAA and markers of neuronal activity and neuron-glioma interactions and extended host survival without altering the growth of in vitro GBM cell monocultures. Inspired by clinical management of GAMT deficiency, we modeled an arginine restricted, ornithine supplemented diet. After confirming that ornithine accumulation inhibited GAA synthesis in vitro, we tested this diet in mice and found that it was well tolerated and recapitulated effects of genetic AGAT inhibition on survival of glioma-bearing mice. These observations underscore the neuroactivity of GAA and suggest that dietary interventions used in the clinical management of GAMT deficiency may display therapeutic activity in patients with HGG.

Advances in cancer neuroscience demonstrate that gliomas elevate neuronal activity of glioma-infiltrated circuits to further augment neuron-glioma interactions and accelerate tumor growth^26–28, 31, 33, 62, 63^. Concordantly, tumor-infiltrated brain tissues are more electrically active than healthy tissue^31^. However, the mechanisms through which glioma cells enhance neuronal firing are only beginning to be elucidated. Release of glutamate^28, 34^ and synaptogenic factors^36, 37, 62^ are implicated in this process, but the full complement of molecules involved in glioma-to-neuron communication is still emerging. Our data implicate GAA accumulation as a novel mechanism of neuronal activation in HGG. GAA is structurally similar to GABA, a classically inhibitory neurotransmitter. Our results in acute cortical and hippocampal brain slices confirm previous studies that GAA is a GABA_A_ receptor agonist^58, 59^. Recent work has shown that GABAergic signaling promotes progression of diffuse midline gliomas^73^ and that [Cl^-^]_i_ dysregulation converts GABAergic signaling from inhibitory to excitatory within the GBM microenvironment^65, 66^. Mechanistically, these changes in GBM are tied to KCC2 downregulation in tumor-resident neurons. We therefore hypothesized that chloride dysregulation and GAA accumulation cooperate to drive excitatory GABAergic signaling in HGG-infiltrated brain. In glioma bearing brain slices, GAA preferentially increased firing within tumor infiltrated regions. This effect of GAA was abrogated with pretreatment of the KCC2 activator CLP-257. Finally, we observed diminished KCC2 expression and a GAA-dependent increase in electrochemical signaling markers in xenograft HGG tissues. These findings support a model in which extracellular GAA produced by HGG cells activates GABA_A_ currents that are excitatory under an aberrant local chloride set point.

Together, our results define how GAA becomes enriched in HGGs, establish its origin in tumor cells, connect its accumulation to neuronal hyperexcitability, and outline a tractable treatment strategy to disrupt metabolite signaling that drives neuronal activity-dependent glioma growth.

## ACKNOWLEDGEMENTS

This study was supported by National Institutes of Health (NIH) grants R01CA258586 and R01CA289260 to S.K.M. and K.G.A., R01NS142141 to S.K.M., P50CA165962 and U19CA264504 to S.K.M., K99CA277576 to Y.X., and R35CA220449 to R.J.D. This work was also supported by awards from Oligo Nation to S.K.M. and K.G.A., Cancer Prevention and Research Institute of Texas (CPRIT) grants RR190034 and RP230344 to S.K.M., CPRIT grant RP2400489 to S.K.M. and B.L., a Distinguished Scientist Award from the Sontag Foundation to S.K.M., and by a Human Frontier Science Program (HFSP) postdoctoral fellowship award (LT0018/2022-L) to Y.X. D.D.S. was supported by NIH/NCI K12CA0903354, a Burroughs Wellcome Career Award for Medical Scientists, and a Lubin Family Foundation Scholar Award. L.G.Z., T.P.M., and the Children’s Research Institute Metabolomics facility are supported by CPRIT grant RP240494. M.R.S. was supported by NIH/NCI grant F30CA271634. K.G.A received funding from CPRIT RP210140 as well as NIH grant P30CA047904 awarded to UPMC Hillman Cancer Center. S.A.W. is supported by CPRIT Training Grant RP210041. This project used the service of the University of Pittsburgh Small Molecule Biomarker Core facility, which was supported, in part, by the University of Pittsburgh Office of the Senior Vice Chancellor, Health Sciences, and NIH grants S10RR023461 and S10OD028540. This project used Zeiss Axioscan 7 in the Whole Brain Microscopy Facility (WBMF) at the UT Southwestern Medical Center, which is supported by the NIH grant 1S10OD032267-01 (to Denise Ramirez). Some data used were generated by the TCGA Research Network (https://www.cancer.gov/tcga) or by the Genotype-Tissue Expression (GTEx) Project, which was supported by the Common Fund of the Office of the Director of the National Institutes of Health, and by NCI, NHGRI, NHLBI, NIDA, NIMH, and NINDS. Some figures were constructed using BioRender. The authors wish to express their gratitude to all patients who contributed to this study.

## AUTHOR CONTRIBUTIONS

Conceptualization: K.G.A. and S.K.M.

Methodology: K.G.A., K.M., C.K.E., S.A.W., Y.X., M.R.S., M.T.G., Y.-T.H., J.I.T., D.D.S., M.T., N.M., V.T.P., R.E.W.III, A.K., T.D.N., B.J.K., N.M.C., C.W., D.R.M., M.A.G., T.P.M., J.R.G., K.M.H. and S.K.M.

Investigation: K.M., C.K.E., S.A.W., Y.X., M.R.S., M.T.G., S.E.K., Y.-T.H., J.I.T., K.G., W.H.H., M.T., M.M.L., M.E.-S., B.C.S., V.T.P., P.K., T.S., C.L., L.G.Z., F.C. and S.A.

Formal analysis: K.M., C.K.E., S.A.W., Y.X., M.R.S., M.T.G., Y.-T.H., J.I.T., L.G., N.M., K.J.H., T.E.R. and S.K.M.

Data curation: S.A.W., Y.X., M.R.S., B.J.K., N.M.C. and C.W.

Software: M.R.S. and L.G.

Validation: K.M., C.K.E., S.A.W. and Y.X.

Resources: K.G.A., T.R.P., B.C.L., P.O.Z., D.R.M., M.A.G., M.A.C., L.X. and O.H.K.

Supervision: K.G.A., P.R., S.M., T.D.N., D.R.M., M.A.G., L.X., J.-A.L., K.M.H., S.C., R.J.D. and S.K.M.

Project administration: K.G.A., S.O., P.K. and S.K.M.

Writing-Original draft: K.G.A., K.M., C.K.E., S.A.W., X.Y. and S.K.M.

Writing-Review & Editing: K.G.A., C.K.E., S.A.W., Y.X., M.R.S., D.D.S., T.D.N., M.A.C., B.L., M.M., K.M.H., S.C., R.J.D. and S.K.M.

Funding acquisition: B.L., K.G.A. and S.K.M.

## STAR METHODS

### RESOURCE AVAILABILITY

#### Lead Contact

- Further information and requests for resources and reagents should be directed to and will be fulfilled by the Lead Contact, Samuel K. McBrayer.

#### Materials Availability

- Plasmids generated in this study will be deposited to Addgene prior to publication.

#### Data and Code Availability

- Metabolomics data will be deposited to the National Metabolomics Data Repository (NMDR) and will be publicly available as of the date of publication. Accession numbers will be listed in the key resources table. All other data reported in this paper will be shared by the Lead Contact upon request.
- This paper does not report original code.
- Any additional information required to reanalyze the data reported in this paper is available from the Lead Contact upon request.

#### Key Resources Table

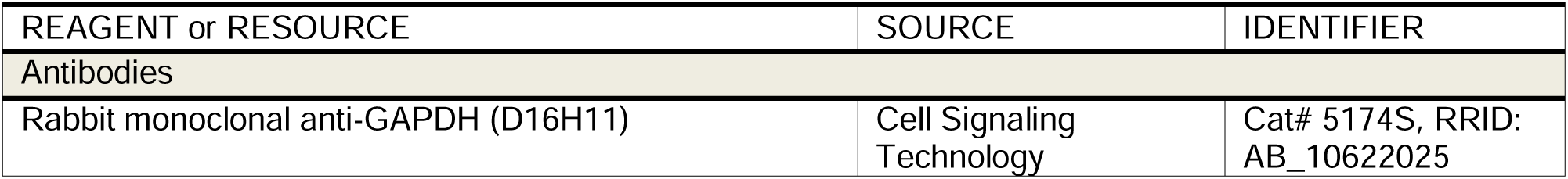
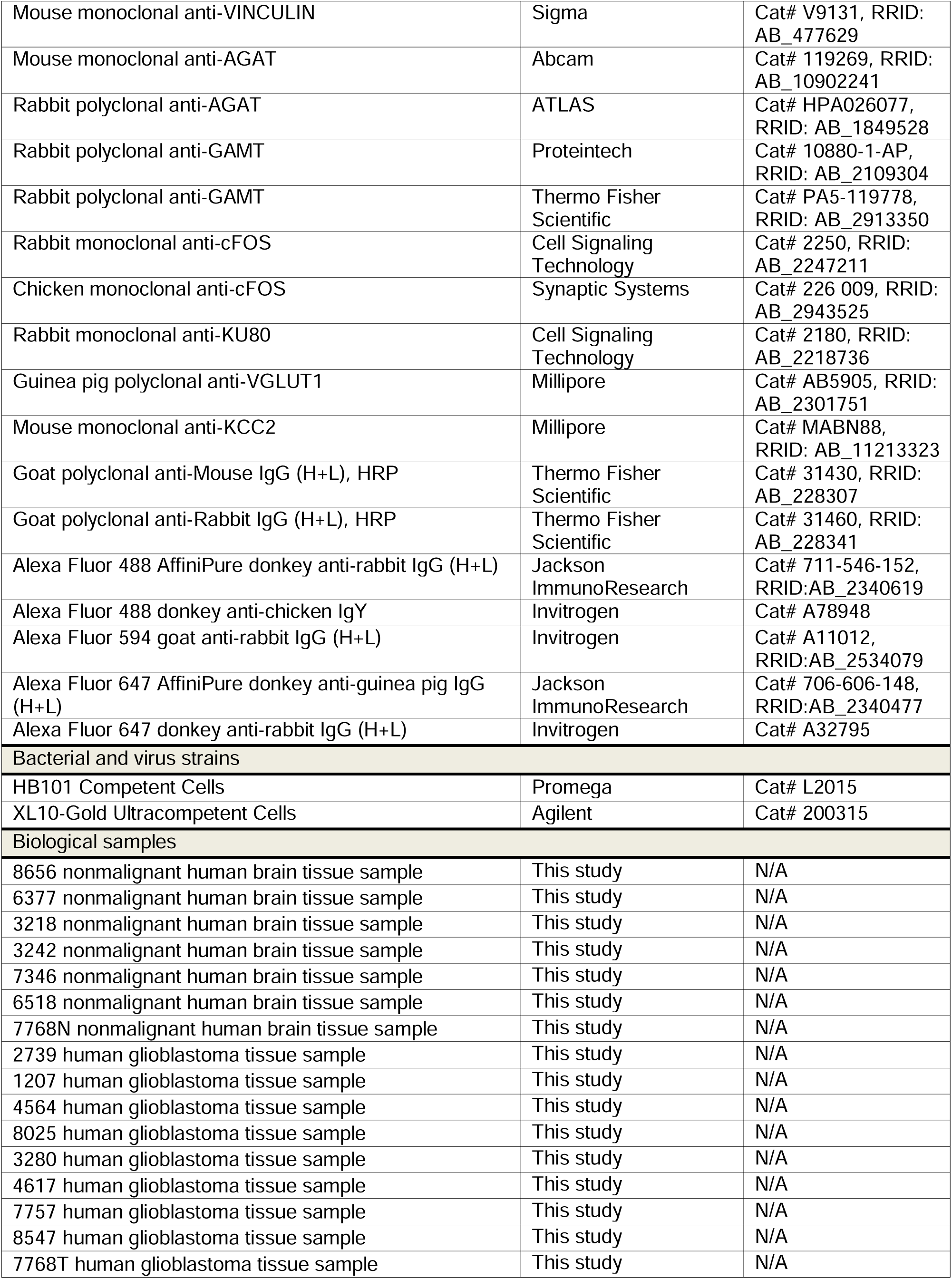
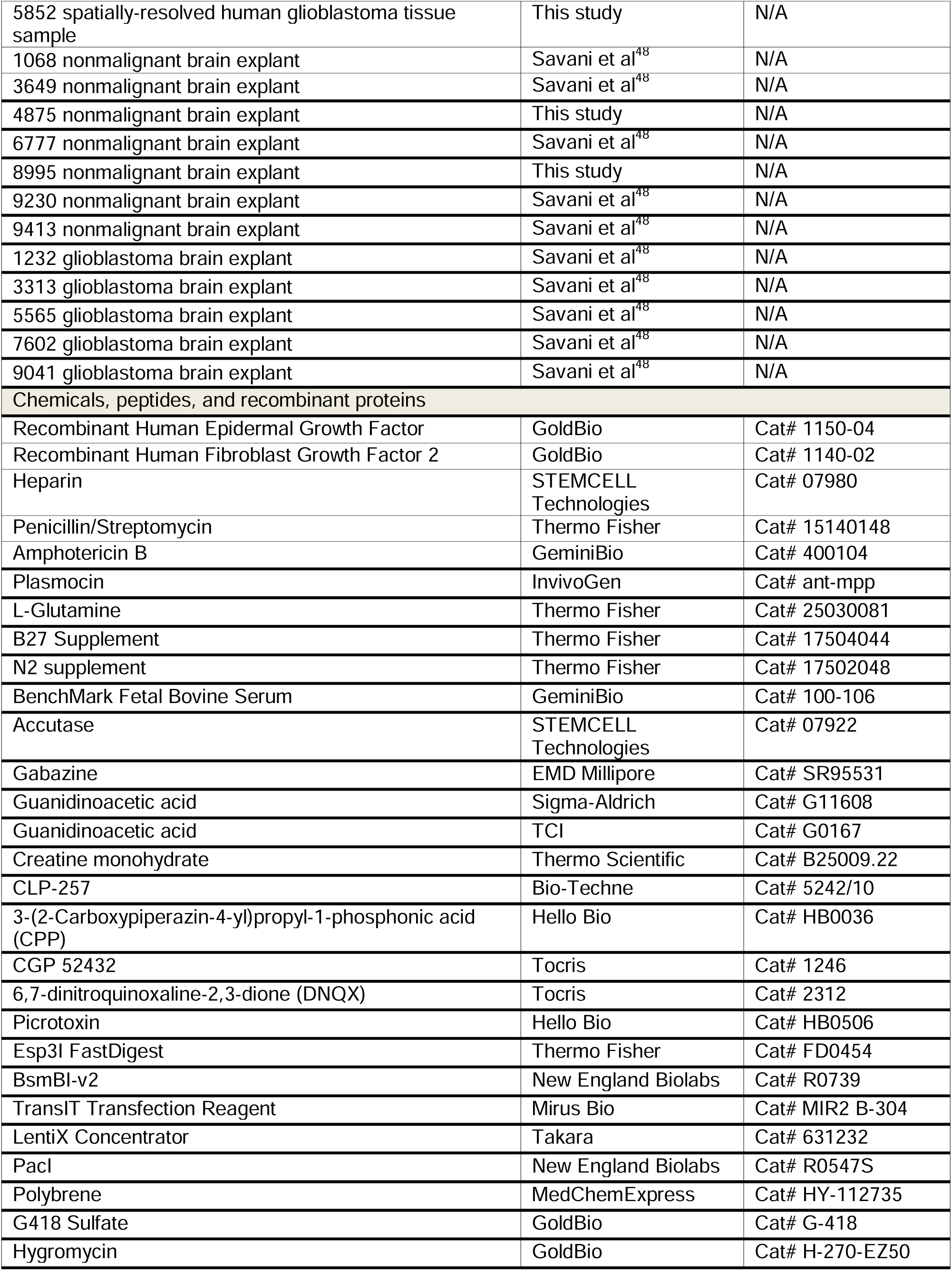
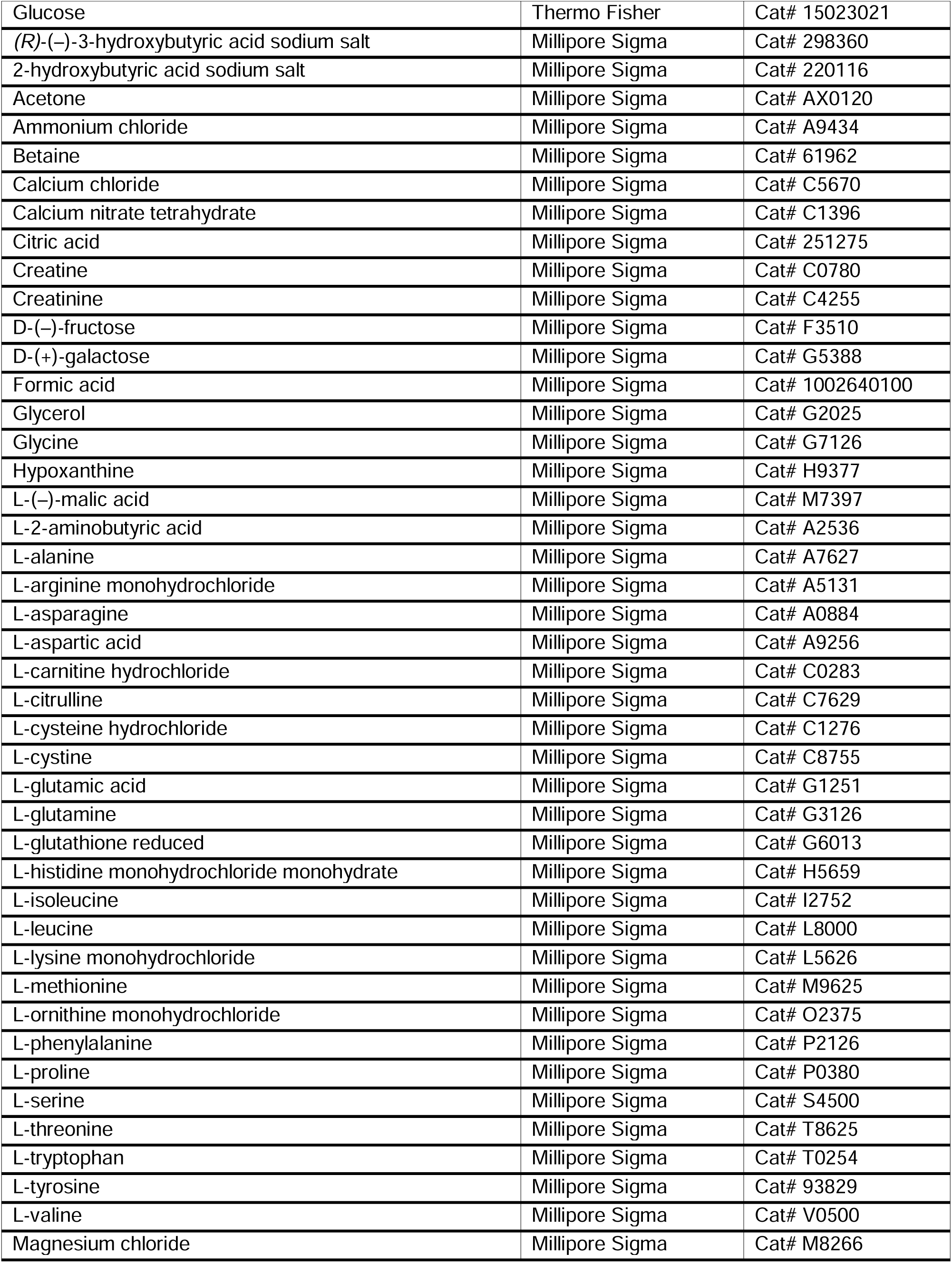
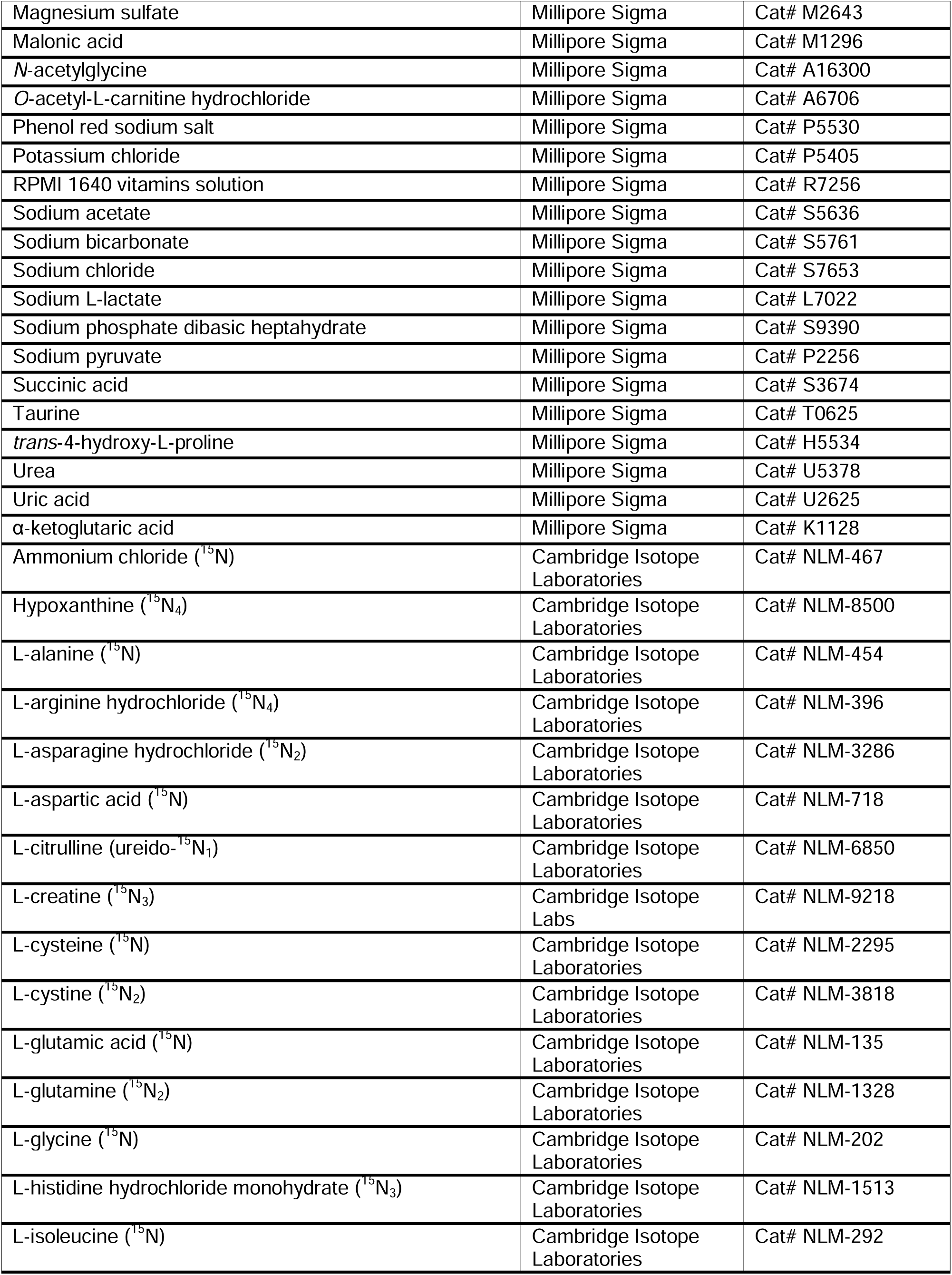
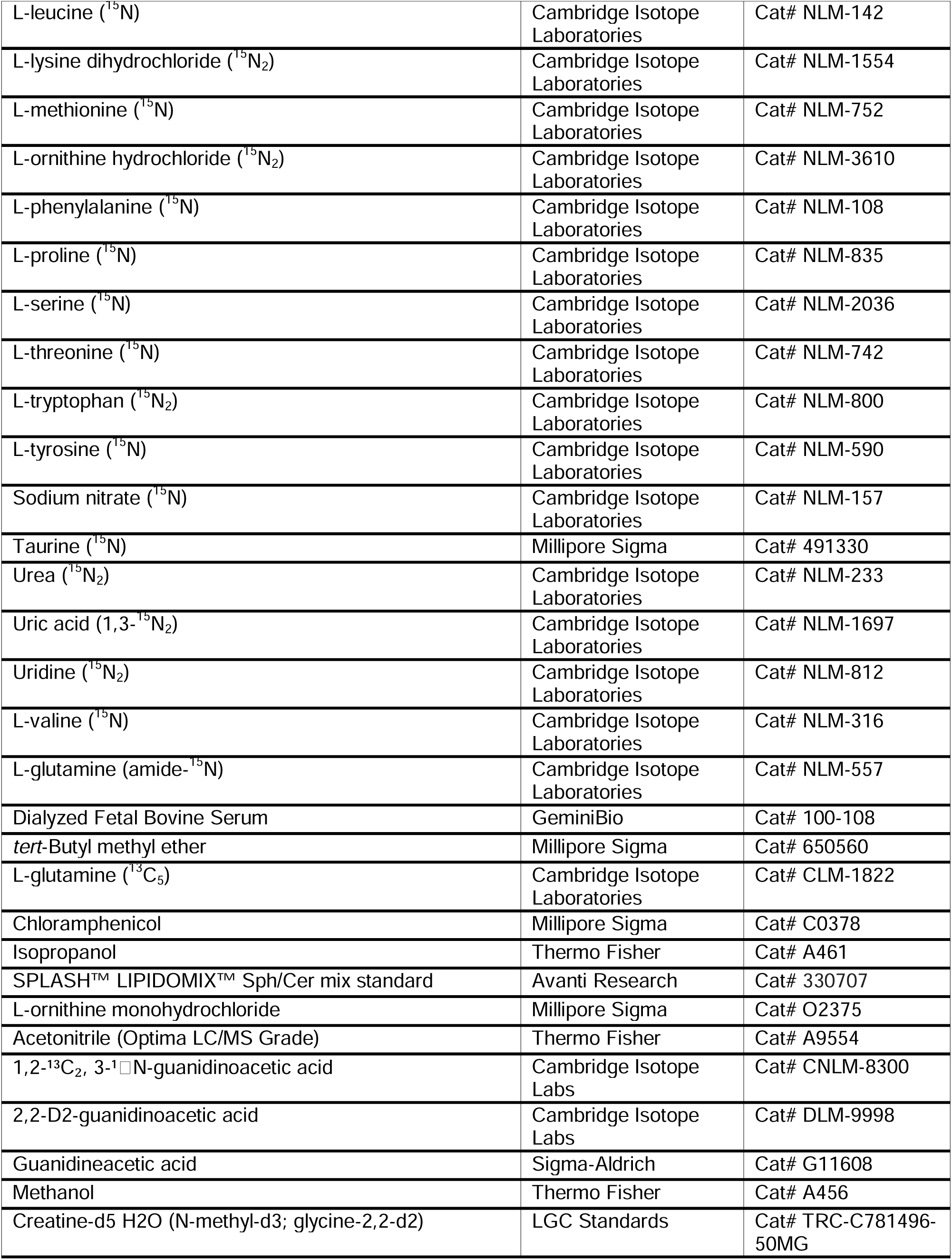
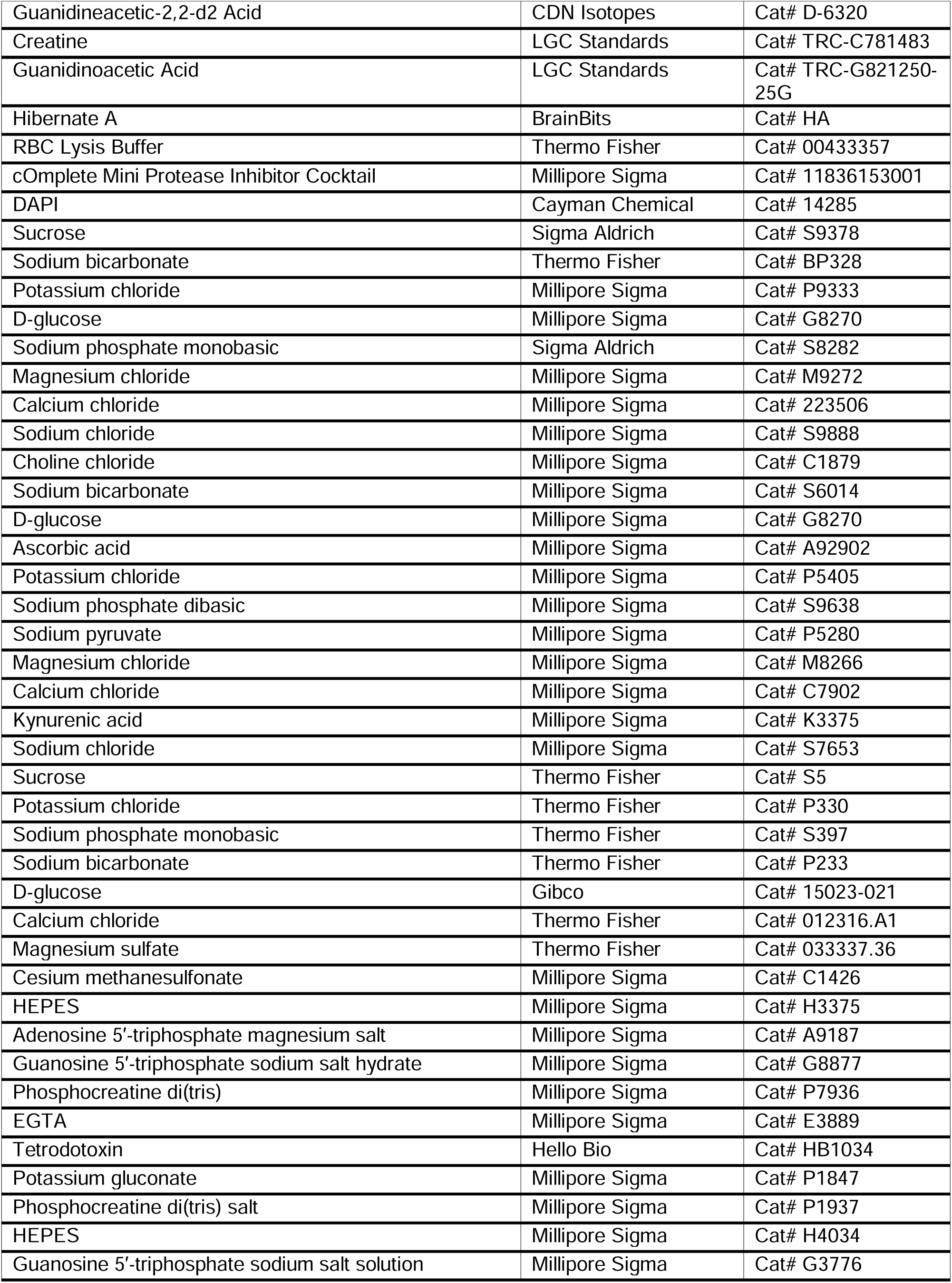
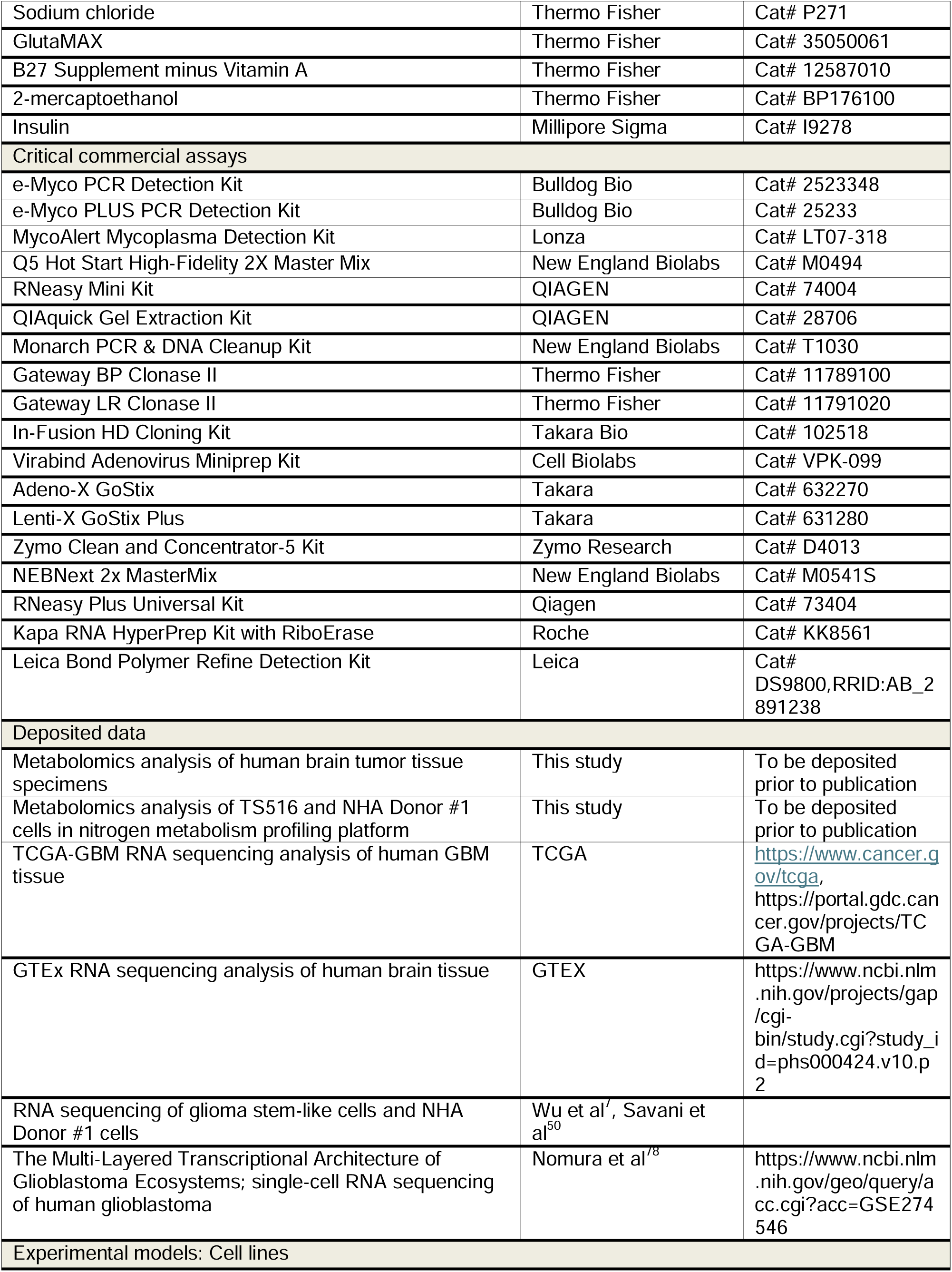
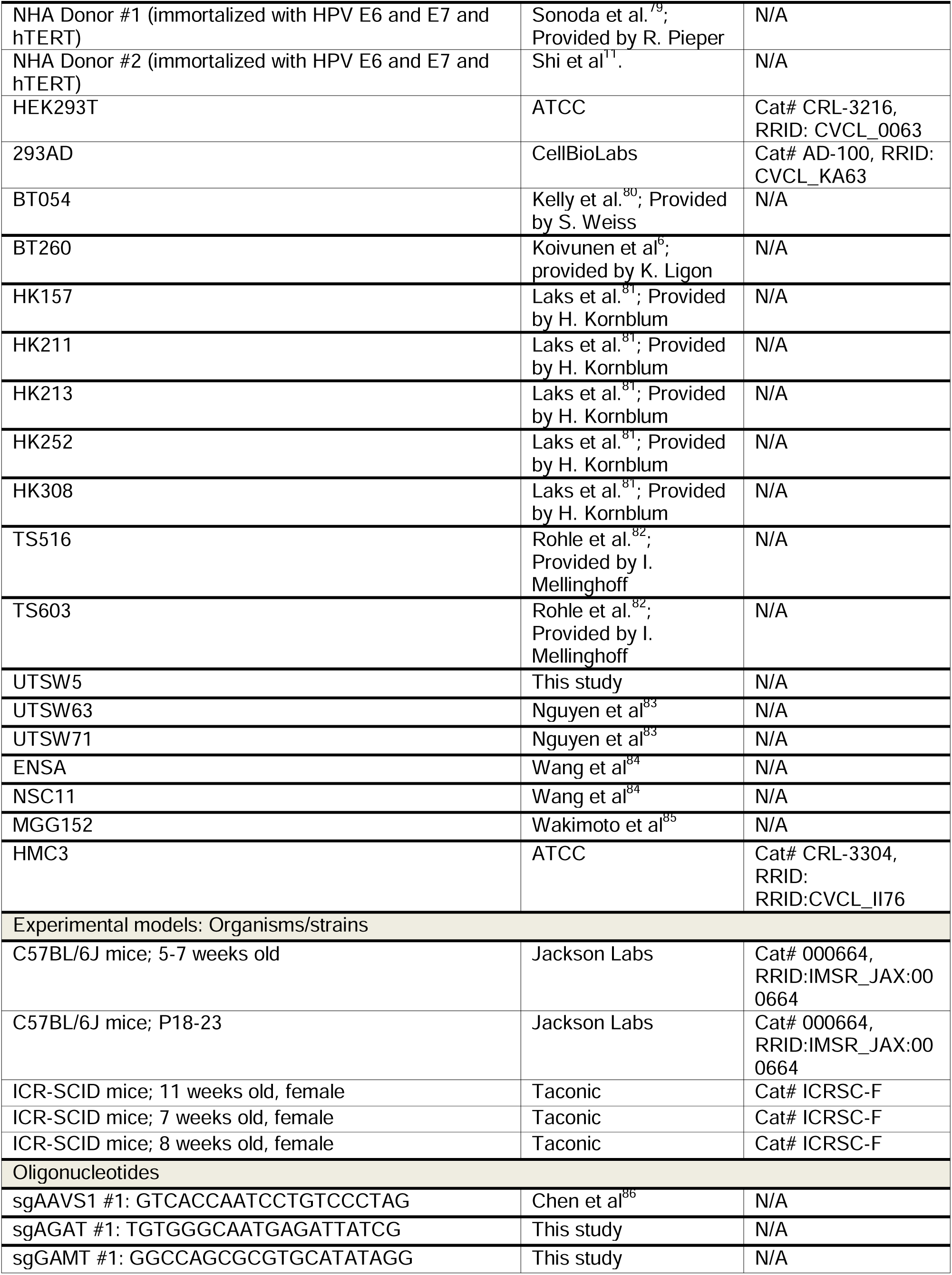
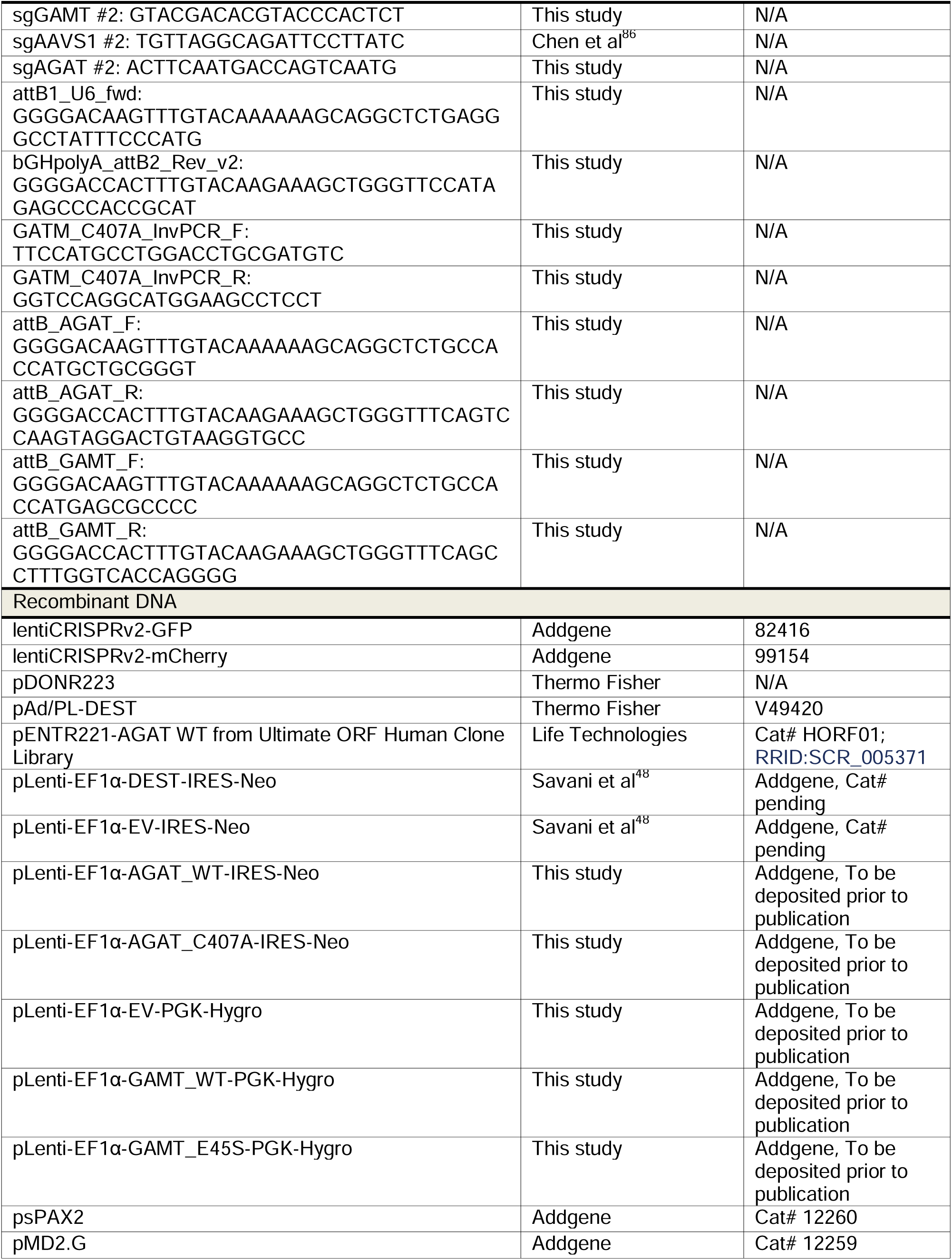
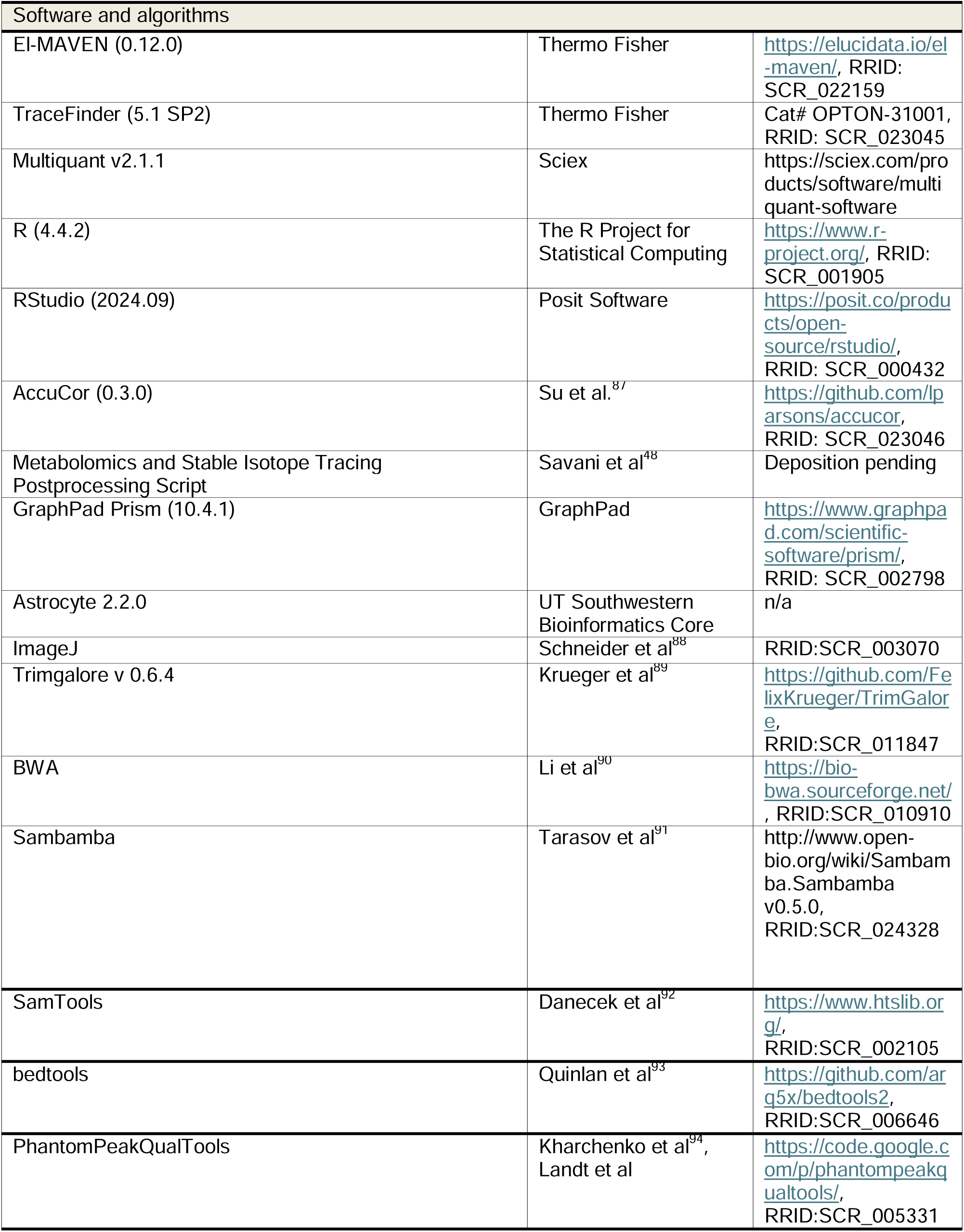
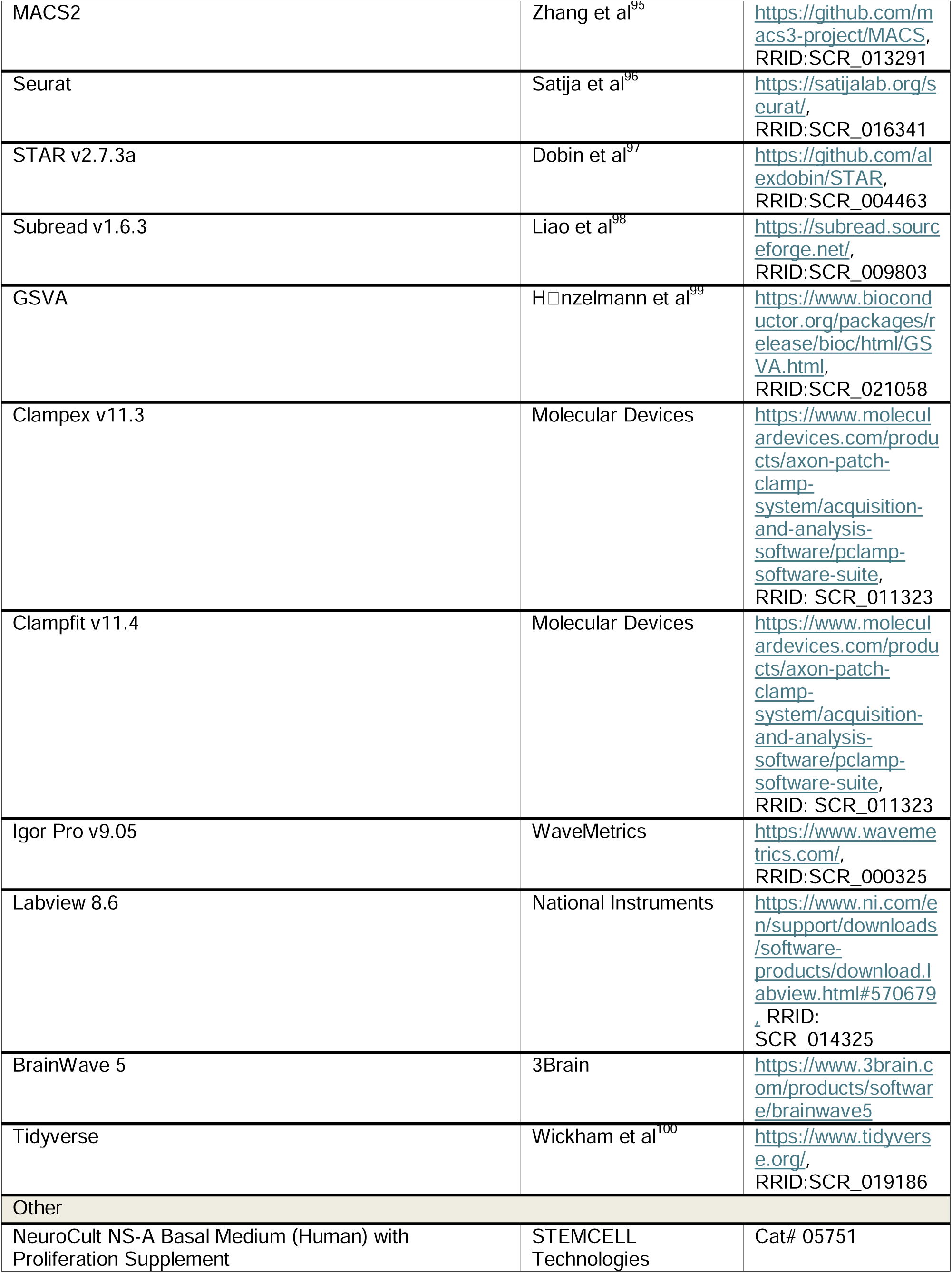
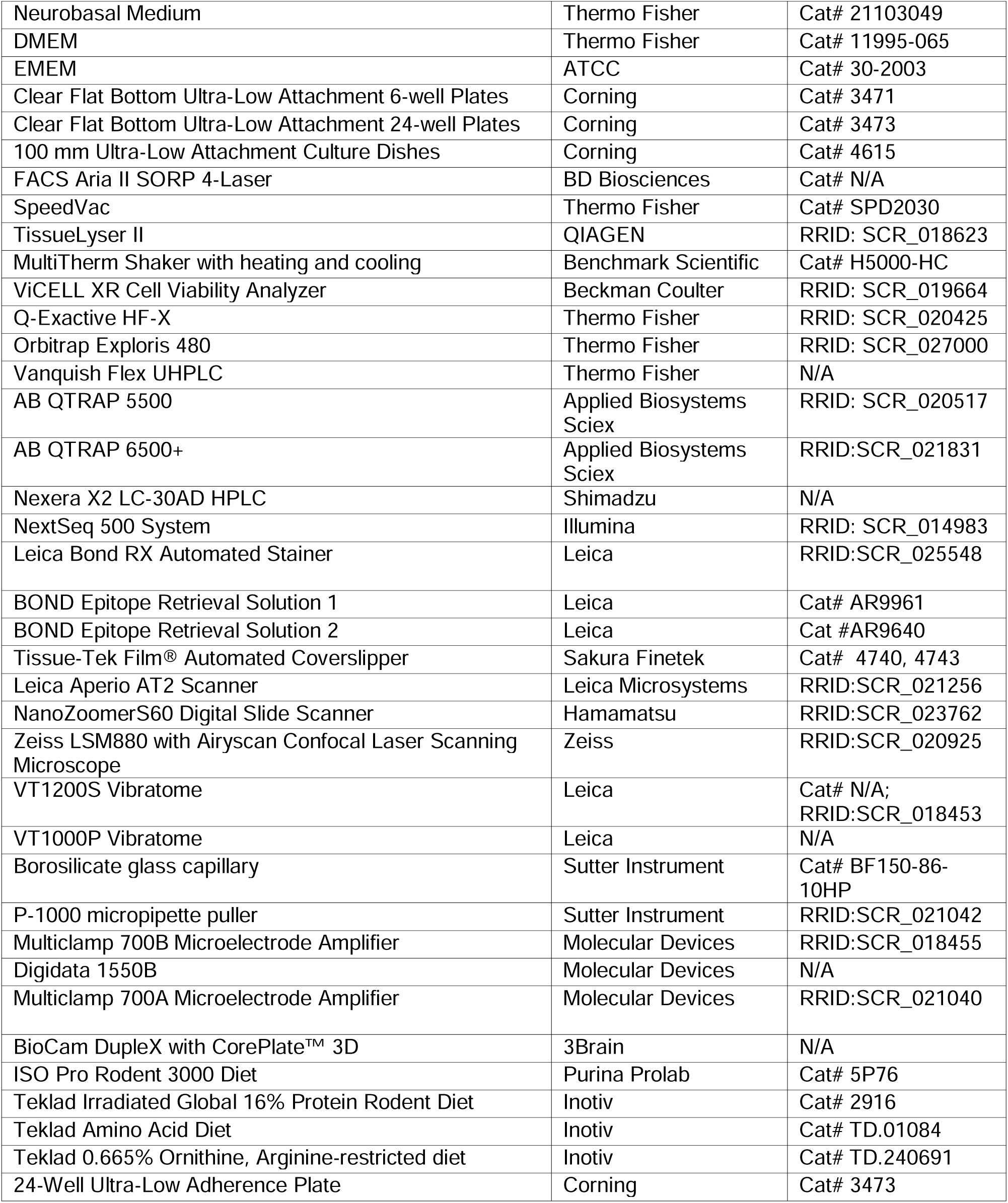

#### Experimental model and study participant details

##### Human Subjects

The study was conducted according to the principles of the Declaration of Helsinki. Patient tissue and blood were collected following ethical and technical guidelines on the use of human samples for biomedical research after informed patient consent under an Institutional Review Board (IRB)-approved protocol. All patient samples were de-identified before processing. All patient samples and organoid explants were diagnosed and graded according to the 2021 WHO Classification of Tumors of the Central Nervous System (CNS), 5th edition^1^. Patient samples for the Discovery Cohort were collected under a protocol approved by the IRB of the University of Texas Southwestern Medical Center (STU 022011-070). Patient samples for Validation Cohort #1 were collected under a protocol approved by the IRB of Northwestern University (17-048-CDH). Patient samples for the Validation Cohort #2 are part of a collection of specimens associated with a study supported by the NCI CPTAC group and are described in a manuscript in preparation. Patient samples used to generate organoid explants were collected under a protocol approved by the IRB of the University of Pittsburgh (STUDY19080321). Information related to primary samples can be found in Tables S1 and S3.

##### Cell lines

NHA Donor #1 cells (non-malignant human astrocytes, immortalized with human papillomavirus E6, E7, and hTERT; sex unknown)^79^ were obtained from R. Pieper at the University of California in San Francisco. NHA Donor #2 cells were generated from commercially available astrocytes (Lonza CC-3187) as described previously^11^. HEK293T cells (female, ATCC CRL-3216) and 293AD cells (female, CellBioLabs AD-100) were obtained commercially. BT054^80^ (female) were provided by S. Weiss at the University of Calgary. BT260^6^ cells (sex unknown) were provided by K. Ligon at Dana Farber Cancer Institute. HK157 (female), HK211 (female), HK213 (male), HK252 (male), and HK308 (female) cells were provided by H. Kornblum at the University of California in Los Angeles^81^. TS516 (sex unknown, RRID: CVCL_A5HY) and TS603 (sex unknown, RRID: CVCL_A5HW) were provided by I. Melinghoff at Memorial Sloan-Kettering Cancer Center^82^. ENSA (sex unknown) and NSC11 (sex unknown) cells were provided by J. Rich at University of North Carolina-Chapel Hill^84^. MGG152 (male) cells were provided by D. Cahill at MGH^85^. UTSW63 (male) and UTSW71 (male) were generated as previously described^83^. HMC3 cells (Sex unknown, ATCC CRL-3304, RRID: CVCL_II76) were acquired commercially.

UTSW5 GBM cells (female) were generated as described previously^83^. Briefly, fresh tumor tissue from the operating room was suspended in ice cold Hibernate A (BrainBits HA) with 100 U/mL and 100 μg/mL, respectively, of penicillin/streptomycin (Thermo Fisher 15-140-148) for transfer on ice to the laboratory, then transferred to RBC lysis buffer (ThermoFisher 00433357) for 10 minute incubation at room temperature before washing in Hibernate A supplemented with 2mM Glutamax (ThermoFisher 35050061), penicillin/streptomycin (100 U/mL and 100 μg/mL, respectively; Thermo Fisher 15-140-148), and Amphotericin B (0.25 μg/mL, Gemini Bio-Products 400104). Following washing, tissue was dissociated using the Brain Tumor Dissociation Kit (Miltenyi Biotec) per manufacturer instructions, and single-cell suspensions were cultured in NeuroCult NS-A Basal Medium (Human) with 1x Proliferation Supplement (STEMCELL Technologies 05751), supplemented with 20 ng/mL EGF (GoldBio 1150-04); 20 ng/mL bFGF (GoldBio 1140-02); 2 µg/mL heparin (STEMCELL Technologies 07980); 100 U/mL and 100 μg/mL, respectively, of penicillin/streptomycin (Thermo Fisher 15-140-148); 250 ng/mL amphotericin B (GeminiBio 400104); and 0.25 µg/mL Plasmocin (InvivoGen ant-mpp) on ultra-low adherence plates (6-well plates: Corning 3471, 10 cm dishes: Corning 4615) in 5% CO2 and at ambient oxygen at 37°C.

All cell lines were routinely evaluated for mycoplasma contamination with the e-Myco Mycoplasma PCR Detection Kit (Bulldog Bio 2523348), e-Myco PLUS Mycoplasma PCR Detection Kit (Bulldog Bio 25233), or MycoAlert Mycoplasma Detection Kit (Lonza LT07-318), per manufacturer’s instructions, and routinely tested negative throughout the course of this study. As reference short term tandem repeat profiles have not been established for these lines, no cell line authentication was performed. Sex and source of each line is stated above or listed as unknown if unreported in the original publication describing its derivation.

##### Cell Culture

BT054, BT260, HK157, HK211, HK213, HK252, HK308, TS516, TS603, UTSW5, UTSW63, and UTSW71 human GSCs were cultured in NeuroCult NS-A Basal Medium (Human) with 1x Proliferation Supplement (STEMCELL Technologies 05751), supplemented with 20 ng/mL EGF (GoldBio 1150-04); 20 ng/mL bFGF (GoldBio 1140-02); 2 µg/mL heparin (STEMCELL Technologies 07980); 100 U/mL and 100 μg/mL, respectively, of penicillin/streptomycin (Thermo Fisher 15-140-148); 250 ng/mL amphotericin B (GeminiBio 400104); and 0.25 µg/mL Plasmocin (InvivoGen ant-mpp) on ultra-low adherence plates (6-well plates: Corning 3471, 10 cm dishes: Corning 4615) in 5% CO2 and at ambient oxygen at 37°C.

ENSA, NSC11, and MGG152 cells were cultured in Neurobasal Medium (Thermo Fisher 21103049) supplemented with 3mM glutamine (Thermo Fisher 25030081); 1× B27 supplement (Thermo Fisher 17504044); 0.25× N2 supplement (Thermo Fisher 17502048); 20 ng/mL EGF (GoldBio 1150-04); 20 ng/mL bFGF (GoldBio 1140-02); 2 µg/mL heparin (STEMCELL Technologies 07980); 50 U/mL and 50 μg/mL, respectively, of penicillin/streptomycin (Thermo Fisher 15-140-148); 125 ng/mL amphotericin B (GeminiBio 400104); and 0.25 µg/mL Plasmocin (InvivoGen ant-mpp) on ultra-low adherence plates (6-well plates: Corning 3471, 10 cm dishes: Corning 4615) in 5% CO2 and at ambient oxygen at 37°C.

NHA Donor #1, NHA Donor #2, HMC3, HEK293T, and 293AD were cultured in DMEM (Thermo Fisher 11995-065) supplemented with 10% fetal bovine serum (FBS; GeminiBio 100-106) and 100 U/mL and 100 μg/mL, respectively, of penicillin/streptomycin (Thermo Fisher 15-140-148) on tissue culture-treated plates in 5% CO2 and at ambient oxygen at 37°C. HMC3 microglia were cultured in EMEM (ATCC 30-2003) supplemented with 10% fetal bovine serum (FBS; GeminiBio 100-106) and 100 U/mL and 100 μg/mL, respectively, of penicillin/streptomycin (Thermo Fisher 15-140-148) on tissue culture-treated plates in 5% CO2 and at ambient oxygen at 37°C.

##### Animals

All care and treatment of experimental animals were carried out in strict accordance with Good Animal Practice as defined by the US Office of Laboratory Animal Welfare and approved by the University of Pittsburgh Medical Center (protocols 25066832 and 25087128) and UT Southwestern Medical Center (protocols 2019-102795 and 2015-101252) Institutional Animal Care and Use Committee. Animal welfare assessments were carried out daily during treatment periods. Animals were housed in a pathogen-free environment between 20-26°C and at 30-70% humidity, with a 12 hour:12 hour light:dark cycle. ICR-SCID (Taconic) mice were obtained from Taconic at 6-7 weeks of age. C57BL/6J mice were obtained from Jackson Laboratories. Mice were housed together (2-5 mice of the same sex per cage) and provided free access to water and chow diet (Purina ISO Pro Rodent 3000 “5P76” or Teklad 2916).

##### Generation of patient-derived xenograft (PDX) mouse models

TS516 orthotopic xenografts for experiments assessing neuronal network activity in tumor-bearing and non-tumor-bearing brain hemispheres were established by intracranial injection of 5,000 cells into 11-week-old female ICR SCID mice (IcrTac:ICR-*Prkdc^scid^*, Taconic ICRSC). TS516 orthotopic xenografts for xenograft studies of AGAT expression were established by intracranial injection of 1,000 TS516 cells into 8-week-old female ICR SCID mice (Taconic ICRSC). TS516 orthotopic xenografts for xenograft studies of dietary ornithine supplementation and arginine restriction were established by intracranial injection of 1,000 TS516 cells into 7-week-old female ICR SCID mice (Taconic ICRSC). Mice were anesthetized with isoflurane and immobilized using a stereotactic frame. An incision was made to expose the skull surface, and a hole was drilled into the skull. Cells suspended in 5 µL cell culture medium were injected into the brain through the hole using a 5 μL syringe (Hamilton) at 0.5-1 mm anterior and 2 mm lateral to the bregma and a depth of 3 mm from the brain surface. The skin was closed with surgical clips and analgesia was administered. Mice receiving experimental diets received either Teklad amino acid diet (Inotiv TD.01084) or diet supplemented with 0.665% ornithine and arginine-free (Inotiv TD.240691). Survival analyses were performed by researchers who were not blinded to the treatment arms or genotypes of the mice. Mice were euthanized when they displayed neurological symptoms or became moribund.

#### Method Details

##### Chemicals

Where indicated, cells or brain slices were treated with gabazine (EMD Millipore SR95531), GAA (Millipore Sigma G11608, or TCI G0167), creatine monohydrate (Thermo Fisher B25009.22), CLP-257 (Bio-Techne 5242/10), 10 µM 3-(2-carboxypiperazin-4-yl)propyl-1-phosphonic acid (CPP) (Hello Bio HB0036), 20 µM (6,7-dinitroquinoxaline-2,3-dione (DNQX) (Tocris 2312), 100 µM picrotoxin (Hello Bio HB0506), and/or 2 µM CGP 52432 (Tocris 1246).

##### Vectors

lentiCRISPRv2-GFP (Addgene 82416) and lentiCRISPRv2 mCherry (Addgene 99154) were digested with FastDigest Esp3I (Thermo Fisher FD0454) or BsmBIv2 (New England Biolabs R0739). sgRNAs targeting the AAVS1 safe harbor locus^86^ (sg #1: GTCACCAATCCTGTCCCTAG), AGAT (sg #1: TGTGGGCAATGAGATTATCG), or GAMT (sg #1: GGCCAGCGCGTGCATATAGG; sg #2: GTACGACACGTACCCACTCT) were ligated into lentiCRISPRv2-GFP to generate a construct expressing Cas9 and the desired sgRNA. sgRNAs targeting the AAVS1 safe harbor locus (sg #2: TGTTAGGCAGATTCCTTATC) or AGAT (sg #2: ACTTCAATGACCAGTCAATG) were ligated into lentiCRISPRv2-mCherry to generate a construct expressing Cas9 and the desired sgRNA. Reaction mixtures were transformed into XL10-Gold Ultracompetent Cells (Agilent 200315) and validated by whole plasmid sequencing (Plasmidsaurus).

CRISPR/Cas9 and fluorophore expression cassettes for lentiCRISPRv2-GFP-sgAAVS1 #1, lentiCRISPRv2-GFP-sgAGAT #1, lentiCRISPRv2-mCherry-sgAAVS1 #2, and lentiCRISPRv2-mCherry-sgAGAT #2 were amplified from these constructs and appended with 5’ attB1 and 3’ attB2 sites using the following primers:

*attB1_U6_Fwd_v2*: GGGGACAAGTTTGTACAAAAAAGCAGGCTCTGAGGGCCTATTTCCCATG

*bGHpolyA_attB2_Rev_v2*: GGGGACCACTTTGTACAAGAAAGCTGGGTTCCATAGAGCCCACCGCAT

PCR product was gel-purified with the QIAquick Gel Extraction Kit (Qiagen 28706), purified with the Monarch PCR & DNA Cleanup Kit (New England Biolabs T1030) and cloned into the Gateway vector pDONR223 by BP reaction (Thermo Fisher 11789100) overnight. The resultant products were transformed into HB101 Competent Cells (Promega L2015), and identity of the final products were validated by whole-plasmid sequencing (Plasmidsaurus).

pENTR223 plasmids containing CRISPR/Cas9 expression cassettes for GFP-sgAAVS1 #1, GFP-sgAGAT #1, mCherry-sgAAVS1 #2, and mCherry-sgAGAT #2 were cloned into the Gateway vector pAd/PL-DEST (Thermo Fisher V49420) by LR reaction (Thermo Fisher 11791020) overnight. Resultant pAdCRISPR-GFP-sgAAVS1 #1, pAdCRISPR-GFP-sgAGAT #1, pAdCRISPR-mCherry-sgAAVS1 #2 and pAdCRISPR-mCherry-sgAGAT #2 vectors for adenoviral transduction of mammalian cells were transformed into HB101 Competent Cells (Promega L2015) and validated by whole plasmid sequencing (Plasmidsaurus).

AGAT (RefSeq NM_001482.2) WT cDNA was generated from the Ultimate ORF Human Clone Library (Life Technologies HORF01) in a pENTR221 Gateway vector. The C407A point mutation was generated by inverse PCR of pENTR221-AGAT using CloneAmp HiFi PCR Master Mix (Takara 639298) and the following primers:

*GATM_C407A_InvPCR_F*: TTCCATGCCTGGACCTGCGATGTC

*GATM_C407A_InvPCR_R:* GGTCCAGGCATGGAAGCCTCCT

The amplified inverse PCR product was then gel-purified with the QIAquick Gel Extraction Kit (Qiagen 28706), PCR purified with the Monarch PCR & DNA Cleanup Kit (New England Biolabs T1030) and assembled using the In-Fusion HD Cloning Kit (Takara Bio 102518). The assembled product was then transformed into XL10-Gold Ultracompetent Cells (Agilent 200315). Final pENTR221-AGAT_C407A product was validated by whole plasmid sequencing (Plasmidsaurus).

5’ attB1 and 3’ attB2 sites were appended to AGAT WT and AGAT C407A cDNA using Q5 Hot Start High-Fidelity 2X Master Mix polymerase (New England Biolabs M0494) and the following primers:

*attB_AGAT_F:* GGGGACAAGTTTGTACAAAAAAGCAGGCTCTGCCACCATGCTGCGGGT

*attB_AGAT_R:* GGGGACCACTTTGTACAAGAAAGCTGGGTTTCAGTCCAAGTAGGACTGTAAG GTGCC

PCR product was gel-purified with the QIAquick Gel Extraction Kit (Qiagen 28706), purified with the Monarch PCR & DNA Cleanup Kit (New England Biolabs T1030) and cloned into the Gateway vector pDONR223 by BP reaction (Thermo Fisher 11789100) overnight. The resultant products were transformed into HB101 Competent Cells (Promega L2015). pENTR223-AGAT_WT and pENTR223-AGAT_C407A were validated by whole plasmid sequencing (Plasmidsaurus).

GAMT WT and GAMT E45S cDNA were synthesized as gBlocks by Twist Biosciences. attB sites were added and sequences amplified using Q5 Hot Start High-Fidelity 2X Master Mix polymerase (New England Biolabs M0494) and the following primers:

*attB_GAMT_F*: GGGGACAAGTTTGTACAAAAAAGCAGGCTCTGCCACCATGAGCGCCCC

*attB_GAMT_R:* GGGGACCACTTTGTACAAGAAAGCTGGGTTTCAGCCTTTGGTCACCAGGGG

PCR product was gel-purified with the QIAquick Gel Extraction Kit (Qiagen 28706), PCR purified with the Monarch PCR & DNA Cleanup Kit (New England Biolabs T1030) and cloned into the Gateway vector pDONR223 by BP reaction (Thermo Fisher 11789100) overnight to generate pENTR223-GAMT_WT and pENTR223-GAMT_E45S plasmids. The resultant products were transformed into HB101 Competent Cells (Promega L2015). pENTR223-GAMT_WT and pENTR223-GAMT _E45S were validated by whole plasmid sequencing (Plasmidsaurus).

pENTR223-EV, pENTR223-AGAT_WT, and pENTR223-AGAT_C407A were cloned into pLenti-EF1α-DEST-IRES-Neo^48^ (Addgene deposition pending) by Gateway LR reaction (Thermo Fisher 11791020) overnight. The lentiviral vectors pLenti-EF1α-EV-IRES-Neo, pLenti-EF1α-AGAT_WT-IRES-Neo, and pLenti-EF1α-AGAT_C407A-IRES-Neo were transformed into HB101 Competent Cells (Promega L2015) and validated by whole plasmid sequencing (Plasmidsaurus).

pENTR223-EV, pENTR223-GAMT_WT, and pENTR223-GAMT_E45S were cloned into pLenti-EF1α-DEST-PGK-Hygro by Gateway LR reaction (Thermo Fisher 11791020) overnight. The lentiviral vectors pLenti-EF1α-EV-PGK-Hygro, pLenti-EF1α-GAMT_WT-PGK-Hygro, and pLenti-EF1α-GAMT_E45S-PGK-Hygro were transformed into HB101 Competent Cells (Promega L2015) and validated by whole plasmid sequencing (Plasmidsaurus).

##### Transfection and viral transduction

Lentiviral particles were produced by transfection of HEK293T cells with expression vectors and packaging plasmids psPAX2 (Addgene 12260, gift of Didier Trono) and pMD2.G (Addgene 12259, gift of Didier Trono) in a ratio of 4:3:1 using TransIT-LT1 transfection reagent (Mirus Bio MIR2 B-304). Media was discarded and replaced at 24 hours post-transfection, then virus-containing media was collected 48 and 72 hours post-transfection and passed through a 0.45 μm filter (Corning 431220). 1 volume LentiX Concentrator (Takara 631232) was added per 3 volumes clarified supernatant. The mixture of viral supernatant and LentiX Concentrator was incubated at 4°C overnight, then centrifuged at 1500 × g for 45 minutes. After centrifugation, supernatant was discarded, and the remaining pellet was resuspended in original viral supernatant volume of DMEM supplemented with 10% fetal bovine serum (FBS; GeminiBio 100-106) and 100 U/mL and 100 μg/mL, respectively, of penicillin/streptomycin (Thermo Fisher 15-140-148).

To generate adenovirus, pAdCRISPR constructs were linearized by PacI (New England Biolabs R0547S) digestion at 37°C overnight. 293Ad cells were subsequently transfected with 2.5 µg linear pAdCRISPR constructs using TransIT-LT1 transfection reagent (Mirus Bio MIR2 B-304). Media was changed at 24 hours after transfection, and cells were expanded to a 10 cm dish and incubated for 10-14 days with regular media changes. 10-14 days following transfection, 293Ad cells and media were harvested and centrifuged at 3739 × g for 5 minutes. Supernatant was discarded and the pellet was resuspended in 1 mL media. Resuspended pellet was subjected to 3 freeze-thaw cycles with 30 minutes at -80°C followed by 15 minutes at 37°C. After final thaw, suspensions were centrifuged at 2093 × g for 15 minutes to generate crude adenovirus. Presence of adenovirus capsid in crude adenovirus prep was confirmed using Adeno-X GoStix (Takara 632270). Crude adenovirus was then amplified 2-3 times by inoculating 293Ad cells with crude virus and harvesting as above after 2-4 days. Amplified virus was purified using Virabind Adenovirus Miniprep Kit (Cell Biolabs VPK-099) according to manufacturer’s instructions.

NHA Donor #1 cells were plated at a density of 1.5 × 10^5^ cells per well in 2 mL DMEM in a six well plate and allowed to adhere overnight. The next day, 3 uL polybrene (8 µg/mL, MedChemExpress HY-112735) was added to each well along with 1 mL viral pLenti-EF1a-EV-IRES-Neo, pLenti-EF1a-AGAT_WT-IRES-Neo, or pLenti-EF1a-AGAT_C407A-IRES-Neo supernatant. Plates were centrifuged at 4,000 × g for 30 minutes at room temperature, then incubated overnight. The following day, cells were expanded and replated in a 10 cm dish. After initial selection at 1500 µg/mL G418 (GoldBio G-418), stable cell lines were maintained in 600 µg/mL G418.

NHA Donor #1 EV, AGAT WT, and AGAT C407A cells were plated at a density of 1.5 × 10^5^ cells per well in 2 mL DMEM without G418 in a six well plate and allowed to adhere overnight. The next day, 3 uL polybrene (8 µg/mL, MedChemExpress HY-112735) was added to each well along with 1 mL virus for pLenti-EF1a-EV-PGK-hygro, pLenti-EF1a-GAMT_WT-PGK-Hygro, or pLenti-EF1a-GAMT_E45S-PGK-Hygro; or 1 mL lentiCRISPRv2-GFP-sgAAVS1 #1, lentiCRISPRv2-GFP-sgGAMT #1, or lentiCRISPRv2-GFP-sgGAMT #2 lentivirus. Plates were centrifuged at 4,000 × g for 30 minutes at room temperature, then incubated overnight. The following day, cells were expanded and replated in a 10 cm dish. For cells transduced with pLenti-EF1a-EV-PGK-hygro, pLenti-EF1a-GAMT_WT-PGK-Hygro, or pLenti-EF1a-GAMT_E45S-PGK-Hygro, following initial selection at 250 µg/mL hygromycin (GoldBio H-270-EZ50), stable cell lines were maintained in 100 µg/mL hygromycin. Cells transduced with lentiCRISPRv2-GFP constructs were sorted for GFP positivity 14 days post-transduction using a FACS Aria II SORP Four-Laser instrument (BD Biosciences) in the Children’s Research Institute Flow Cytometry Core.

For adenoviral transduction, TS516 cells were plated at 1 × 10^7^ cells per dish in 10 mL NeuroCult NS-A Basal Medium (Human) and treated with AdCRISPR GFP sgAAVS1 #1 or sgAGAT #1 adenovirus at a concentration empirically determined to yield 30-50% transduction efficiency for each viral preparation. At 16-18 hours, media was changed to fresh NeuroCult NS-A Basal Medium (Human) and cells were split out to 2 dishes each. At 7 days post-transduction, cells were sorted for GFP positivity using a FACS Aria II SORP Four-Laser instrument (BD Biosciences) in the Children’s Research Institute Flow Cytometry Core. Sorted GFP^+^ cells were transduced with AdCRISPR mCherry sgAAVS1 #2 or sgAGAT #2 adenovirus 14 days following initial sort as previously done with GFP-containing virus, then sorted 7 days following secondary transduction using a FACS Aria II SORP Four-Laser instrument (BD Biosciences) in the Children’s Research Institute Flow Cytometry Core. After two weeks in culture, sorted mCherry^+^ cells were sorted for GFP and mCherry dual-negative populations to establish stably-transduced AGAT WT and AGAT KO lines.

##### Preparation of HPLM and tracer HPLM

HPLM was prepared as previously described^49^. Tracer-containing HPLM library was as previously described^48^, with the addition of a medium containing ^15^N_3_-creatine (Cambridge Isotope Laboratories NLM-9218). Tracer-containing HPLM used in nitrogen metabolism profiling, profiling validation, and ^15^N_4_-arginine tracing experiments were generated identically to the unlabeled version, with labeled metabolites replacing their unlabeled counterparts at equimolar concentrations. HPLM for ^15^N_4_-arginine tracing in TS516 under ornithine-supplemented conditions was generated without arginine or ornithine, then 400 µM ^15^N_4-_arginine (Cambridge Isotope Laboratories NLM-396) and either 0 µM or 150 µM ornithine (Millipore Sigma O2375) were added. Tracer HPLM was assembled by mixing thawed frozen or freshly-prepared stocks of unlabeled HPLM pools, replacing one pool per tracing condition with a version containing an individual ^15^N tracer. Media was adjusted to pH 7.4 using NaOH or HCl, then diluted to the final volume with ultrapure water, sterile filtered with a 0.22 µm PES filter (Millipore Sigma SCGP00525, Corning 431153, Corning 431097, or Corning 431098), and supplemented as described below. To prevent glutamate toxicity in culture, HPLM was prepared without glutamate for experiments in GSCs, NSCs, and immortalized astrocyte cells, except for conditions in which glutamate was traced.

For experiments in GSCs and NSCs, HPLM was supplemented with 1 × B27 supplement (Thermo Fisher 17504044); 0.25 × N2 supplement (Thermo Fisher 17502048); 20 ng/mL EGF (GoldBio 1150-04); 20 ng/mL bFGF (GoldBio 1140-02); 2 µg/mL heparin (STEMCELL Technologies 07980); 50 U/mL and 50 μg/mL, respectively, penicillin/streptomycin (Thermo Fisher 15-140-148); 125 ng/mL amphotericin B (GeminiBio 400104); and 0.25 µg/mL Plasmocin (InvivoGen ant-mpp). For experiments in differentiated cells, HPLM was supplemented with 10% dialyzed FBS (GeminiBio 100-108) and 100 U/mL and 100 μg/mL, respectively, of penicillin/streptomycin (Thermo Fisher 15140148).

##### Nitrogen metabolism profiling platform

Nitrogen metabolism profiling was conducted as described previously^48^. Briefly, NHA Donor #1 cells were plated in 6-well plates (2.5 × 10^4^ cells per well) in 2 mL DMEM prepared as described above. TS516 cells were plated in 6-well ultra-low adherence plates (2 × 10^5^ cells per well) in 2 mL NeuroCult NS-A Basal Medium (Human) prepared as described above. After 24 hours, 2 mL unlabeled HPLM prepared as described above was added to produce a mixture of 50% native medium and 50% HPLM. 24 hours later, media was changed to 100% unlabeled HPLM (10 mL for NHA Donor #1 cells and 4.5 mL for TS516 cells). 24 hours later, media was changed to fresh 100% HPLM with one nitrogen-containing metabolite per sample exchanged for an equimolar amount of its ^15^N-labeled counterpart (10 mL for NHA Donor #1 cells, 4.5 mL for TS516 cells). After 18 hours, samples were harvested and prepared for LC-MS analysis as described below.

##### Creatine synthesis pathway tracing in cells

For tracing of creatine synthesis pathway substrates, differentiated cells (NHA Donor #1 and #2) were plated at 2.5 × 10^4^ cells per well for NHA Donor #1 and 1 × 10^5^ cells per well for NHA Donor #2 in 2 mL DMEM. GSCs with exception of MGG152 were plated at 2 × 10^5^ cells per well in 2 mL NeuroCult NS-A Basal Medium (Human) prepared as described above in 6-well ultra-low adherence plates. NSCs and MGG152 were plated at 2 × 10^5^ cells per well in 2 mL NeuroBasal prepared as described above in 6-well ultra-low adherence plates. 24 hours after plating, 2 mL unlabeled HPLM prepared as described above was added to produce a 1:1 mixture of HPLM and native medium. 24 hours later, media was changed to 10 mL 100% HPLM for NHA Donor #1 and #2 or 4.5 mL HPLM for GSCs or NSCs. 24 hours later, media was changed to 10 or 4.5 mL fresh HPLM in which the unlabeled metabolite was exchanged for an equimolar concentration of ^15^N_4_-arginine (Cambridge Isotope Laboratories NLM-396), ^15^N_3_-creatine (Cambridge Isotope Laboratories NLM-9218), ^15^N-glycine (Cambridge Isotope Laboratories NLM-202), or ^15^N-serine (Cambridge Isotope Laboratories NLM-2036). Cells were incubated with tracer for 18 hours, then harvested and prepared for LC-MS as described below. For each cell line, counts and diameter were quantified from parallel cell cultures using a Beckman Coulter Vi-CELL XR cell viability analyzer (RRID: SCR_019664) at the time of sample harvest.

##### Intracellular GAA content in cultured cells

Differentiated cells (NHA Donor #1 and #2) were plated at 5 × 10^4^ cells per well for NHA Donor #1 and 2 x 10^5^ cells per well for NHA Donor #2 in 2 mL DMEM. GSCs with exception of MGG152 were plated at 4 × 10^5^ cells per well in 2 mL NeuroCult NS-A Basal Medium (Human) prepared as described above in 6-well ultra-low adherence plates. NSCs and MGG152 were plated at 4 × 10^5^ cells per well in 2 mL NeuroBasal medium prepared as described above in 6-well ultra-low adherence plates. 24 hours after plating, 2 mL unlabeled HPLM prepared as described above was added to produce a 1:1 mixture of HPLM and native medium. 24 hours later, media was changed to 10 mL 100% HPLM for NHA Donor #1 and #2 or 4.5 mL HPLM for GSCs or NSCs. 24 hours later, media was changed to 2 mL fresh HPLM. 24 hours later, media was changed to 2 mL fresh HPLM per well. 24 hours later, cells were collected, washed in Optima saline, and snap frozen. For each cell line, cell counts and cell diameter were quantified from parallel cell cultures using a Beckman Coulter Vi-CELL XR cell viability analyzer (RRID: SCR_019664) at the time of sample harvest.

##### Secretion of GAA in cell culture

Differentiated cells (NHA Donor #1 and #2) were plated at 5 × 10^4^ cells per well for NHA Donor #1 and 2 × 10^5^ cells per well for NHA Donor #2 in 2 mL DMEM in 6-well plates. HMC3 microglia were plated at 5 × 10^4^ cells per well in 2 mL EMEM in 6-well plates. GSCs with exception of MGG152 were plated at 4 × 10^5^ cells per well in 2 mL NeuroCult NS-A Basal Medium (Human) prepared as described above in 6-well ultra-low adherence plates. NSCs and MGG152 were plated at 4 × 10^5^ cells per well in 2 mL NeuroBasal prepared as described above in 6-well ultra-low adherence plates. 24 hours after plating, 2 mL unlabeled HPLM prepared as described above was added to produce a 1:1 mixture of HPLM and native medium. 24 hours later, media was changed to 10 mL 100% HPLM for NHA Donor #1 and #2 or 4.5 mL HPLM for GSCs or NSCs. 24 hours later, media for all cells was changed to 2 mL fresh HPLM. 48 hours later, media samples were collected from wells, spun at 4,000 × g for 3 minutes, 100 µL transferred to a fresh microcentrifuge tube, and snap frozen in liquid nitrogen. For each cell line, cell counts and cell diameter were quantified from parallel cell cultures using a Beckman Coulter Vi-CELL XR cell viability analyzer (RRID: SCR_019664) at the time of sample harvest.

##### Preparation of cultured cells for LC-MS

Prior to LC-MS analysis, cell samples were suspended in 1 µL ice-cold 80% acetonitrile (Thermo Fisher A9554) per 1,000 cells. Adherent cells (NHA Donor #1 and NHA Donor #2) were lifted from plates by scraping to facilitate resuspension. Cell samples resuspended in acetonitrile were platform-vortexed for 20 minutes at 4°C and centrifuged for 10 minutes at 17,000-21,100 × g at 4°C. The supernatant was transferred to a fresh tube and again centrifuged for 10 minutes at 17,000-21,100 × g at 4°C. The resultant supernatant was then analyzed by LC-MS. For experiments in which absolute quantification of GAA in cell samples was performed, internal standards 1,2-¹³C, 3-¹ N-guanidinoacetic acid (Cambridge Isotope Laboratories CNLM-8300) or 2,2-D_2_-guanidinoacetic acid (Cambridge Isotope Laboratories DLM-9998) at concentrations of 20, 100, or 200 nM were added. Standard curves for GAA (Millipore Sigma G11608) were created fresh for each submission in neat 80% acetonitrile over a range of 1-10,000 nM. All reagents used were Optima-grade.

##### Preparation of media for LC-MS

Prior to LC-MS analysis, media was diluted 1:10 in ice-cold 80% acetonitrile (Thermo Fisher A9554) and vortexed for 20 minutes at 4°C, then centrifuged for 10 minutes at 17,000-21,100 × g at 4°C. The supernatant was transferred to a fresh tube and again centrifuged for 10 minutes at 17,000-21,100 × g at 4°C. The resultant supernatant was then analyzed by LC-MS. For experiments in which absolute quantification of GAA was performed in media, an internal standard of 1,2-¹³C, 3-¹ N-guanidinoacetic acid (Cambridge Isotope Laboratories CNLM-8300) was added to all samples and standards at concentrations of 20, 100, or 200 nM. Standard curves for GAA (Millipore Sigma G11608) were created fresh for each submission in neat 80% acetonitrile over a range of 1-10,000 nM. All reagents used were Optima-grade.

##### Preparation of tissue for LC-MS

Tissues collected for metabolomic profiling of human brain tumors were obtained in the operating room, suspended in ice-cold Hibernate A (BrainBits HA), and snap frozen. Samples were prepared as previously described^101^, with some modifications. Briefly, samples were thawed, washed in 1 mL ice-cold normal saline, and resuspended in 50 µL ice-cold 80% methanol (Thermo Fisher A456) per mg of tissue. Tissue samples were homogenized in a TissueLyser II (Qiagen 85300, RRID: SCR_018623) at 4°C at an oscillation rate of 25 Hz. 520 µL tert-Butyl methyl ether (Millipore Sigma 650560) and 380 µL water were added to 500 µL of tissue homogenate in a glass vial, then vortexed at room temperature for 1 hour at 1,000 rpm in a thermal mixer (Benchmark Scientific H5000-HC). Samples were centrifuged for 10 minutes at 1,000 × g at 4°C.

For polar metabolomics analysis, 800 µL of the lower aqueous phase was combined with 300 µL of the upper organic phase, avoiding interlayer debris. A SpeedVac (Thermo Fisher SPD2030) was used to dry down polar metabolite extracts, and dried metabolites were then resuspended in 100 µL ice-cold 80% acetonitrile (Thermo Fisher A9554). A mixture of internal standards, composed of 3 µM ^13^C_5_-glutamine (Cambridge Isotope Laboratories CLM-1822), 3 µM ^15^N-valine (Cambridge Isotope Laboratories NLM-316), 3 µM ^15^N-methionine (Cambridge Isotope Laboratories NLM-752), and 3 µM chloramphenicol (Millipore Sigma C0378) was added. Samples were sonicated in a room temperature water bath for 10 minutes, then centrifuged for 10 minutes at 21,100 × g at 4°C. The resultant supernatant was then analyzed by LC-MS.

For lipidomics analysis, 100 µL of upper organic phase was transferred to a new glass vial. The lipid extract was dried using nitrogen to avoid oxidation, and dried lipids were resuspended in 500 µL isopropanol and methanol mixture (65:35, V/V) with 15 µL of Avanti Sph/Cer mix standard (Avanti Research, 330707). Samples were then vortexed at 4°C for 10 minutes, then centrifuged for 10 minutes at 1000 × g at 4°C. The resultant supernatant was then analyzed by LC-MS.

For experiments in which absolute quantification of GAA in human tissue samples was performed, tissues were homogenized in 50 µL ice-cold 80% Optima methanol (Thermo Fisher A456) per mg of tissue using a TissueLyser II (Qiagen 85300, RRID: SCR_018623) with two carbide beads per tube (Qiagen 69997) at an oscillation rate of 25 Hz. Samples were vortexed for 20 minutes at 4°C, then centrifuged for 10 minutes at 17,000-21,100 × g at 4°C. The supernatant was transferred to a fresh tube then centrifuged for 10 minutes at 17,000-21,100 × g at 4°C. 200 or 250 µL of supernatant was transferred to a fresh tube, then dried using a SpeedVac (Thermo Fisher SPD2030). Dried metabolites were resuspended in 7.5 µL ice-cold 80% acetonitrile (Thermo Fisher A9554) per mg tissue, vortexed for 20 minutes at 4°C, and centrifuged for 10 minutes at 17,000-21,100 × g at 4°C. The resultant supernatant was then diluted 1:10 and analyzed by LC-MS. For experiments in which absolute quantitation of GAA was performed in tissues, 1,2-¹³C, 3-¹ N-guanidinoacetic acid (Cambridge Isotope Laboratories CNLM-8300) was added as an internal standard to all samples and standards. Standard curves for GAA (Millipore Sigma G11608) were created fresh for each submission in neat 80% acetonitrile over a range of 1-10,000 nM. All reagents used were Optima-grade.

Extraction of TS516 AGAT WT and AGAT KO xenograft tissues collected for quantification of GAA and creatine was performed using pre-chilled 80% methanol (Thermo Fisher A456). A volume of 80% MeOH equivalent to 50 µL per mg of tissue was added to each sample tube. Two tungsten carbide beads (Qiagen 69997) were added to each tube for homogenization. Tissue homogenization was carried out using a TissueLyser set to 25 Hz for 1-minute cycles. Following homogenization, samples were vortexed for 20 minutes at 4 °C. Samples were centrifuged twice at maximum speed (∼13,000–15,000 × g) for 10 minutes at 4 °C. The resulting supernatant was transferred and analyzed by LC-MS/MS and internal standards for both GAA (Guanidineacetic-2,2-d2 Acid, CDN Isotopes D-6320) and creatine [Creatine-d5 H2O (N-methyl-d3; glycine-2,2-d2), LGC Standards TRC-C781496] were added to final supernatant. A standard curve of creatine (LGC Standards TRC-C781483) and GAA (TRC-G821250) was created for each run with duplicate QCs (low, medium, and high) to ensure the results for each day were accurate. The linear range for GAA and creatine in brain was 1-25,000 ng/mL.

##### Metabolite Quantification by LC-MS

10-20 µL of metabolite extract in acetonitrile was injected and analyzed with a Q-Exactive HF-X (Thermo, RRID: SCR_020425), Orbitrap LUMOS (Thermo, RRID: SCR_020562), or Orbitrap Exploris 480 (Thermo, RRID: SCR_027000) hybrid quadrupole-orbitrap mass spectrometer coupled to a Vanquish Flex UHPLC system (Thermo Fisher), as described previously^102, 103^; or with an AB QTRAP 5500 (Applied Biosystems Sciex) or AB QTRAP 6500+ (Applied Biosystems Sciex) with ESI source coupled to a Nexera X2 LC-30AD HPLC (Shimadzu Corporation). Chromatographic separation of samples acquired on the Q-Exactive, LUMOS, or Exploris Orbitrap instruments was accomplished using a Millipore-Sigma ZIC-pHILIC column using a binary gradient of 10 mM ammonium formate pH 9.8 and acetonitrile. Spectra acquired on Orbitrap instruments were acquired with resolving power of 60,000, 120,000, or 240,000 full width at half maximum (FWHM), and a scan range set to 60-1,050 or 80-1,200 m/z, collecting data in both positive and negative polarities. To improve signal resolution of GAA and its isotopologues, a targeted selected ion monitoring (tSIM) scan event using a 7 Da window around GAA was used in tandem with full scan data acquisition to capture all relevant stable isotope labeling events with an AGC targeted of 1 × 10^5^ ions.

Chromatographic separation of samples acquired on the AB QTRAP 5500 or AB QTRAP 6500+ was accomplished with a SeQuant® ZIC®-pHILIC HPLC column (150 × 2.1 mm, 5 µm, polymeric) or a BEH Z-HILIC VanGuardTM Fit HPLC column (150 × 2.1 mm, 5 µm), using an electrospray ionization (ESI) source in multiple reaction monitoring (MRM) mode. Samples acquired on the AB QTRAP 5500 were resolved using a binary gradient of 10 mM ammonium acetate in water (pH 9.8 adjusted with ammonium water; mobile phase A) and 100% ACN (mobile phase B), with gradient elution as follows: 0-15 minutes, linear gradient 90-30% B, followed by 3 minute wash with 30% B before reconditioning for 6 minutes using 90% B. Samples acquired on the AB QTRAP 6500 were resolved using a binary gradient of 10 mM ammonium acetate in water (pH 9.8 adjusted with ammonium water; mobile phase A) and 90% ACN:10% mobile phase A (mobile phase B), with gradient elution as follows: 0-15 minutes, linear gradient 100-33% B, followed by 3 minute wash with 33% B before reconditioning for 6 minutes using 90% B. A flow rate of 0.25 mL/minute was used with injection volume 10-20 µL. MRM data was acquired with Analyst 1.6.3 software (SCIEX).

For samples acquired on the AB QTRAP 5500 in which isotopologue species of GAA and creatine pathway-associated metabolites were quantified, the MRMs used for each metabolite were as follows: For GAA, M+0 (Q1/Q3: 118/76, CE: 16), M+1 (Q1/Q3: 119/76 and 119/77, CE: 16), M+2 (Q1/Q3: 120/76, 120/77, 120/78; CE: 16), M+3 (Q1/Q3: 121/76, 121/77, 121/78, CE: 16), internal standard M+3 (Q1/Q3: 121/79, CE: 16); for arginine, M+0 (Q1/Q3: 175/70, CE: 25), M+1 (Q1/Q3: 176/70, 176/71; CE: 25), M+2 (Q1/Q3: 177/70, 177/71, 177/72; CE: 25), M+3 (Q1/Q3: 178/70, 178/71, 178/72, 178/73; CE: 25), M+4 (Q1/Q3: 179/70, 179/71, 179/72, 179/73, 179/74; CE: 25); for ornithine, M+0 (Q1/Q3: 133/70, CE: 25), M+1 (Q1/Q3: 134/70, 134/71; CE: 25), M+2 (Q1/Q3: 135/70, 135/71, 135/72; CE: 25); for creatine, M+0 (Q1/Q3: 132/90, CE: 34), M+1 (Q1/Q3: 133/90, 133/91; CE: 34), M+2 (Q1/Q3: 134/90, 134/91, 134/92; CE: 34), M+3 (Q1/Q3: 135/90, 135/91, 135/92, 135/93; CE: 34); for creatinine, M+0 (Q1/Q3: 114/86, CE: 15), M+1 (Q1/Q3: 115/86, 115/87; CE: 15), M+2 (Q1/Q3: 116/86, 116/87, 116/88; CE: 15), M+3 (Q1/Q3: 117/86, 117/87, 117/88, 117/89; CE: 15); and for phosphocreatine, M+0 (Q1/Q3: 210/79, 210/97; CE: -26), M+1 (Q1/Q3: 211/79, 211/97; CE: -26), M+2 (Q1/Q3: 212/79, 212/97; CE: -26), M+3 (Q1/Q3: 213/79, 213/97; CE: -26).

Peaks for full scan data were integrated using El-Maven 0.12.0 software (Elucidata) using a targeted in-house spectral library detecting metabolites using precursor and product ion spectra^104^, and peaks for GAA tSIM data were integrated using a targeted method in TraceFinder 5.1 SP2 (Thermo Fisher OPTON-31001). Peaks from data obtained on the AB QTRAP 5500 and 6500+ were integrated using MultiQuant v2.1.1 (Sciex) software. For tracing experiments acquired on the AB QTRAP 5500, peak areas for each MRM corresponding to a given isotopologue species, as outlined above, were summed to yield the total abundance of each isotopologue. Total ion counts for full scan data were quantified using TraceFinder 5.1 SP2 software (Thermo Fisher OPTON-31001, RRID: SCR_023045). Peaks were normalized using probabilistic quotient normalization^105^ and total ion counts using R^48^ (RRID: SCR_001905) or by normalization to an internal standard. For stable isotope tracing studies, correction for natural abundance of metabolite labeling was performed using AccuCor 0.3.0^87^ (RRID: SCR_023046) in R (RRID: SCR_001905) or by manual correction.

##### Tissue Collection, Processing, and Extraction – Glioma Metabolomics Validation Cohort 1

De-identified tumor and matched normal brain tissue specimens were collected intraoperatively, immediately snap-frozen in liquid nitrogen, and stored at -80°C until shipment. Approximately 10 mg of frozen tissue per sample was submitted to Metabolon, Inc. (Morrisville, NC) for global metabolomic profiling. Recovery standards were added to each specimen prior to extraction to enable quality control monitoring of instrument performance and data normalization. Protein was precipitated by vigorous shaking with methanol, followed by centrifugation to recover chemically diverse metabolites. The resulting supernatant was split into multiple fractions optimized for four parallel UPLC-MS/MS methods: (1) reverse-phase positive ion electrospray ionization (ESI) optimized for hydrophilic species, (2) reverse-phase positive ion ESI optimized for hydrophobic species, (3) reverse-phase negative ion ESI, and (4) hydrophilic interaction chromatography (HILIC) negative ion ESI. A fifth aliquot was reserved as backup. Following drying under nitrogen, extracts were concentrated with a TurboVap® and maintained under inert gas overnight until analysis by MS.

Each fraction was reconstituted in a solvent system compatible with the intended UPLC-MS/MS method, each containing internal standards at fixed concentrations to verify injection and chromatographic consistency. Analyses were performed on a Waters ACQUITY UPLC system coupled to a Thermo Fisher Q-Exactive high-resolution mass spectrometer (Orbitrap, 35,000 FWHM resolution) equipped with a heated electrospray ionization (HESI-II) source. For reverse-phase methods, extracts were eluted from a Waters BEH C18 column using gradients of water, methanol, and acetonitrile containing 0.05% perfluoropentanoic acid and 0.1% formic acid. HILIC separations employed a BEH Amide column with water/acetonitrile and 10 mM ammonium formate at pH 10.8. MS analysis alternated between full MS and data-dependent MS^n^ scans across 70-1000 m/z, with dynamic exclusion applied to enhance coverage. Metabolites were identified by comparison with Metabolon’s spectral library, using retention time, accurate mass, and MS/MS data. Relative quantitation was performed by peak area integration, with values normalized to tissue mass. Missing data were imputed with the minimum observed value. Quality control included pooled technical replicates, blanks, and internal standards in every run. Instrument performance and reproducibility were monitored by calculating relative standard deviations, with experimental samples randomized and QC injections interspersed.

##### Absolute quantification of GAA in cells, conditioned media, and tissues

Metabolite preparation for absolute quantification of GAA in cells, media, and tissue was performed as described above. Standard curves for absolute quantification were generated using MultiQuant v2.1.1 or GraphPad Prism 10.4.1 by straight-line, non-linear regression with 1/x^2^ weighting. Parallel cultures for each cell line evaluated were used to obtain cell counts and average cellular diameter values using a Vi-CELL XR cell viability analyzer (Beckman Coulter). Total cellular volume was calculated using the formula 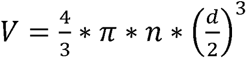, where *V* represents cell volume per µL acetonitrile extract, *d* = average cell diameter, and *n* = number of cells per µL extract. Media GAA content was calculated as 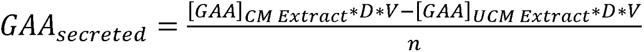, where [GAA] is the calculated concentration of GAA in conditioned media (CM) and unconditioned media (UCM) extract, *D* is the dilution factor of media in acetonitrile, *V* is the volume of media in which cells were incubated for the assay, and *n* represents the cell count in thousands. For quantification of media GAA secretion, a signal:noise threshold of conditioned > 2.5 × unconditioned media was applied to all samples, values below which were considered undetectable. For tissue samples, volumes were calculated using tissue weights and a previously published value of brain tissue density^106^.

##### LC-MS/MS assays for absolute quantification of GAA and creatine in tumor xenograft tissue

LC-MS/MS assays for the quantification of GAA and creatine in AGAT KO tumor studies were developed and validated according to the FDA guidelines for bioanalytical method validation^107^. Briefly, a LC-MS/MS system consisting of a Thermo Fisher Vanquish UPLC and Thermo Fisher TSQ Quantis Plus that was equipped with a heated ESI (HESI) source was used. The SRM transitions used for quantitation are as follows: for GAA, m+0 (Q1/Q3: 118.1/76.0), m+2 ISTD (Q1/Q3: 120.1/78.0); for creatine, m+0 (Q1/Q3: 132.1/90.0), m+5 ISTD (Q1/Q3: 137.1/95.2). 50 μL of plasma or brain tissue was protein precipitated using 450 μL of methanol containing the internal standards. The sample was vortexed and then centrifuged at 10,000 × g for 10 minutes before the supernatant was transferred and analyzed by LC-MS/MS. Chromatographic separation of the samples were accomplished with a Waters Acquity BEH c18 (2.1 x 100 mm, 1.7 μm) column with an isocratic elution using 80:20 (A:B, v:v) of water with 0.1% formic acid (A) and methanol (B) at a flow rate of 0.25 mL/minute. The total runtime was 3 minutes.

A standard curve was created for each run with duplicate QCs (low, medium, and high) to ensure the results for each day were accurate. The linear range of the standard curves for GAA and creatine in plasma was 50-25,000 ng/mL and 1-25,000 ng/mL in brain tissue. For both matrices, each analyte showed excellent linearity (r^2^ ≥ 0.9955). All FDA guideline criteria were met for each analyte in brain tissue. High analyte recovery was demonstrated (94% to 112%), and the intra- and inter-day accuracy (±11.2%) and precision (±8.7%) for quality controls were excellent. Other criteria, including dilution (±9.6%), stability after three 24 hour freeze thaw cycles (±4.7%), autosampler stability for 72 hours (±10.2%), bench top stability at RT for 4 hours (±5.3%), and matrix effect (±11.0%) for GAA and creatine also passed.

##### Immunoblot analysis of protein expression

Cells were lysed in EBC lysis buffer containing a protease inhibitor cocktail (Millipore Sigma 11836153001). Lysates were resolved by SDS-PAGE and transferred to nitrocellulose membranes (Bio-Rad 1620112 or Bio-Rad 1704158). Primary antibodies used included: anti-GAPDH (D16H11, Cell Signaling Technology 5174S, 1:5000, rabbit monoclonal, RRID: AB_10622025), anti-vinculin (Sigma V9131, 1:100,000, mouse monoclonal, RRID:AB_477629), anti-AGAT (Abcam 119269, 1:1000, mouse monoclonal, RRID: AB_10902241), anti-GAMT (PTG 10880-1-AP, 1:1000, rabbit polyclonal, RRID: AB_2109304). HRP-conjugated secondary antibodies used included: anti-Mouse IgG (Thermo Fisher 31430, 1:2,000, goat polyclonal, RRID: AB_228307) and anti-Rabbit IgG (Thermo Fisher 31460, 1:2,000, goat polyclonal, RRID: AB_228341). Densitometry analysis was performed in ImageJ.

##### ATAC-sequencing and Transcription Factor Footprinting analysis of *GATM*

Omni-ATAC-seq was performed as described by Corces et al^108^. 50,000 cells were collected and DNase treated for 30 minutes at 37°C prior to nuclei extraction and transposition. After transposition, cleanup was performed using Zymo Clean and Concentrator-5 Kit (Zymo Research D4013) and DNA was eluted in 21 uL of elution buffer. To add adaptor sequences, 5 cycles of amplification were performed using NEBNext 2× Master Mix and qPCR (New Engalnd Biolabs M0541S) was used to determine the addition of 0-3 cycles based on sample amplification profiles. Samples were purified using Zymo Clean and Concentrator-5 Kit. Libraries were sequenced on the NextSeq 500 (Illumina, RRID:SCR_014983) using paired-end sequencing of 75 base pair reads by the Molecular Biology Core Facilities at Dana Farber Cancer Institute.

Fastq files associated with NHA, AGAT-low, and AGAT-high GSCs were analyzed using the Nextflow-based BICF Astrocyte 2.2.0 ATAC-seq analysis workflow from the UT Southwestern Bioinformatics Core. Briefly, adaptors were trimmed with TrimGalore^89^, and reads mapped with BWA^90^, filtered with SamTools^92^, and sorted with Sambamba^91^. Duplicates were then marked with Sabamba, reads filtered with SamTools, and percentage of reads in mitochondria calculated and library complexity calculated with SamTools and bedtools^93^. Cross-correlation was calculated using PhantomPeakQualTools^94, 110^, peaks were called with MACS2^95^ from overlaps of pooled replicates, and consensus peaks were called and annotated from ChipSeeker output files.

Footprinting analysis was performed using TOBIAS through UT Southwestern BioHPC. Peaks were filtered for only Chr peaks and to remove blacklisted peaks. .narrowPeak output from the above ATACseq workflow was then merged from all samples and replicate bam files were merged with SamTools. Bam files were then merged according to AGAT expression and malignancy status.

##### Splice isoform analysis of *GATM*

Isoform-specific *GATM* gene expression data were generated by reanalysis of RNAseq fastq files using the hg38 build through Astrocyte 2.2.0 to derive relative expression of transcript variants. Differential gene expression analysis was then performed through Astrocyte 2.2.0.

##### Single cell RNA sequencing analysis of human GBM samples

Human GBM single cell RNA sequencing analysis was conducted using a previously published dataset (GSE274546)^78^. Single cell UMI count files from all primary tumors were downloaded from the NCBI Sequence Read Archive (SRA) and processed using the R package Seurat. Cells with less than 1000 or more than 7000 genes, or with greater than 3% mitochondrial fraction were excluded from further analysis. Subsequent steps including normalization, finding highly variable genes, scaling, cell clustering and UMAP construction were performed. Marker genes for each Seurat cluster were identified by using the Seurat “FindAllMarkers” function, cell clusters were annotated by marker gene expression as in the original publication^78^. Gene expression was visualized by using the Seurat “VlnPlot” function.

##### RNA sequencing and transcriptional subtype classification of human primary glioma samples

Total RNA from primary human glioma samples was extracted using RNeasy Plus Universal kit (Qiagen 73404) according to the manufacturer’s instructions. RNA libraries were generated using Kapa RNA HyperPrep Kits with RiboErase (HMR) (Roche KK8561) following manufacturer instructions. Briefly, all RNA samples underwent rRNA depletion, RNAse H and DNase treatment and fragmentation. The RNA fragments were used as template for cDNA synthesis. The cDNA fragments went through the ligation process with unique molecular identifier adapters synthesized by IDT. The products were purified and enriched with PCR amplification to create the final cDNA library. The library quality was verified on Agilent 2100 Bioanalyzer and sequenced on Illumina NovaSeq 6000 sequencer platform using 150 bp paired-end protocol at the UT Southwestern Genomics & Microarray core.

Sequencing adapters from raw reads were trimmed using Trim Galore (v0.6.4). Trimmed reads were aligned to the human genome (hg38) using STAR (v2.7.3a). Mapped reads were quantified using featureCounts from the Subread package (v1.6.3). Transcriptional subtype classification was done by scoring each glioma samples with gene signatures^25, 111^ of each GBM subtype by the ssGSEA method from the R package GSVA.

##### Immunohistochemistry of human brain samples

H&E and immunohistochemistry procedures for the spatially-resolved human glioma case (patient ID #5852) were performed at HistoWiz, Inc, using the Leica Bond RX automated stainer (Leica Microsystems), a Standard Operating Procedure, and a fully automated workflow. Samples were fixed in 10% formalin for 24 hours, washed in 70% ethanol, and embedded in paraffin. Samples were shipped as paraffin embedded blocks and sectioned at 4 μm. One section per sample was stained in hematoxylin and eosin (H&E). Immunohistochemistry (IHC) was conducted using a Leica Bond RX automated stainer (Leica Microsystems). HistoWiz in-house murine skeletal muscle and kidney tissue were used as negative and positive controls, respectively, for optimization of IHC staining for both AGAT and GAMT. Tissue slides were dewaxed with xylene and alcohol-based solutions. Heat-induced epitope retrieval (HIER) was performed for 20 minutes using BOND Epitope Retrieval Solution 1 (H1-20; citrate-based buffer, pH 6.0; Leica Biosystems AR9961) for the GATM antibody, and Epitope Retrieval Solution 2 (H2-20; EDTA-based buffer, pH 9.0 AR9640) for the GAMT antibody. Slides were then incubated with anti-GATM (HPA026077, Atlas Antibodies, RRID: AB_1849528) at a 1:100 dilution and anti-GAMT (PA5-119778, Thermo Fisher, RRID:AB_2913350) at a 1:5,000 dilution for 30 minutes at room temperature. Signal detection was carried out using the Leica Bond Polymer Refine Detection Kit (Leica Biosystems DS9800, RRID:AB_2891238), with 3,3’- diaminobenzidine (DAB) as the chromogen and counterstained with hematoxylin following the manufacturer’s protocol. After staining, slides were dehydrated and coverslipped using the Tissue-Tek Film® Automated Coverslipper (Sakura Finetek). Whole-slide images were acquired at 40× magnification using a Leica Aperio AT2 scanner (Leica Microsystems, RRID:SCR_021256).

##### Histology and immunofluorescence of murine PDX samples

Mouse brain tissue for histology and immunofluorescence was dissected and fixed in 10% neutral buffered formalin (VWR 95042-908) and further processed for embedding and sectioning by the Histology Core at the University of Pittsburgh. H&E was performed by the Tissue Management and Shared Resource at the UT Southwestern Medical Center (UTSW), and the slides were imaged using a Hamamatsu Nanozoomer S60 (RRID:SCR_023762) at the UT Southwestern Whole Brain Microscopy Facility.

For immunofluorescence labeling, paraffin-embedded brain was sectioned at 4 µm and deparaffinized in xylene and re-hydrated using graded ethanol series and distilled water. Antigen retrieval was performed by boiling slides in 1 mM EDTA (pH 8.0) or citric acid buffer (pH 6.0) in a microwave oven for 25 minutes, followed by blocking (10% normal donkey serum, 0.4% Triton X-100 in PBS) at room temperature for 1 hour. Slides were incubated at 4 °C overnight in the following primary antibodies: anti-KCC2 (1:500, Millipore Sigma C2366), anti-cFOS (1:1000, Synaptic System 226009, RRID: AB_2943525); anti-cFOS (1:250, Cell Signaling Technology 2250, RRID:AB_2247211) ; anti-KU80 (1:250, Cell Signaling Technology 2180, RRID: AB_2218736), and anti-vGLUT1 (1:500, Millipore Sigma AB5905, RRID: AB_2301751). The following day, slides were incubated at room temperature for 1 hour with secondary antibodies: 1:500 Alexa Fluor 488 AffiniPure donkey anti-rabbit IgG (H+L) (Jackson ImmunoResearch, 711-546-152, RRID:AB_2340619); 1:1000 Alexa Fluor 488 donkey anti-chicken IgΥ (H+L) (Invitrogen, A78948); 1:1000 Alexa Fluor 594 goat anti-rabbit IgG (H+L) (Invitrogen, A11012, RRID:AB_2534079); 1:500 Alexa Fluor 647 AffiniPure donkey anti-guinea pig IgG (H+L) (Jackson ImmunoResearch, 706-606-148, RRID:AB_2340477), and 1:1000 Alexa Fluor 647 donkey anti-rabbit IgG (H+L) (Invitrogen A32795).

Both primary and secondary antibodies were diluted in PBS containing 10% normal donkey serum and 0.2% Triton X-100. The nuclei and slides were stained and mounted by prolong gold antifade reagent with DAPI (Cayman Chemical 14285) followed by imaging using the Zeiss LSM880 with Airyscan (RRID:SCR_020925) Confocal Scanning Microscope in the Quantitative Light Microscopy Core (QLMC) at the UTSW or Zeiss Axioscan 7 Digital Slide Scanner (RRID:SCR_027284) at the Whole Brain Microscopy Facility at the UTSW. Quantification of immunofluorescent images was performed using ImageJ 1.43p (Wayne Rasband and contributors, National Institutes of Health, USA).

##### Acute brain slice preparation

Acute hippocampal slices used in single-cell electrophysiological recordings in experiments in which slices were treated with GAA or creatine in the presence or absence of gabazine were prepared as previously described^109^ from C57BL/6J mice aged 5-7 weeks old. Briefly, mice were decapitated and the brain was rapidly extracted in oxygenated (95% O2 and 5% CO2) and ice-cold sucrose-based artificial cerebrospinal fluid (sucrose-aCSF) at pH 7.4 and 300 mOsm containing: 185 mM sucrose (Sigma Aldrich S9378), 25 mM NaHCO_3_ (Thermo Fisher, BP328), 2.5 mM KCl (Millipore Sigma, P9333), 25 mM glucose (Millipore Sigma, G8270), 1.25 mM NaH_2_PO_4_ (Sigma Aldrich S8282), 10 mM MgCl_2_ (Millipore Sigma, M9272), and 0.5 mM CaCl_2_ (Millipore Sigma, 223506). The brain was dissected, glued to the specimen holder, and acute slices (300 µm) were prepared on a vibratome (Leica VT1200 S). Slices were transferred to oxygenated and heated (32°C) sucrose-aCSF for 30 minutes. After 30 minutes, the water bath temperature was turned off and slices were transferred to oxygenated recording aCSF at pH 7.4 and 300 mOsm containing: 125 mM NaCl (Millipore Sigma S9888), 25 mM NaHCO_3_ (Thermo Fisher BP328), 2.5 mM KCl (Millipore Sigma P9333), 10 mM glucose (Millipore Sigma G8270), 2 mM CaCl_2_ (Millipore Sigma 223506), and 2 mM MgCl_2_ (Millipore Sigma M9272), and allowed to recover for an additional 30 minutes before recordings. Slices were maintained in oxygenated aCSF at room temperature for the rest of the day.

Brains for experiments measuring changes in input resistance in response to GAA in the presence or absence of inhibitors of glutamatergic and GABAergic signaling were prepared as described previously^112^. Briefly, wild-type male mice on postnatal days 18-23, were anesthetized by intraperitoneal injection of ketamine/xylazine. After toe pinch to confirm sufficient depth of anesthesia, mice were euthanized by rapid decapitation. Acute coronal slices (300 µm thickness) containing somatosensory barrel cortex were generated using a Leica VT1200S Vibratome in a semi-frozen 300 mOsm dissection buffer containing: 110 mM choline chloride (Millipore Sigma, C1879), 25 mM NaHCO_3_ (Millipore Sigma, S6014) 25 mM D-glucose (Millipore Sigma, G8270), 11.6 mM ascorbic acid (Millipore Sigma, A92902), 2.5 mM KCl (Millipore Sigma, P5405), 1.25 mM Na_2_H_2_PO_4_ (Millipore Sigma, S9638), 3.1 mM Na-pyruvate (Millipore Sigma, P5280), 7 mM MgCl_2_ (Millipore Sigma, M8266), 0.5 mM CaCl_2_ (Millipore Sigma, C7902), and 1 mM kynurenic acid (Millipore Sigma, K3375). Brains and slices were continually aerated with 95% O_2_ and CO_2_ prior to and during the slicing procedure. After slicing, slices were then transferred to 300mOsm normal artificial cerebrospinal fluid (ACSF) solution containing: 125 mM NaCl (Millipore Sigma, S7653), 25 mM NaHCO_3_ (Millipore Sigma, S6014), 10 mM D-glucose (Millipore Sigma, G8270), 2.5 mM KCl (Millipore Sigma, P5405), 1.25 mM Na_2_H_2_PO_4_ (Millipore Sigma, S9638), 2 mM MgCl_2_ (Millipore Sigma, M8266), and 2 mM CaCl_2_ (Millipore Sigma, C7902) to recover at 37°C for 30 minutes, then at 21°C for 30 minutes prior to recording.

Acute brain slices for experiments assessing neuronal network activity in tumor-bearing and contralateral brain hemispheres were obtained from 14-week-old adult mice, 3 weeks following stereotactic xenograft injection. Slices were prepared at a thickness of 300 µm using a VT1000P vibratome (Leica) in oxygenated, ice-cold cutting buffer containing 205 mM sucrose (Thermo Fisher S5), 2.5 mM KCl (Thermo Fisher P330), 1.25 mM NaH PO (Thermo Fisher S397), 26 mM NaHCO (Thermo Fisher P233), 10 mM glucose (Gibco 15023-021), 0.5 mM CaCl (Thermo Fisher 012316.A1), and 5 mM MgSO (Thermo Fisher 033337.36). Following dissection, slices were incubated in recording buffer at 34°C for 40 minutes.

##### Single cell patch-clamp electrophysiological recordings

Slices generated for electrophysiology experiments in which slices were treated with GAA or creatine in the presence or absence of gabazine were transferred under an upright microscope (Scientifica) and held under a harp in a chamber. Slices were perfused at 2 mL/minute with oxygenated aCSF heated to 32°C. The CA1 pyramidal cell layer was visually identified under a 4× objective (NA = 0.1, Olympus) and visually targeted for whole-cell patch clamp recordings using a 40× objective (NA = 0.8, Olympus). Electrodes were made from borosilicate glass capillaries (BF150-86-10HP, Sutter Instrument) on a P-1000 micropipette puller (Sutter Instrument). Recording electrodes were filled using a pH 7.3 and 290 mOsm solution containing: 130 mM Cs-methanesulfonate (Millipore Sigma C1426), 10 mM HEPES (Millipore Sigma H3375), 2 mM MgCl_2_ (Millipore Sigma M9272), 4 mM ATP (Millipore Sigma A9187), 0.3 mM GTP (Millipore Sigma G8877), 7 mM Phosphocreatine di(tris) (Millipore Sigma P7936), 0.6 mM EGTA (Millipore Sigma E3889), and 5 mM KCl (Millipore Sigma P9333). The electrophysiological signal was amplified using a Multiclamp 700B (Molecular Devices) and digitized at 10 kHz with a Digidata 1550B (Molecular Devices) before being displayed and recorded on a personal computer using Clampex v11.3 (Molecular Devices). After establishing the whole cell configuration, neurons were assessed for electrophysiological parameters and voltage-clamped at 0 mV to begin experiments. Only neurons with a resting membrane potential more hyperpolarized than -55 mV and access resistance < 30 MΩ were included for analysis. The liquid junction potential was not corrected. Recording epochs consisted of a 3 minute baseline with aCSF, followed by 1 minute application of GAA (100 µM) or creatine (100 µM), and 6 minutes aCSF. GAA and creatine were dissolved in aCSF and prepared fresh every day. 10 µM gabazine (EMD Millipore SR95531) was pre-applied and continuously applied in experiments where indicated. aCSF and all compounds were delivered using a gravity-driven perfusion system. Data from hippocampal patch-clamp slice recordings was analyzed in Clampfit v11.4 and Igor Pro v9.05. The area under curve was measured in Clampfit between recording time 4-10 minutes.

Recordings for experiments measuring changes in input resistance in response to GAA in the presence or absence of inhibitors of glutamatergic and gabaergic signaling were conducted at 21°C in a submerged patch-clamp setting and perfused with oxygenated (95% O_2_/5% CO_2_) ACSF (1mL/minute) as described previously^112^. Depending on experiment, the ACSF contained the following inhibitors of neurotransmitter-gated ion channels: 10 µM CPP (Hello Bio HB0036), 20 µM DNQX (Tocris 2312), 100 µM PTX (Hello Bio HB0506), 2 µM CGP 52432 (Tocris 1246), and/or 1 µM tetrodotoxin (Hello Bio HB1034). Layer V pyramidal neurons were visualized with differential interference contrast microscopy. Patch pipettes with 5-7 MΩ were pulled from borosilicate glass. Whole-cell recordings were performed on layer 5a pyramidal neurons using a Multiclamp 700A amplifier (Molecular Devices). Internal solution contained: 130 mM K-gluconate (Millipore Sigma P1847), 0.2 mM EGTA (Millipore Sigma E3889), 6 mM KCl (Millipore Sigma P5405), 3 mM NaCl (Millipore Sigma S7653), 10 mM HEPES (Millipore Sigma H4034), 14 mM phosphocreatine-tris (Millipore Sigma P1937), 4 mM Mg-ATP (Millipore Sigma A9187) and 0.4 mM Na-GTP (Millipore Sigma G3776). Recordings were conducted in voltage clamp, holding the membrane potential at -65 mV. Neurons with resting membrane potential >-60mV and series resistance <40 MΩ were included in analysis. Input resistance in voltage clamp was measured with a -10 mV voltage step. For each recording, baseline electrophysiological measurements were obtained, followed by a 100 µM GAA wash-in for 5 minutes. Following 5-minute GAA wash-in, post wash-in measurements were made. Data acquisition and analysis was performed using custom Labview 8.6 software (National Instruments). Input resistance following GAA treatment was normalized to pre-treatment input resistance averages for each neuron.

##### Multielectrode Array Recording

Spike counts were recorded from acute slices using a 4096-electrode Complementary Metal– Oxide Semiconductor (CMOS)-based HD-MEA (BioCam DupleX with CorePlate™ 3D, 3Brain). Prior to use, chips were rinsed three times with deionized water, treated with 70% ethanol, re-rinsed, and preconditioned with PBS.

After incubation at 34°C, slices were incubated an additional 40-minute incubation at room temperature prior to recording. CLP257-treated slices were incubated in 30 μM CLP257 (Bio-Techne 5242/10) for 30 minutes immediately before recording. The recording buffer consisted of 126 mM NaCl (Thermo Fisher P271), 3.5 mM KCl (Thermo Fisher P330), 1.25 mM NaH_2_PO_4_ (Thermo Fisher S397), 26 mM NaHCO_3_ (Thermo Fisher P233), 10 mM glucose (Gibco 15023-021), 1.6 mM CaCl_2_ (Thermo Fisher 012316.A1), and 1.25 mM MgSO_4_ (Thermo Fisher 0333337.36). Spontaneous activity was sampled continuously for 5 minutes at 20 Hz across sequential pharmacological conditions: baseline aCSF and 20 µM GAA. Recordings were acquired under the following conditions (*n* = 3 slices per condition): Non-tumor-bearing left hemisphere, tumor-bearing right hemisphere, non-tumor-bearing left hemisphere plus 30 µM CLP-257 (Bio-Techne 5242/10), and tumor-bearing right hemisphere plus 30 µM CLP-257.

Spike detection was performed using the Precise Timing Spike Detection (PTSD)^113^ algorithm in BrainWave 5 software (3Brain), with a threshold set at 8 standard deviations. Spike counts were analyzed in R (v4.5.0); data were reshaped into long format using tidyverse^100^ tools and log-transformed as log (spike count + 1).

##### Cell growth assay

TS516 AGAT WT and AGAT KO cells were plated in 6-well ultra-low adherence plates at 2 × 10^5^ cells per well in NeuroCult NS-A Basal Medium (Human) prepared as described above. Cell counts were obtained at 0, 24, 48, 72, 96, and 120 hours following dissociation in Accutase (StemCell 07922) using a Vi-CELL XR cell viability analyzer. Media on remaining wells was changed at 48 and 96 hours. Growth rates were compared using nonlinear regression to fit exponential growth curves in GraphPad Prism 10.4.1.

##### Xenograft studies of AGAT expression

Mice were randomly assigned to receive intracranial injections of either TS516 AdCRISPR AGAT WT or TS516 AdCRISPR AGAT KO cells. Cells were injected in 5 µL NeuroCult NS-A Basal Medium (Human) 0.5-1 mm anterior and 2 mm lateral to the bregma, at a depth of 3 mm from the brain surface. Animals were monitored daily until the appearance of neurological symptoms, at which point they were euthanized.

##### Xenograft studies of dietary GAA modulation

Mice were randomly assigned to receive one of the following diets: Amino Acid Diet (Inotiv TD.01084) or Arginine-Deficient, 0.665% Ornithine Diet (GAMT Deficiency Diet, Inotiv TD.240691). Cells were injected in 5 µL NeuroCult NS-A Basal Medium (Human) 0.5-1 mm anterior and 2 mm lateral to the bregma, at a depth of 3 mm from the brain surface. Mice were monitored daily for the development of neurological symptoms, at which point they were euthanized.

##### Secretion of GAA by human tissue explants

Tumor tissue was collected and tissue explants generated as described previously^48^. Briefly, tissue collected from the operating room was suspended in Hibernate A (BrainBits HA) and transferred to RBC Lysis Buffer (Thermo Fisher 00433357) within 30 minutes of excision for 10 minutes with rocking. Tissue was washed three times with Hibernate A supplemented with 2 mM GlutaMAX (Thermo Fisher 35050061), 100 U/mL and 100 μg/mL (respectively) penicillin/streptomycin (Thermo Fisher 15140148), and 250 ng/mL amphotericin B (GeminiBio 400104). Dissection scissors or scalpels were used to parcellate tissues into 1-2 mm^3^ pieces which were then plated in separate wells of a 24-well ultra-low adherence plate (Corning 3473) in 1 mL HPLM supplemented with 1× B27 supplement minus vitamin A (Thermo Fisher 12587010), 1× N2 supplement (Thermo Fisher 17502048), 100 U/mL and 100 μg/mL (respectively) penicillin/streptomycin (Thermo Fisher 15140148), 250 ng/mL plasmocin (InvivoGen ant-mpp), 55µM 2-mercaptoethanol (Thermo Fisher BP176100), and 2.375-2.875 µg/mL insulin (Millipore Sigma I9278). Tissues were randomized to groups of 1 to 4 technical replicates per tracer from each biologic sample, depending on available tissue; remaining tissue was used for histopathology. After 30 minutes, media was exchanged for 1 mL HPLM as described previously^48^. Tissues were incubated for 18 hours in a 37°C incubator at ambient oxygen and 5% CO2 with shaking. After 18 hours, conditioned media was harvested, snap-frozen in liquid nitrogen, and stored at -80°C until analysis. Tissues were then prepared for LC-MS analysis and quantification of GAA secretion in media as described above. As in quantification of media GAA secretion by GSCs, a signal:noise threshold of GAA concentration in conditioned > 2.5 × unconditioned media was applied to all samples to discriminate GAA-secreting samples. As no cell count information could be obtained for these explants, secreted GAA was rendered as the unconditioned-subtracted GAA content of conditioned media for those samples whose GAA content exceeded this threshold. Outliers within technical replicates were identified and excluded by ROUT test (Q=1%).

##### Histopathology of human explants

Residual human tumor and brain tissues from explant preparation were fixed for 1 hour in neutral buffered 10% formalin solution (Millipore Sigma HT501128). Following fixation, tissues were washed and stored in 70% ethanol. Tissues were embedded in paraffin, sectioned, and stained with hematoxylin and eosin (H&E) by the University of Texas Southwestern Histo Pathology Core. H&E sections were reviewed by a board-certified neuropathologist (T.E.R).

### QUANTIFICATION AND STATISTICAL ANALYSIS

#### Analysis of nitrogen metabolism profiling platform data

Analysis of nitrogen metabolism profiling platform data was performed as previously described^48^. Briefly, integrated peaks from LC-MS analysis of nitrogen metabolism profiling platform data were analyzed using R (RRID: SCR_001905). Metabolites which were not quantified in more than one sample, had a mean total pool size below a threshold of 1 × 10^6^, had calculated total labeling across all conditions of less than 1%, had quantified labeling of >4% in any unlabeled sample, had calculated total labeling across all conditions of greater than 500% were filtered and removed from subsequent analysis. Differential labeling scores were calculated using the formula: 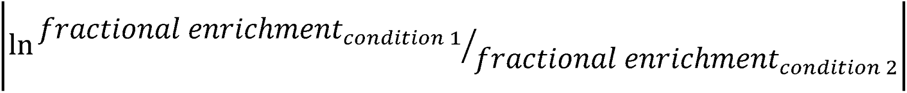 × 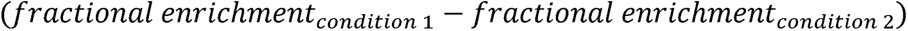For visualization, metabolites with less than 5% total labeling were filtered and removed from Sankey diagrams.

#### TCGA and GTEX RNA sequencing analysis

Results are based in part upon data generated by the TCGA Research Network: https://www.cancer.gov/tcga^74^. RNA sequencing data for patient glioblastoma samples were downloaded from the TCGA portal (https://portal.gdc.cancer.gov/projects/TCGA-GBM), comprising 173 samples including 154 index tumor samples, 14 recurrent tumor samples, and 5 normal controls sequenced by Illumina HiSeq 2000 RNA Sequencing. Of these, 151 primary GBM tissue samples with accompanying survival and prognostic information were selected for analysis. RNA sequencing data from non-malignant brain tissue samples were downloaded from the GTEx portal (https://gtexportal.org, file version: 2017-06-05_v8_brain_cortex, accession date August 26, 2019)^75, 76^. Gene expression values for *GATM, GAMT, CKB, CKMT1A, CKMT2A,* and *CKMT2B* were first converted to Transcripts Per Million (TPM) to enable comparison across samples. A log (TPM + 1) transformation was then applied to stabilize variance and normalize the distribution. For each gene, Z-score transformation was performed across all samples to standardize expression levels and highlight relative up- or down-regulation. Samples were annotated according to associated histological subtype, IDH mutation status, CIMP status, and MGMT promoter methylation.

#### Other statistical analyses

Information related to data presentation and statistical analysis for individual experiments can be found in the corresponding figure legends. Statistical analyses were carried out using GraphPad Prism software (version 10.4.1). Significance of all comparisons involving two groups was calculated by unpaired two-tailed *t*-test. For comparisons of two groups with significantly different variances, Welch’s *t*-test was used. For comparisons of two groups without significant differences in variances, Student’s *t*-test was used. Mann-Whitney tests or Kolmogorov-Smirnoff tests were used for comparisons between groups in which assumptions of normality were not appropriate. Significance of all comparisons involving three or more groups was calculated by one-way ANOVA. Survival data from mouse xenograft studies was analyzed and statistical significance was determined using log-rank tests. For all tests, two-tailed *p-*values < 0.05 were considered statistically significant.

## SUPPLEMENTAL INFORMATION

### SUPPLEMENTAL FIGURE LEGENDS

**Figure S1. Composition of metabolomics dataset, related to Figure 1**. (**A**) Pie chart depicting composition of human tissue sample cohort by tissue type. LGG = lower grade glioma. HGG = high grade glioma. (**B**) Heatmap depicting gene expression analyses and transcriptional subtype clustering of human HGG (*n*=39) and LGG (*n*=25) tissue samples. (**C-D**) Partial least-squares discriminant analysis (PLS-DA) of IDH mutant tumors by (C) tumor histology and (D) tumor grade. (**E**) PLS-DA of GBM tissue samples categorized by transcriptional subtype. (**F**) Peak intensities of 2-hydroxyglutarate (2HG) in IDH-mutant (IDHmut) and IDH-wild-type (IDHwt) glioma samples and non-malignant brain samples. (**G-K**) Peak intensities of (G) L-DOPA, (H) aminoadipate, (I) orotate, (J) deoxyguanosine, and (K) ratio of peak intensities of N-acetylaspartate to creatine in HGG, LGG, and non-malignant brain samples. Data are presented as means. n.s.= not significant, *p<0.05, **p<0.01, ***p<0.001 (Welch’s *t*-test).

**Figure S2. Comparison of creatine metabolism pathway intermediates by tumor subtype, related to Figure 2**. (**A-D**) Peak intensities for (A) GAA, (B) creatine, (C) phosphocreatine, and (D) creatinine in tissue samples from index (*n*=15) and recurrent (*n*=10) LGG tumors. Data are means; *p<0.05 (Unpaired t-test for creatine; Welch’s t-test for all others). (**E-H**) Peak intensities for (E) GAA, (F) creatine, (G) phosphocreatine, and (H) creatinine in tissue samples from both index and recurrent HGG tumors and non-malignant brain specimens, stratified by diagnostic and transcriptomic subtypes (Astro, IDHmut, Gr4; *n*=4) (GBM; *n*=35) (non-malignant brain; *n*=9). GBMs were classified as classical (*n*=17), mesenchymal (*n*=12), or proneural (*n*=6) transcriptomic subtypes. (**I-L**) Peak intensities for (I) GAA, (J) creatine, (K) phosphocreatine, and (L) creatinine in tissue samples from squamous cell lung carcinoma (*n*=35) or tumor-adjacent non-malignant lung (*n*=35). Peak intensities are reanalyzed from Moreno et al. (**M-P**) ROC analyses of GAA content as a discriminant marker of (M) HGG (*n*=39) compared to non-malignant brain (*n*=9), (N) HGG (*n*=39) compared to LGG (*n*=25), or LGG (*n*=25) compared to non-malignant brain (*n*=9) including (O) both index and recurrent tumor resection samples, and (P) index LGGs only (*n*=15) compared to non-malignant brain (*n*=9). Data are presented as means. n.s.=not significant, *p<0.05, ***p<0.001 (Welch’s t-test in A, C-D, I, and K-L; unpaired *t*-test in B and J; one-way ANOVA in E-H).

**Figure S3. Arginine tracer accumulation in non-malignant and glioma cell lines, related to Figure 3**. Fractional enrichment of label from ^15^N_4_-arginine in intracellular arginine pools following 18 hour incubation of non-malignant cells (*n*=4), IDH-mutant GSCs (*n*=6), and IDH-wild-type GSCs (*n*=7) in tracer-containing HPLM (*n*=3 per cell line). Data are means ± SEM.

**Figure S4. Transcriptional regulation of creatine synthesis pathway enzymes, related to Figure 4**. (**A-C**) Expression of (A) AGAT, (B) GAMT, and (C) ratio of AGAT:GAMT expression by cell type in human GBM by single-cell RNA sequencing. Data are reanalyzed from Nomura et al^78^. (**D**) GAA secretion by HMC3 microglia, NHA Donor #1, and TS516 cells (*n*=3 per line). n.s.= not significant, ***p<0.0001. (**E**) Diagram of mRNA splice isoforms expressed from the *GATM* gene. (**F**) Heatmap of RNA sequencing of non-malignant (NHA Donor #1), AGAT^low^ (MGG152, BT260), and AGAT^high^ (all other) GSC cell lines comparing relative abundance of *GATM* mRNA splice isoforms (*n*=2 per line). Data are fragments per kilobase transcript per million mapped reads (FPKM). n.s.= not significant, ***p<0.001 (unpaired *t*-test).

**Figure S5. Arginine tracer accumulation in NHA stable lines, related to Figure 5**. **(A)** Fractional enrichment of label from ^15^N_4_-arginine in intracellular arginine pools following 18 hour incubation in tracer-containing HPLM of NHA Donor #1 stable lines expressing either empty vector (EV), wild-type AGAT cDNA (AGAT WT), or catalytically-dead AGAT cDNA (AGAT C407A) and Cas9 with either an sgRNA targeting the *AAVS1* safe-harbor locus control or one of two sgRNAs targeting *GAMT.* **(B)** Fractional enrichment of label from ^15^N_4_-arginine in intracellular arginine pools following 18 hour incubation in tracer-containing HPLM of NHA Donor #1 stable lines expressing either empty vector (EV), wild-type AGAT cDNA (AGAT WT), or catalytically-dead AGAT cDNA (AGAT C407A) and EV, wild-type GAMT cDNA (GAMT WT), or catalytically-dead GAMT cDNA (GAMT E45S) (*n*=3 per line). Data are means ± SEM.

**Figure S6. Investigation of creatine synthesis pathway enzyme expression and GAA-induced hyperexcitability in the HGG microenvironment, related to Figure 6**. (**A-B**) Immunohistochemistry staining of kidney (positive control) and skeletal muscle (negative control) tissues for (A) AGAT and (B) GAMT. Scale bars=100 µm. (**C**) Neuronal activity in tumor-infiltrated brain slices before (0-5 minutes) and after (5-10 minutes) treatment with 20 µM GAA as measured by MEA assay. Representative neuronal spiking is shown in aggregate (top panels) and across electrodes (bottom panels).

**Figure S7. Investigation of GAA-induced hyperexcitability in vivo, related to Figure 7**. (**A-B**) Standard curves with quality control samples (*n=*2 per concentration) for absolute quantification of (A) GAA and (B) creatine in a custom LC-MS assay. (**C**) Representative immunofluorescence analysis of cFOS and NeuN markers in tumor-adjacent neocortex regions of AGAT WT and AGAT KO TS516 xenografted mouse brains. (**D**) Quantitation of peritumoral cortical cFOS-positive cells co-expressing NeuN in AGAT WT TS516 xenografted mouse brains (*n*= 3 mice). (**E-F)** Immunofluorescence analysis of vGLUT1 and Ku80 markers in (E) AGAT WT and (F) AGAT KO TS516 xenografts. ROIs denote regions highlighted in Figures 7K and 7L. Scale bars = 1 mm (wide view) and 100 µm (close view insets). (**G**) Relative body weights and (**H**) diet consumption for mice receiving standard diet or GAMT deficiency diet (*n*=3 mice per arm). Data are means ±SEM.

