## Supplementary figures and images for "Gliomas phenocopy an inborn error of metabolism to drive neuronal activity and tumor growth"

### Supplemental Figure 1

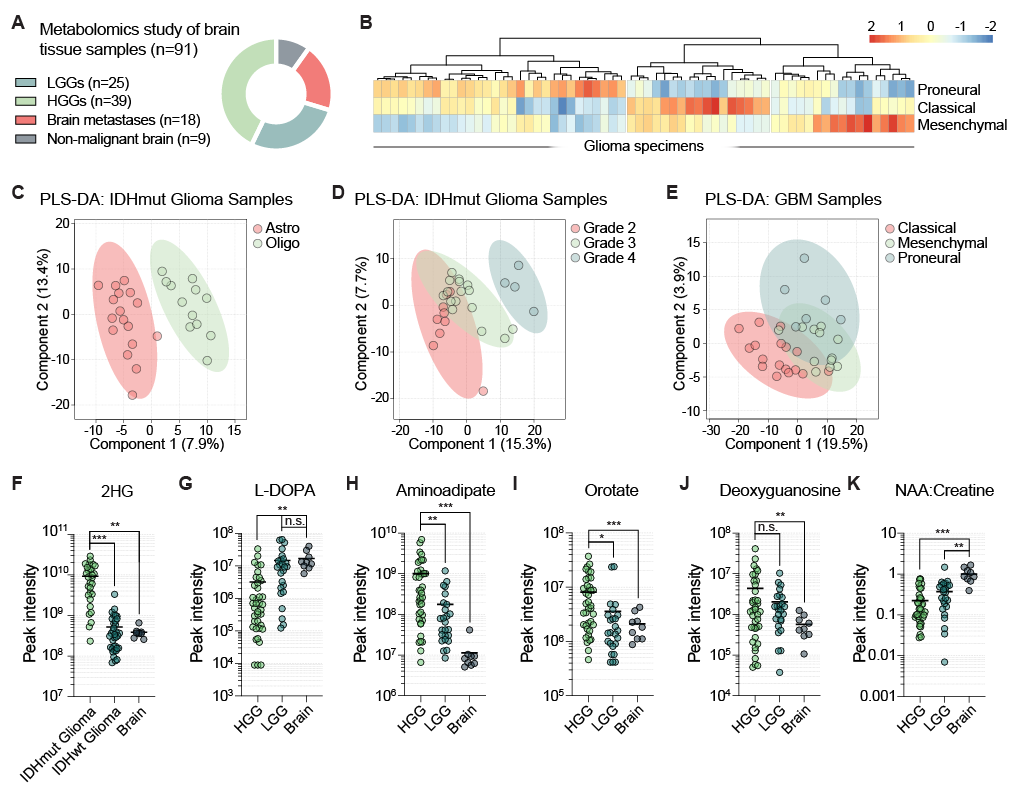

### Supplemental Figure 2

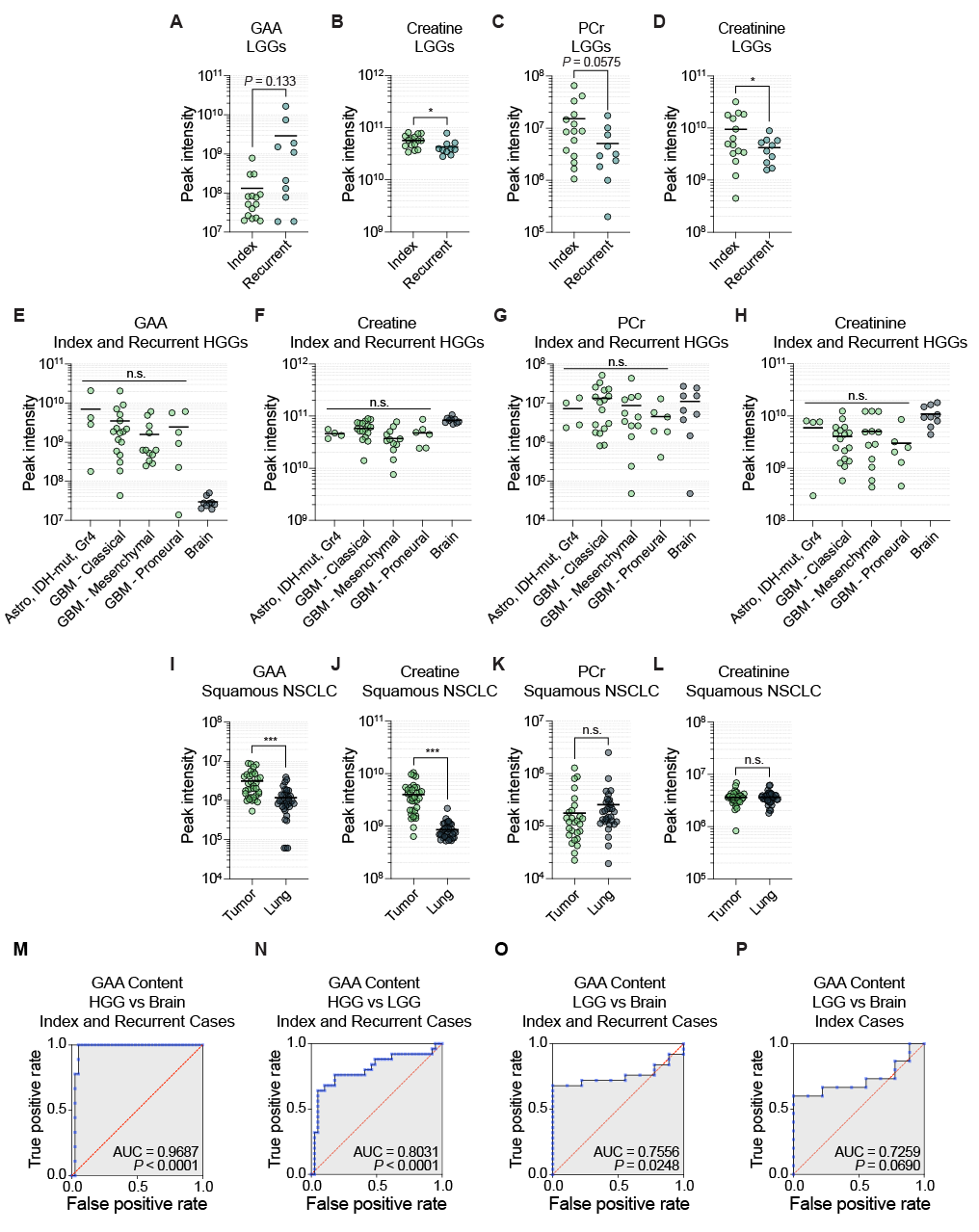

### Supplemental Figure 3

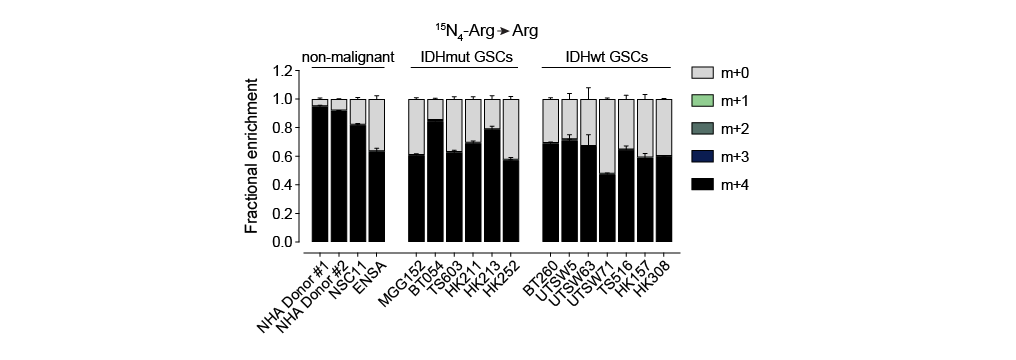

### Supplemental Figure 4

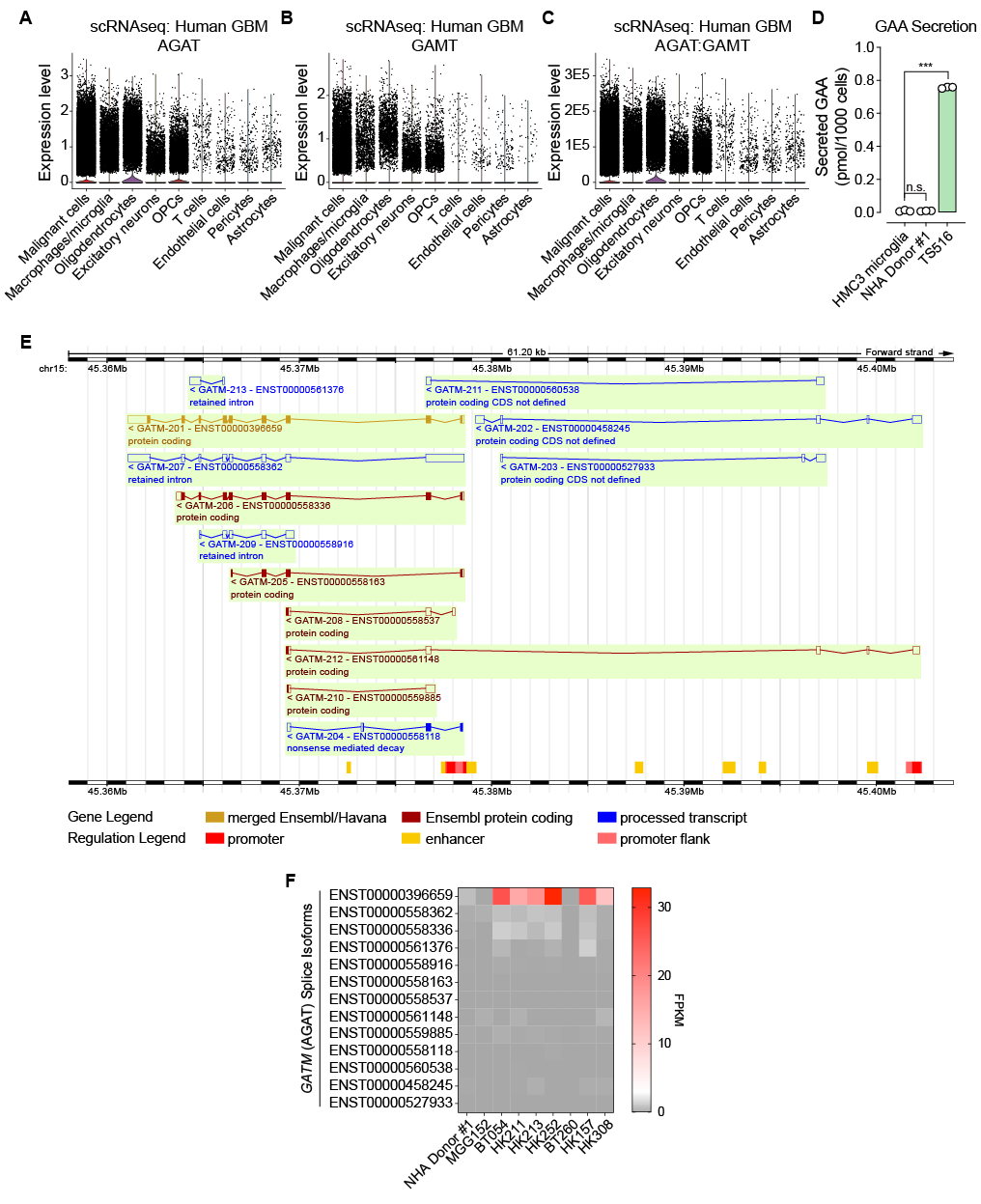

### Supplemental Figure 5

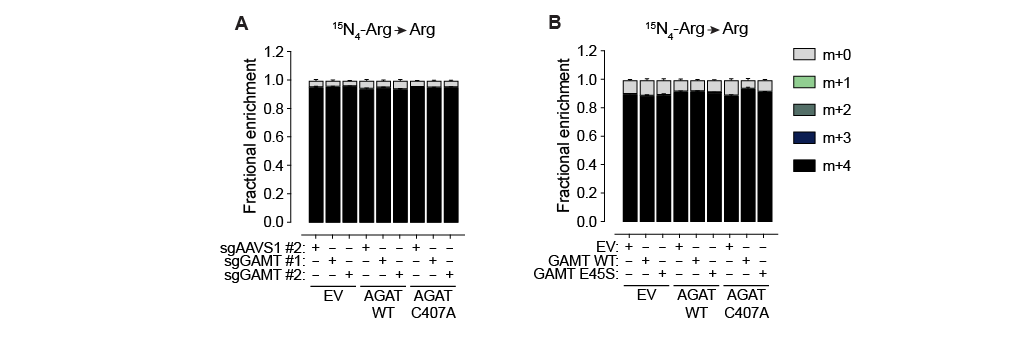

### Supplemental Figure 6

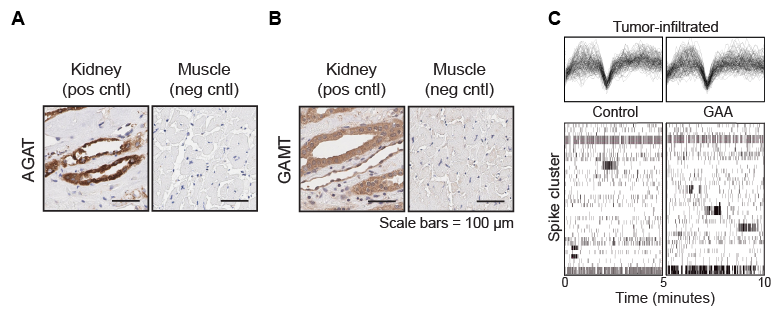

### Supplemental Figure 7

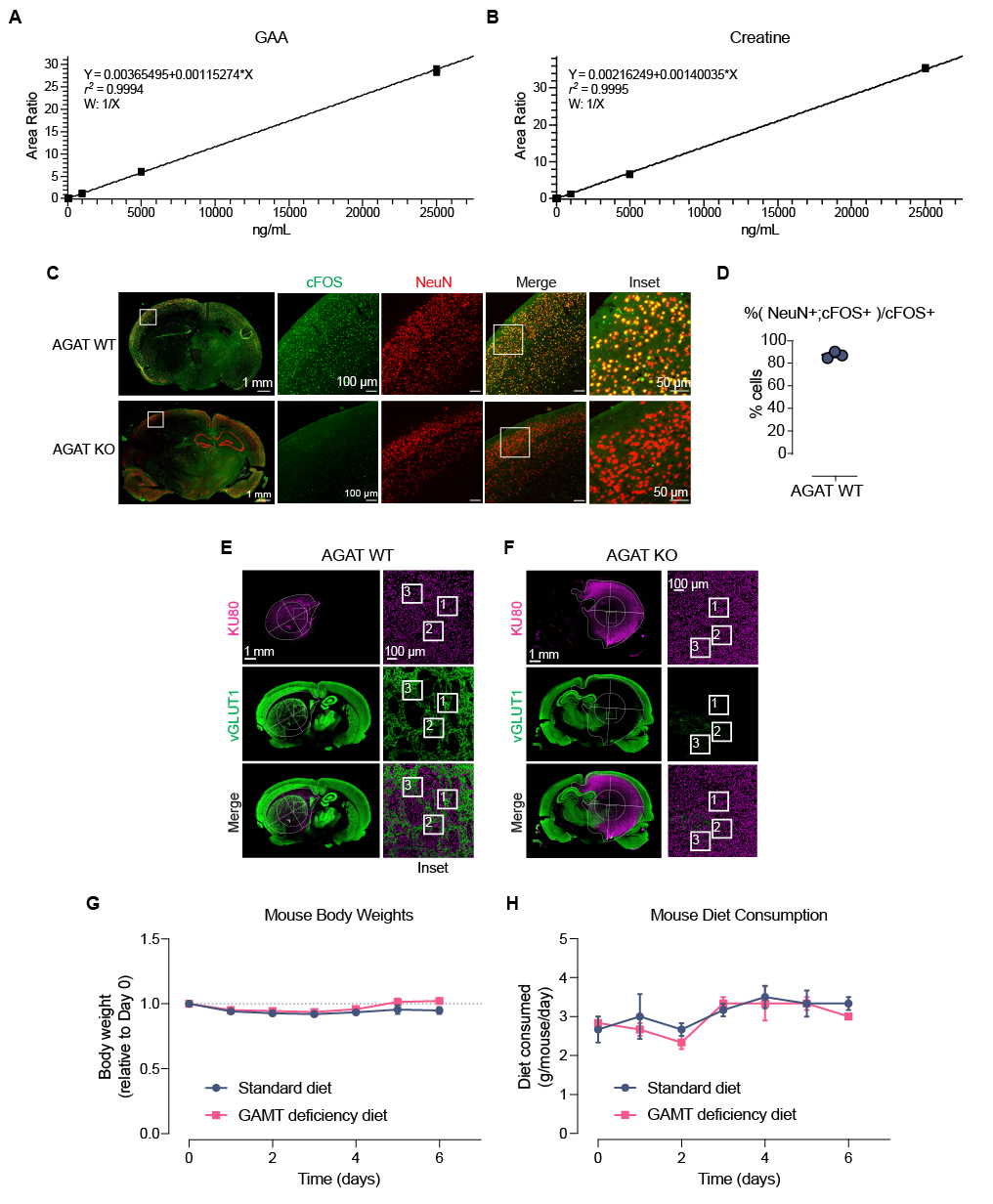
